# Benchmarking coarse-grained simulation methods for investigation of transport tunnels in enzymes

**DOI:** 10.1101/2024.09.16.613244

**Authors:** Nishita Mandal, Jan A. Stevens, Adolfo B. Poma, Bartlomiej Surpeta, Carlos Sequeiros-Borja, Aravind Selvaram Thirunavukarasu, Siewert J. Marrink, Jan Brezovsky

**Affiliations:** Laboratory of Biomolecular Interactions and Transport, Department of Gene Expression, Institute of Molecular Biology and Biotechnology, Faculty of Biology, Adam Mickiewicz University, Uniwersytetu Poznanskiego 6, 61-614 Poznan, Poland; International Institute of Molecular and Cell Biology in Warsaw, Ks Trojdena 4, 02-109 Warsaw, Poland; Molecular Dynamics Group, Groningen Biomolecular Sciences and Biotechnology Institute, University of Groningen, Nijenborgh 7, 9747 AG Groningen, the Netherlands; Biosystems and Soft Matter Division, Institute of Fundamental Technological Research, Polish Academy of Sciences, ul. Pawińskiego 5B, 02-106 Warsaw, Poland

## Abstract

Enzymes are pivotal to numerous biological processes, often featuring buried active sites linked to the surrounding solvent through intricate and dynamic tunnels. These tunnels are vital for facilitating substrate access, enabling product release, and regulating solvent exchange, which collectively influence enzymatic function and efficiency. Consequently, knowledge of tunnels is key for a holistic understanding of the effect of mutations as well as predicting drug residence times. Unfortunately, most transport tunnels are transient, i.e., equipped by molecular gates, rendering their opening a rare event that is often notoriously hard to study with conventional molecular dynamics simulations. To overcome the sampling limitation of such simulations, this study investigated the efficacy of three different coarse-grained (CG) molecular dynamics simulation methods for inferring enzyme tunnel structure and dynamics. Here, we covered the Martini and SIRAH models with different restraint protocols providing stability to CG proteins while to some extent biasing the sampling towards a reference structure. By contrasting CG results with all-atom simulations, we benchmarked the ability of CG methods to replicate ensemble characteristics of complex tunnel networks in haloalkane dehalogenase LinB and two of its mutants with engineered tunnel networks. The assessed tunnel parameters are essential for prioritizing functionally relevant tunnels and delineating the effect of mutations on transport tunnels. Our findings reveal that while CG methods significantly enhance the efficiency of tunnel analyses, some of them, like Martini with Elastic network restraints, were limited in recapitulating all-atom tunnel dynamics due to the structural bias applied. In contrast, the Martini Gō model even captured the intricate details of mutation perturbing tunnel dynamics. All studied CG methods performed well in capturing the geometry of tunnel ensembles in line with all-atom simulations. Additionally, the wider applicability of CG methods was verified by analyzing tunnel networks of nine enzymes from different combinations of structural and functional classes, demonstrating their potential to uncover new tunnel phenomena and validate their utility in broader biological and functional contexts. This comprehensive evaluation underscores the strengths and constraints of CG simulations in capturing enzyme tunnels and benefiting from their computational speed for studying huge datasets of enzymes. These insights are valuable for enzyme engineering, drug design, and understanding enzyme function while benefitting from the efficiency of coarse-grained models.

## Introduction

Enzymes are essential for life, enabling organisms to grow, maintain homeostasis, and reproduce. Despite extensive research, the intricate mechanisms underlying enzyme action remain only partially understood due to their immense complexity and diversity. Enzymatic catalysis occurs at the active site, which can be located on the enzyme’s surface or buried within a cavity^1,2^. This study focuses on enzyme tunnels that connect these buried active sites to the surrounding solvent^2–5^. Tunnels play critical roles in various processes, such as ligand selectivity at the active site, biomolecular exchange, and the fine-tuning of biochemical reactions^6^. To facilitate these functions, the tunnels are often equipped with one or more molecular gates^6,7^, the dynamical movement of which allows tunnels to adopt open and closed conformations, making them transient. To understand the structure-dynamics-function relationships of enzyme tunnels, it is crucial to consider tunnel-lining residues, bottleneck residues (the narrowest part of the tunnel), tunnel mouth residues (the initial point of contact with the substrate and solvent), and the dynamic nature of the tunnels. These tunnels, found in over 50% of enzymes across all six functionally distinct Enzyme Commission (EC) classes, are vital for regulating substrate entry, product release, and solvent movement, thereby influencing enzymatic reactions^1^. Their dynamics and geometry are key to understand enzyme activity and are particularly important in structure-based drug design, as mutations in tunnel residues can significantly alter enzyme properties^8^. Given their role in disease and potential as drug targets, studying tunnel dynamics is crucial in structural biology^7^.

Molecular dynamics (MD) methods are the preferred approach for studying these dynamic tunnels and observing conformational changes in residues^7,9^. Conventional MD (cMD) simulations, however, are computationally intensive and struggle to capture conformational changes that occur on longer timescales (milliseconds to seconds)^10^. Advanced enhanced sampling methods, such as metadynamics^11^ and umbrella sampling^12^, address some limitations of cMD simulations but remain computationally demanding and often require extensive system knowledge to set up collective variables (CVs), which are challenging to define for transient tunnels. In our recent work, we demonstrated that Gaussian accelerated MD^13^, which does not require CVs, can overcome the limitations of cMD simulations and successfully sampled rare tunnels in proteins. However, setting up the respective potential boosts is often system-specific and can be demanding, to some extent limiting the broad application of the approach to speed up the identification of transient tunnels.

Another approach to addressing the limitations of cMD simulations is by reducing the complexity of used all-atom (AA) definitions, grouping atoms or molecules into “beads” in coarse-grained MD (CG-MD)^14^, thereby achieving a coarser representation of molecular states. With fewer particles and interactions, CG-MD simulations are approximately 10–20 times faster than AA simulations, allowing for the exploration of longer timescales, from nanoseconds to milliseconds, with greater computational efficiency^14–16^. CG models can range from a single bead representing an entire protein (supra-CG resolution) to near-atomistic models, where multiple beads represent different chemical groups. In this study, we utilized two well-known CG models: SIRAH^17^ and Martini^18^.

The Martini CG force field is a well-established physics-based method. The new Martini 3.0 force field employs 2-4 atoms to a CG bead mapping scheme^19^, where bonded interactions are derived from AA simulations and nonbonded interactions are based on experimental free energy partitioning in chemical compounds^19^. With various particle types and subtypes, Martini accurately reflects the chemical nature of atomistic structures. Its residue-level coarse-graining aligns well with experimental methods^18^. Martini 3.0 features a dedicated water bead type, “W,” optimized separately from other targets to improve water properties, such as preventing freezing at room temperature, a problem in earlier models^19^. To maintain the stability of secondary and tertiary structures of proteins, Martini uses Elastic Network Models (Elastic)^20^ or Gō models (Gō)^21^. In this study, we employed both structural bias methods to stabilize the CG models. Elastic models, which connect pairs of backbone beads with harmonic bonds within a specified cutoff distance, serve as a structural scaffold but limit the observation of significant conformational changes^20^. In contrast, Gō models, which use Lennard-Jones (LJ) interactions instead of harmonic bonds, allow for the sampling of a more diverse conformational space and use the native structure’s contact map as the basis for pair interactions^22^. Martini is compatible with simulation engines such as GROMACS^23^ and OpenMM^24^. One of Martini’s major advantages is its expanding model library, which facilitates the creation and combination of various molecular classes. Martini 3.0 achieves a more realistic representation of protein cavities and ligand fitting through rebalanced cross-interactions and re-parameterization of bonded distances, enhancing the modeling of molecular volume and shape. This improvement allows for better predictions of binding thermodynamics and kinetics^19,25^. However, the coarse-grained nature of Martini can still lack the fine structural detail required for high chemical specificity, particularly in representing pocket-ligand interactions, differentiating enantiomers, and accurately modeling binding directionality^19^.

The CG topologies in SIRAH are obtained by fitting interactions to structural information and mapping actual atom positions to coarse-grained beads^17^. Protein backbone representative beads are placed at the real atom positions N, O, and Cα, allowing for simple equivalence with the all-atom (AA) dihedral angles. Generally, polar and aromatic components involved in hydrogen bonding or stacking interactions are depicted in greater detail, while hydrophobic or bulky components are represented more coarsely. SIRAH is available in popular simulation engines such as AMBER^26^, GROMACS, and NAMD^27^. Although SIRAH’s mapping technique complicates its generalization across all amino acids, it allows for greater resolution in CG methods. Additionally, SIRAH employs its own explicit CG solvent, composed of four beads interconnected in a tetrahedral pattern^28^. This solvent produces dielectric permittivity and permits variations in ionic strength in agreement with experimental setups. SIRAH utilizes a classical Hamiltonian to describe particle-particle interactions, common to most AA potentials. Since SIRAH operates unbiasedly, without constraints to maintain protein scaffolds, it can struggle to maintain secondary structures in larger proteins throughout the simulation^29^. The process of creating new molecules is limited by the complexity of generalizing the parameterization procedure, making it a more challenging task for users. Despite these limitations, SIRAH 2.0 demonstrates strong capabilities in capturing the overall qualitative behavior and compactness of protein-peptide interactions, making it a valuable tool for coarse-grained simulations. However, it has limitations in capturing specific conformational changes, particularly in complex binding events, as shown in the study of PMCA peptide recognition by CaM, a calcium-binding protein^17^.

Although both the SIRAH and Martini CG methods have been employed to investigate many different protein systems^16^, their suitability for simulating the spatiotemporal behavior of transport tunnels has yet to be evaluated. In this study, we address questions such as: How effective is CG-MD in capturing tunnel dynamics despite its coarse nature? How sensitive is CG-MD in capturing mutant tunnel exploration? And does the use of CG-biased models affect tunnel exploration?

We focused on haloalkane dehalogenases as a model system for enzymes with buried active sites to understand the nature of CG tunnels. Critical properties such as ligand transport, substrate inhibition, and cooperativity are often significantly affected by mutations in tunnels, highlighting their role in complex kinetic behavior in this enzyme family^30^. Although passage through transient tunnels is often gated by only a few residues, the degree of coarse-graining can significantly impact tunnel dynamics. Nevertheless, tunnel dynamics in mutants show different preferences compared to wild-type, and these parameters were used to assess the sensitivity of CG methods in capturing tunnel dynamics. We compared the tunnels captured through AA simulations and all three CG methods (Martini Gō, Martini Elastic, and SIRAH) to understand how effectively CG methods capture tunnels despite their coarse nature. LinB-Wt has the main tunnel p1, whereas p2, p3, and side tunnel (ST) are progressively more rare tunnels^13^. In mutants LinB-Closed, the p1 tunnel is less frequently open due to tunnel engineering which results in decreased catalytic efficiency of LinB-Closed, whereas in LinB-Open, the opening of p3 tunnel was engineered^31^, making LinB-Open mutant the most catalytically efficient variant till date^31^. We have compared the tunnel trends in LinB-Wt and mutant enzymes using different CG methods. To understand the effect of restraint bias on the CG tunnels, we compared the tunnel properties such as bottleneck radius, length, and curvature with AA tunnels. Additionally, we tested CG methods more systematically on tunnel networks from three different EC classes (EC 1-3) to capture the essence of tunnels in functionally distinct enzymes that can be studied using CG methods. To maintain also diversity in the structural context of explored tunnels, the selected enzymes comprise all-alpha, all-beta, and alpha/beta classes^1^. Such evaluation of tunnel occurrence and structural properties of their ensembles allowed us to validate the usability of CG methods in more general settings.

## Methods

### System setup and Conventional MD simulation system setup

All initial LinB variant structures: LinB-Wt (PDB code: 1MJ5), LinB-Closed (PDB code: 4WDQ), and LinB-Open (PDB ID: 5LKA) were protonated at pH 8.5^31^ with a salt concentration of 0.1 M using the H++ server^32^. For solvation, the OPC water model^33^ was used with neutralized counterions (Na+ and Cl-) to achieve a final concentration of 0.1 M. Energy minimization was carried out using 500 steps of steepest descent followed by 500 steps of conjugate gradient in 5 rounds with decreasing harmonic restraints, utilizing the PMEMD^34^ module of AMBER18^26^ with the ff14SB force field^35^. Restraints were applied as follows: 500 kcal/mol/Å^2^. on all heavy atoms, then 500, 125, 25, and 0 kcal/mol/Å^2^ on backbone atoms only. Then, the systems were equilibrated for 2 ns gradually heating to 310 K under constant volume using the Langevin thermostat^36^ with a collision frequency of 1.0 and harmonic restraints of 5.0 kcal/mol on all enzyme atoms. Periodic boundary conditions (PBC) and the particle mesh Ewald^37,38^ method were employed with electrostatics interactions beyond 10 Å cutoff. A 4 fs time step was used, enabled by the SHAKE algorithm^39^, and hydrogen mass repartitioning^40^. Subsequently, a 200 ns unrestrained NPT simulation was performed using PMEMD.CUDA with the Monte Carlo barostat^41^ and Berendsen thermostat^42^, storing frames every 20 ps. The thermostat choice was motivated by its fortuitous ability to preserve the intrinsic dynamics of the system and transport properties closely to NVE simulations, while still maintaining temperature control^43^. Clustering analysis was executed with the average-linkage hierarchical agglomerative algorithm using cpptraj^44^ on the 200 ns trajectory for each system. Requested number of clusters was set five while the secodnary cutoff of 4.5 Å was used to identify the five most diverse conformations. For each cluster, the centorid frame was selected as its representative. These representative conformations served as seed structures for an extended 5 μs unrestrained NPT simulation under, with frames stored every 200 ps, for all three LinB variants.

The same simulation protocol was applied across nine enzymes, EC1 (PDB IDs: 1jfb, 1gp4, and 3bur), EC2 (PDB IDs: 1m15, 1oyg, and 1q20), and EC3 (PDB IDs: 2oup, 1dim, and 1cvl) to capture structural and functional diversity of enzymes with buried active sites. Due to the progression in the project timeline, updated AMBER22 software was used for performing these simulations. Single unrestrained NPT simulation was carried out for 2 μs for each of these enzymes with frames stored every 200 ps. The co-factors FE^2+^ ion and CA^2+^ ions were restrained using distance restraints within the enzyme. No restraint was used for HEME and NADP co-factor.

### System setup and Coarse-Grained MD simulations using SIRAH

To set up SIRAH CG models, first, the five most diverse structures obtained using cluster analysis in AA simulations were used as starting seed structures. The atomic seed structures were then mapped to the SIRAH CG model using SIRAH Tools^45^ and SIRAH 2.0 force field. For solvation, pre-equilibrated CG WT4 water molecules were used. The WT4 molecules were randomly replaced by Na+ and Cl– CG ions to achieve ionic strength of 0.1 M, and counterions were added to the system according to Machado et al^46^. System minimization was performed using 5000 steps of steepest descent followed by 5000 steps of the conjugate gradient in 5 rounds with decreasing harmonic restraints, utilizing the PMEMD.CUDA module of AMBER18 with the ff14SB force field. Restraint of 500 Kcal/ mol/Å^2^ was applied on heavy atoms, followed by 500, 125, 25, and 0 kcal/mol/Å^2^ on backbone GN and GO beads. Alongside in-house restraint protocol was applied to dihedral angles of residues in α-helices (4 kcal/mol/ Å^2^) and distance between H-bond forming atoms of residues in β-sheets (20 kcal /mol/ Å^2^) to keep the structure stable (Figure S1). These restrained residues were selected using secondary structure analysis using cpptraj module secstruct on respective input structures. The structures were equilibrated for 2 ns gradually heating it to 310 K followed by 5 ns NVT and 10 ns NPT equilibration. PBC and the particle mesh Ewald method were employed with electrostatics interactions beyond 10 Å cutoff. Finally, 20 fs time-step was used to produce 5 μs of unrestrained production simulation at constant pressure and temperature using a Langevin thermostat at collision frequency 50 ps^-1^ and Berendsen barostat. The choice of parameters was adopted from SIRAH 2.0 manuscript^17^.

Simulation protocol was kept the same for seven enzymes, EC1 (PDB ID: 1gp4), EC2 (PDB IDs: 1m15, 1oyg, and 1q20) and EC3 (PDB IDs: 2oup, 1dim, and 1cvl). Updated AMBER22 software was used for performing these simulations. Single unrestrained NPT simulation was carried out for 2 μs for each of these enzymes with frames stored every 200 ps. Due to the unavailability of small-molecule library in SIRAH, co-factors such as HEME and NADP were absent, thus the enzymes 1jfb and 3bur were not simulated for SIRAH and they are excluded from the analysis. Additionally, the cofactors FE^2+^ ion (replaced by CA^2+^ ions, due to the unavailability of FE^2+^ ion parameters in SIRAH)^47^ and CA^2+^ ions were restrained using distance restraints within the enzyme.

### System setup and Coarse-Grained MD simulation using Martini

For CG Martini system setup, the seed structures were kept the same as obtained from the clustering analysis of AA simulations. All the calculations were performed with GROMACS 2023.3 version. To keep the native structure of the CG stable, Elastic network and GōMartini were applied, keeping the default settings. The bonded parameters of Elastic network and GōMartini were kept the same, to compare the structural propensity. Martini force field version 3.0^19^ was used. For GōMartini model, first, the contact map was obtained from the web server Gō ContactMap^48,49^, followed by conversion of AA structure to CG model using martinize2 tool^50^. For Gō contact map, the distance cutoff between CA-CA was kept in the range of 0.3-1.1 nm. The strength of the Gō potentials was kept at the default value of 9.414 kJ/mol. Then secondary structures were assigned based on AA structure with DSSP module^51^ and martinize2 tool was used to switch on Elastic network with bond force constant 700 kJ.mol^-1^.nm^-2^, and the lower and upper elastic bond cut-offs were kept at their default values of 0.5 and 0.9 nm, respectively. Cutoff values from 0.8 to 1.0 nm were shown to provide sufficient agreement with the AA simulations^20^.

The simulation protocol was kept the same for Elastic and Gō models. Energy minimization was carried out using 400000 steps of steepest descent. Then, the systems were NVT equilibrated position restrained for 20 ns gradually heating to 310 K under constant volume using the V-rescale thermostat^52^ with 10 fs time step. PBC was kept default to xyz, applying full PBC in all three dimensions and Non-bonded cutoff, rlist was set to 1.4 with cutoff scheme Verlet. The electrostatic interaction (rcoulumb) and van der Waals interactions (rvdw) cutoffs were both set to 1.2 nm. NVT was followed by position-restrained NPT equilibration for 20 ns with 10 fs time step with pressure control using the C-rescale barostat^53^. Finally, 5 μs unrestrained production simulation was run using 20 fs time-step with Parrinello–Rahman barostat^54^ and V-rescale thermostat.

The same simulation protocol was applied across nine enzymes, EC1 (PDB IDs: 1jfb, 1gp4, and 3bur), EC2 (PDB IDs: 1m15, 1oyg, and 1q20), and EC3 (PDB IDs: 2oup, 1dim, and 1cvl) to capture the diversity of enzyme classes. Single unrestrained NPT simulation was carried out for 2 μs for each enzyme with frames stored every 200 ps. The CG representation of HEME and NADP were adopted with Martini 2.0 beads^55^, which were updated for Martini 3.0 beads and used for simulation. Due to the larger time-step, we applied positional restraints involving HEME (1jfb) and NADP (3bur) to keep the co-factor in place which is necessary for enzyme function. Additionally, the cofactors FE^2+^ ion (replaced by CA^2+^ ions which is represented by S type bead^47^, due to the unavailability of FE^2+^ ion) and CA^2+^ ions were restrained using distance restraints within the enzyme.

### Protein stability analysis

To confirm the stability of enzymes explored through AA and CG-MD, Root Mean Square Deviation (RMSD), and Root Mean Square Fluctuation (RMSF) were calculated with the reference as initial structure, taking into account all the heavy atoms of the protein. RMSD was calculated on all protein residues. The radius of gyration (Rg) and Solvent accessible surface area (SASA) were calculated to confirm if the protein was intact throughout the simulation. These analyses were performed with, cpptraj and gmx^23^ tools for AMBER and GROMACS simulations, respectively.

### Tunnel ensemble analysis

Tunnels were analyzed through CAVER 3.2.0 software^56^. Amino acid library was updated to respective CG bead and wan-der Waal’s radii to calculate CG-Tunnels for SIRAH and Martini. Starting point for tunnel calculation defined by three catalytic residues (N38, D109, and H272; numbering corresponds to the crystal structure) and a probe radius of 0.9 Å were used to explore potential pathways in enzymes using divide-and-conquer approach^57^ which uses TransportTools software^58^ internally to unify tunnels from caver clusters, to run CAVER 3.2.0 in 10 batches using frame clustering cutoff 3.0 and frame reweighting coefficient 2.0. For AA simulations, the filtering caver cutoff was set to 5 in 1000 frames to preserve rare tunnels and for CG methods it was set to 250 out of 1000 frames due to the high frequency of tunnel detection. Later reclustering algorithm^59^ was used to ensure only the presence of tunnels with consistent entrances in any cluster using HDBSCAN on the tunnel endpoints, with the following parameters: cluster_selection_epsilon of 1.5, allow_single_cluster set to True, and min_samples and min_cluster_size set to 5. The reclustered tunnels were considered as independent tunnel clusters, while the tunnels identified as noise were discarded. Subsequently, we used comparative analysis from TransportTools software v0.9.3 with the minimum number of simulations set to 5 to generate a unified tunnel network from CG and AA simulations. The reference structures were obtained using backmapping protocol for Martini using backwards.py script^60^. Similarly, for SIRAH the reference structures were obtained using the backmapping module of SIRAH tools^45^. The method for clustering the final results was set to average-linkage, with a clustering cutoff of 1.5. The same procedure was followed for enzymes from the respective three EC classes, separate TransportTools comparative analysis was carried out for each protein variant.

### T-test on tunnel properties

Two sample T-tests were performed on tunnel properties such as bottleneck radius, occurrence and length from AA and CG-MD simulations using Python SciPy^61^ library to determine the similarity between the outcomes of the benchmarked methods. The T-test was conducted to determine the statistical significance of the differences between the means of the two datasets. A two-tailed P-value was calculated for each comparison. A P-value below a conventional threshold (i.e., 0.05) is considered a statistically significant difference.

## Results and Discussions

### CG models of three LinB variants remain stable during simulations

The first step to analyze any CG-MD was to ensure the stability of the system throughout the simulation. Despite the use of restraints in Martini CG methods, it was important to check the compactness of the protein during the simulation. For the default SIRAH CG model, as there was no restraint protocol available, it led to the loss of secondary structures (Figures S2-S4). To address this issue, we developed an in-house restraint protocol by applying hydrogen bond restraints on β-sheets and dihedral angle restraints on α-helices (Figure S1). By applying these restraints, we observed a marked improvement in structural stability (Figures S2-S7). Specifically, the percentage of coil content reduced by approximately 10%, and alpha-helix and beta-sheet structures remained stable throughout the simulation.

To confirm the global stability of proteins in our restraint CG-MD simulations, we performed an RMSD analysis of the LinB-Wt protein with all residues involved and contrasted the data with outcomes from AA simulations (Figure 2A). We observed that the Martini Elastic model was closest to the AA simulations, followed by the Gō and SIRAH, in line with the use of much more extensive restraints in the Elastic network (Table S1). Similar observations were seen for LinB-Closed and LinB-Open mutants (Figures S8,S9) On the other hand, because of the more rigid model in Elastic, it results in a more restrained dynamic movement of the protein. The limited protein flexibility with the Elastic model from among CG methods is clearly visible from RMSF data (Figures S10-S12). In contrast, the Gō and SIRAH models allowed more pronounced protein dynamics (Figure 2A). We also confirmed that the functionally important conformation of catalytic pentad (Figure 2B) is not overly disturbed by CG methods through the simulations (Figure S13), the pentad was the most preserved with Elastic and the least with SIRAH model. Additionally, we analyzed Rg and SASA for respective simulations. For Rg, we observed stable behavior of all simulations from LinB-Wt, LinB-Open, and LinB-Closed, with Elastic simulations producing more compact structure, while some SIRAH simulations exhibited an increase of up to 2 Å compared to AA ones (Figure S14-S16). Similarly, SASA values were stable for all simulations irrespective of the model used (Figure S17-S19). However, SASA with SIRAH model was approximately twice larger than AA for all LinB variants, indicating a larger body of the protein, in line with the elevated Rg, and likely a higher porosity as well. Overall, those analyses verified the stability of both AA and CG-MD simulations and their suitability for the follow-up tunnel analyses (Figure S8-S19).

**Figure 1.**
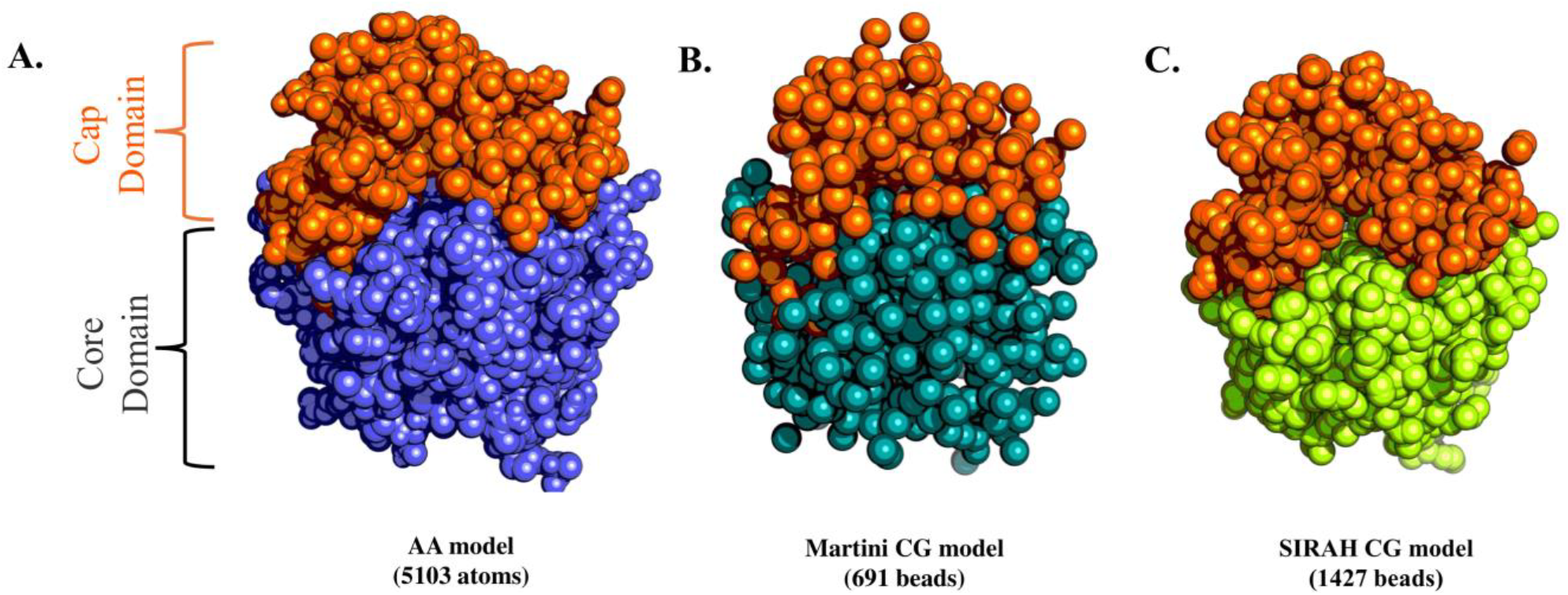
Representations of level-of-granularity in LinB-Wt enzyme: A) in AA model, B) in Martini CG model and C) in SIRAH CG model.

**Figure 2.**
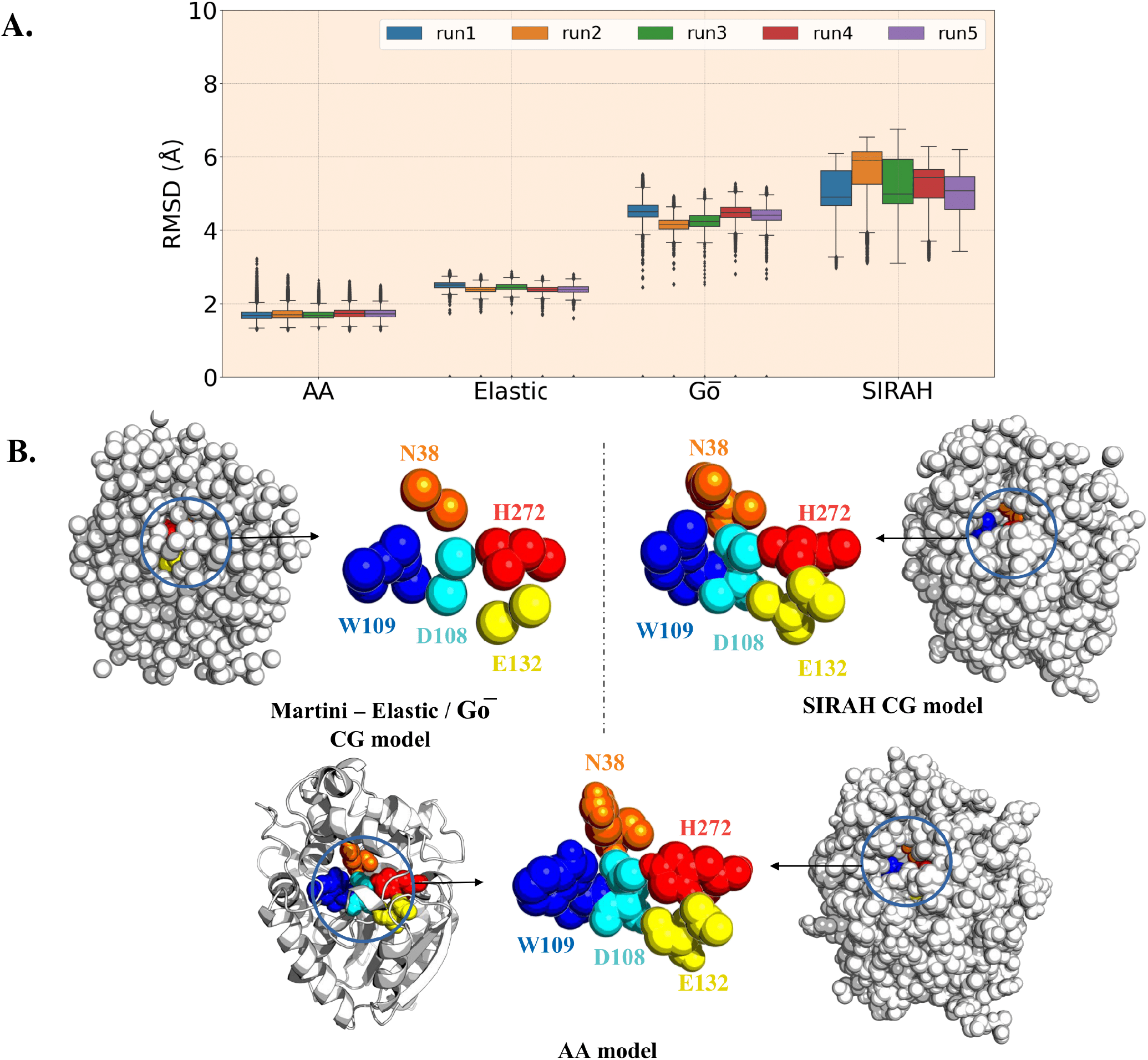
Stability of LinB-Wt and representation of its overall structure and catalytic machinery in the investigated models. **A)** RMSD of whole LinB-Wt protein with all residues in five replicates using AA and CG methods – SIRAH, Elastic, and Gō. The box plot shows the median (middle line in the box) and the box represents the interquartile range, which ranges from the 25^th^ percentile to the 75^th^ percentile and gives a sense of how spread out the middle 50% data is. The whiskers extending from the box represent a range of data. **B)** Representation of LinB-Wt structure with Martini, SIRAH and AA models, with the residues of the catalytic pentad in the active site, showing different granularity levels between the used models.

### Exploration of tunnels using CG models in comparison to AA model

After making sure that all CG simulations were stable, the global comparison between all the CG-MD and AA tunnels was performed using TransportTools comparative analysis. From the result, we can see that all the CG methods were able to capture the main and transient tunnels in LinB-Wt (Figure 3A) and additionally, many new tunnels apart from ones captured through AA simulations were identified (Figure 3B), especially in the more porous SIRAH model. We observed that structural properties of tunnel ensembles obtained with CG-MD, such as bottleneck radius and tunnel length, were similar to AA ones, with the exception of a longer p3 tunnel and shorter ST captured through SIRAH in LinB-Wt (Figures 3A,3C and Table S2). Similar tendencies for the length of ST and p3 tunnels were also observed with both two mutants (Figures S20-S21, Tables S3,S4). The largely prevalent similarity between the structure of tunnel ensembles produced by CG and AA tunnels confirms that CG tunnels can be effectively used to infer their geometry accurately enough to enable further analyses of these tunnels with more advanced computational approaches to assess their role in substrate and product transport explicitely^62^.

**Figure 3.**
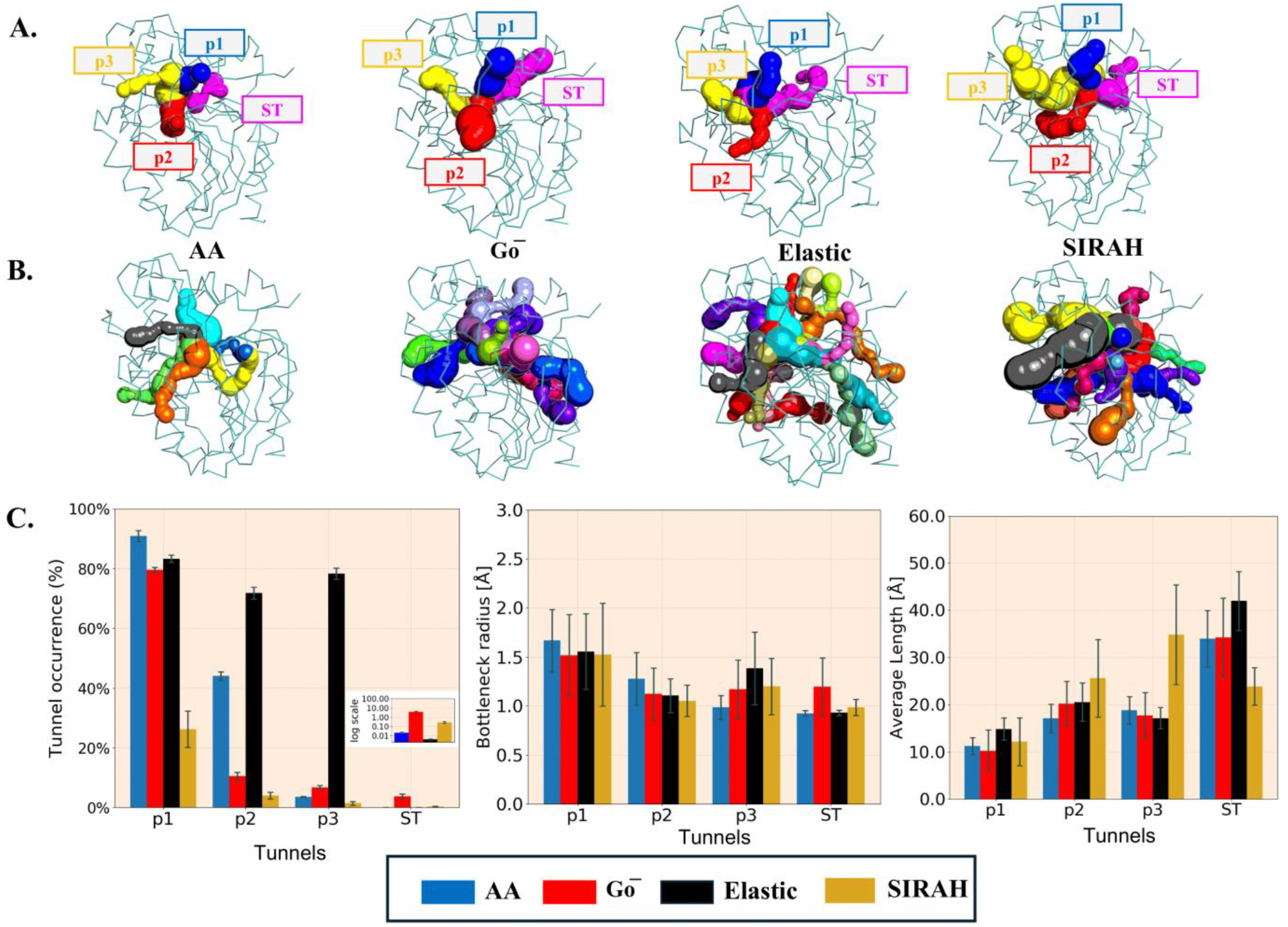
Comparison of tunnel structure and occurrence in LinB-Wt variant. **A)** Representation of all main tunnels obtained through individual methods. **B**) shows additional tunnels captured by each method (cutoff applied tunnels seen in at least 10% of total simulation time and at least three simulations). **C)** Properties of main tunnels captured using different methods. The inset shows tunnel occurrence of the rare ST on the log scale. Data represents average±SD obtained from 5 simulation replicas. See Table S2 for details on statistical significance.

Considering the tunnel dynamics, the rates with which the individual tunnels occur in CG-MD simulations were mostly significantly different from the ones inferred from AA simulations (Figure 3C and Table S2), most likely due to the use of the extra restraints that alter the intrinsic dynamics of the investigated proteins. Analysis of tunnel occurrence is frequently used as a proxy to estimate the functional relevance of tunnels, particularly their priority^63^. Starting from the global tunnel ranking, all methods were able to distinguish the primary p1 tunnel (often considered close to being permanently open in LinB-Wt^31^) from the remaining transient tunnels, as well as maintain the priority of transient tunnels from most common p2, via rare p3, to very rare ST. The only exception was the Elastic model that markedly overestimated the relevance of p3 tunnel, which is equipped with two molecular gates formed from bulky side chains. This unbalanced prioritization led to poor separation of transient and permanent tunnels in simulations with this model. This observation aligns with the well-known drawback of Elastic models, i.e., over-stabilizing the initial conformation of protein backbone, which can introduce inaccuracies in protein-protein interactions and cause proteins to appear too “rigid” limiting their conformational flexibility^64–66^. Interestingly, the overstabilization of open states of p3 tunnel is maintained in the LinB-Closed mutant featuring the exact same tunnel (Figure S20) but is no longer present in LinB-Open (Figure S21), in which the residues of molecular gates of p3 tunnel were substituted with less bulky side-chains^31^. For the same reasons, the Elastic model resulted in the reduced occurrence of ST, the opening of which requires considerable disruption in one of the Cap Domain helices of LinB enzyme^13^. SIRAH model delivered appreciable separation between the priority of the main p1 tunnel and the other transient ones, with the caveat of the most underrepresented p1 tunnel in LinB-Wt (Figure 3C). However, this trend with the SIRAH model was not maintained in either of the two LinB variants with partially blocked p1 tunnel by L177W mutation (Figures S20-S21). Finally, the Gō model performed the best in recapitulating the AA findings, i.e., there was a clear separation between “permanent” and transient tunnels, and the priority of the tunnels was analogous to AA simulations (Figure 3C).

### How sensitive CG-MD simulations are for the assessment of tunnel relevance?

Despite differences in tunnel occurrences, we wanted to check if CG methods are sensitive enough to account for changes in tunnel dynamics due to mutations. For this purpose, we compared p1 and p3 tunnels of LinB-Wt with their engineered counterparts in LinB-Open and LinB-Closed variants^13,31^. The p1 tunnel that serves as the primary transport route, occurs less frequently in both variants due to the mutation at the p1 tunnel mouth (L177W). In contrast, the LinB-Open variant had an engineered p3 tunnel enhancing its functionality due to additional mutations (W140A+F143L+I211L). These changes were specifically driven by modifications in bottleneck residues, aligning the p3 tunnel’s properties with the natively present p2 tunnel. The expected effects of mutations were well reproduced by AA simulations as well as those Gō model employing (Figure 4). Conversely, the Elastic model could recapitulate only the p1 closure trends but lacked the opening effect in p3 tunnel, due to the overestimated occurrence of p3 in LinB-Wt and LinB-Closed. Finally, the SIRAH model failed to capture any of these trends, potentially due to the limitations of the coarse-grained representation, in particular, the application of our local secondary-structure-based restraints, making it less effective in recapitulating the tunnel dynamics than the other CG models.

**Figure 4.**
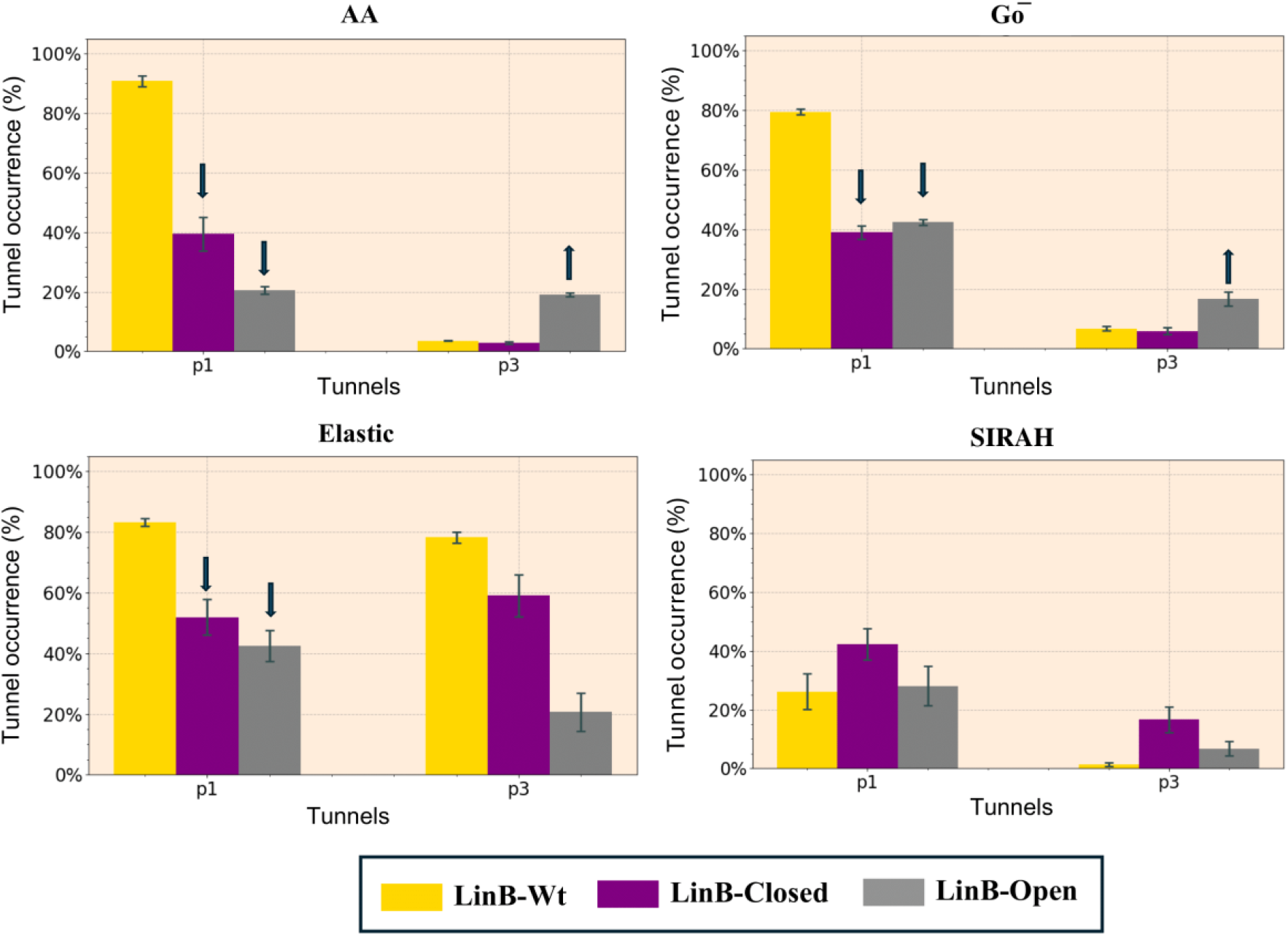
Comparison of the effect of mutations on tunnel occurrence delineated by different methods. Tunnel p1 and p3 from AA, Gō model, Elastic model, and SIRAH model in LinB-Wt, LinB-Closed, and LinB-Open. Data represents average±SD obtained from 5 simulation replicas

### Case study: Diverse enzymes from three distinct functional and structural classes

To evaluate the broader applicability of CG methods, we performed MD simulations and analyzed tunnels in nine enzymes from three functional EC classes (EC1, EC2, and EC3), one enzyme from the following distinct structural classes: all-alpha, all-beta, and alpha/beta. To evaluate the behavior with a diverse range of enzymes with single buried active sites and their tunnel networks. Overall, the stability and dynamics of all proteins simulated with all methods followed the same trends as observed for LinB enzymes (Figures S23-S34).

EC1 class of enzymes represents oxidoreductases, catalyzing redox reactions^67^. These reactions involve the transfer of electrons from a donor to an acceptor. EC1 enzymes often use cofactors such as NAD^+^/NADH or FAD/FADH_2_ to facilitate the electron transfer. These enzymes are vital in processes like detoxification or respiration, where they play key roles in energy production by transferring electrons during metabolic pathways. Here, we studied three enzymes with buried active sites from this class – nitric oxide reductase with HEME cofactor (PDB ID: 1jfb; all-alpha), anthocyanidin synthase (PDB ID: 1gp4; all-beta), and delta(4)-3-ketosteroid 5-beta-reductase with NADP cofactor (PDB ID: 3bur; alpha/beta). We could not perform simulations of two of these proteins (1fjb and 3bur) with the SIRAH model due to the lack of cofactor parameters in the force field. Hence observations for this CG method are limited to a single all-beta protein in this EC class. CG-MD simulations identified equivalent tunnels for the majority of those observed in AA simulations, in particular when considering the most prevalent tunnels (Figure 5A). However, all CG models primarily exhibited higher tunnel occurrences for the analogous tunnels compared to the AA model as well as a clear tendency to identify many additional tunnels with primarily lower occurrences, whereas the conversed cases of AA tunnel not being covered by CG tunnels were much less prevalent (Figures S35-S37). To probe into the quantitative correspondence of key structural and dynamical properties of CG and AA tunnels, we have performed a correlation analysis of their occurrences, bottleneck radii, and lengths (Figure 5B, Figures S38-S40). Similarly to the LinB variants, studied enzymes from the EC1 class exhibited a rather low correlation between CG and AA tunnel occurrences. Interestingly, considerably larger correlations were observed among occurrences inferred by the CG methods alone (Figure 5B). In contrast to the tunnel dynamics, the structural properties of tunnel ensembles from AA and CG simulations were in much better agreement (Figure 5B), analogously to the situation with LinB variants.

**Figure 5.**
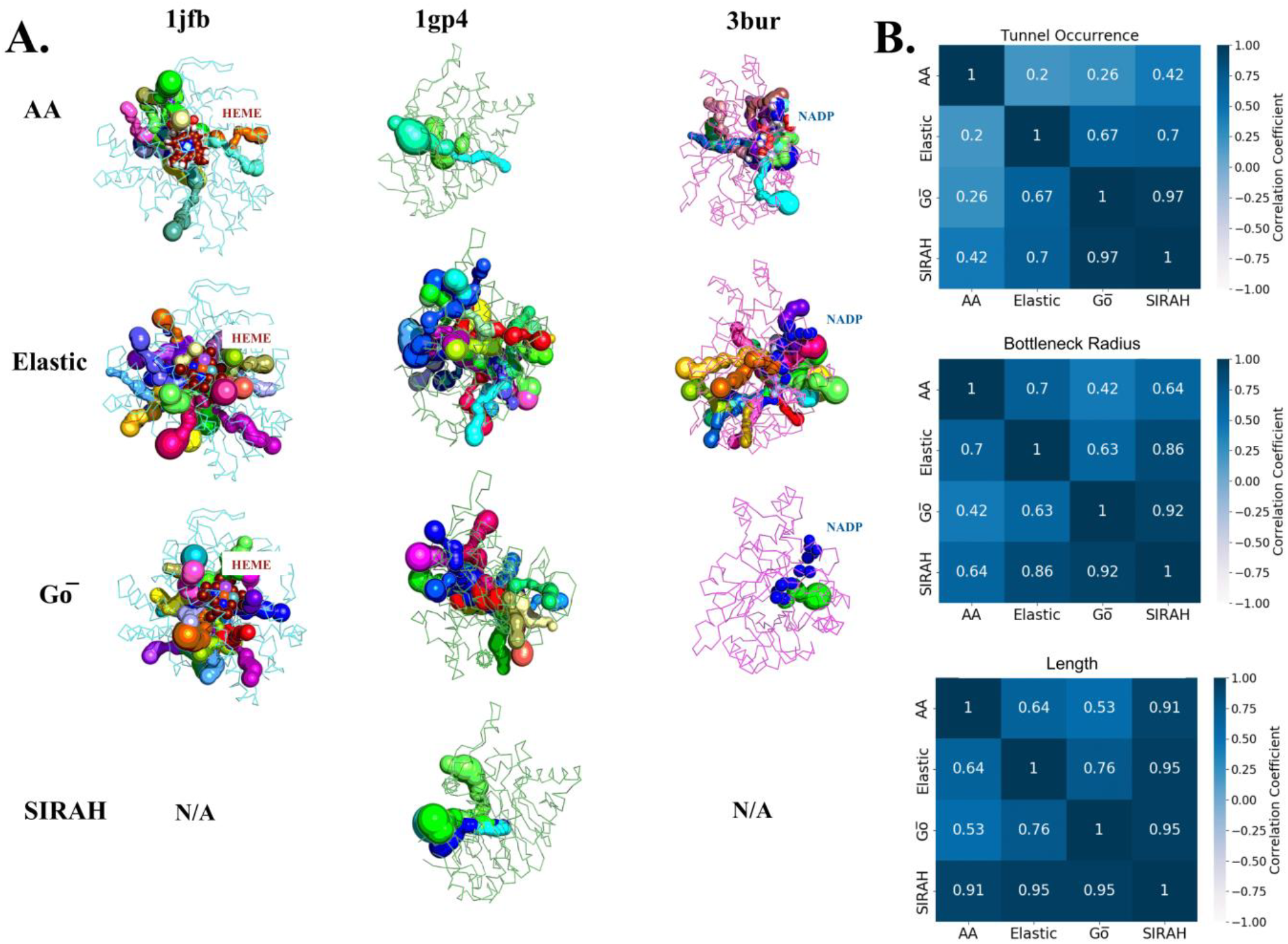
Correspondence of tunnels present in enzymes from EC1 class and their properties between AA and CG models. **A**) Enzymes tunnels with at least 5% and 10% occurrence in AA and CG models, respectively. The tunnels shown are representative of the average length and bottleneck radii for each cluster. Tunnel ensembles from SIRAH simulations of 1jfb and 3bur are not available due to the lack of parameters for the cofactors. **B**) Heatmaps with Pearson correlation coefficients between properties of tunnels present simultaneously in AA and each CG method. Data on the statistical significance of the observed correlations are available in Table S5.

EC2 class of enzyme represents transferases that catalyze the transfer of functional groups from one molecule to another^67^. They are essential for biosynthesis and metabolic regulation and are involved in processes like phosphorylation and glycosylation. Here we studied three enzymes with buried active sites specific to this class – arginine kinase (PDB ID: 1m15; all-alpha), Bacillus subtilis levansucrase (PDB ID: 1oyg; all-beta), and human cholesterol sulfotransferase SULT2B1b (PDB ID: 1q20; alpha/beta). For this class of enzymes, we have again systematically observed many additional CG tunnels of lower occurrences (Figure 6A and Figures S35-S37). Unlike enzymes from the EC1 class, the studied enzymes of the EC2 class exhibited a moderate to high correlation between AA and CG models in both structural and dynamic properties (Figure 6B and Figures S41-S43).

**Figure 6.**
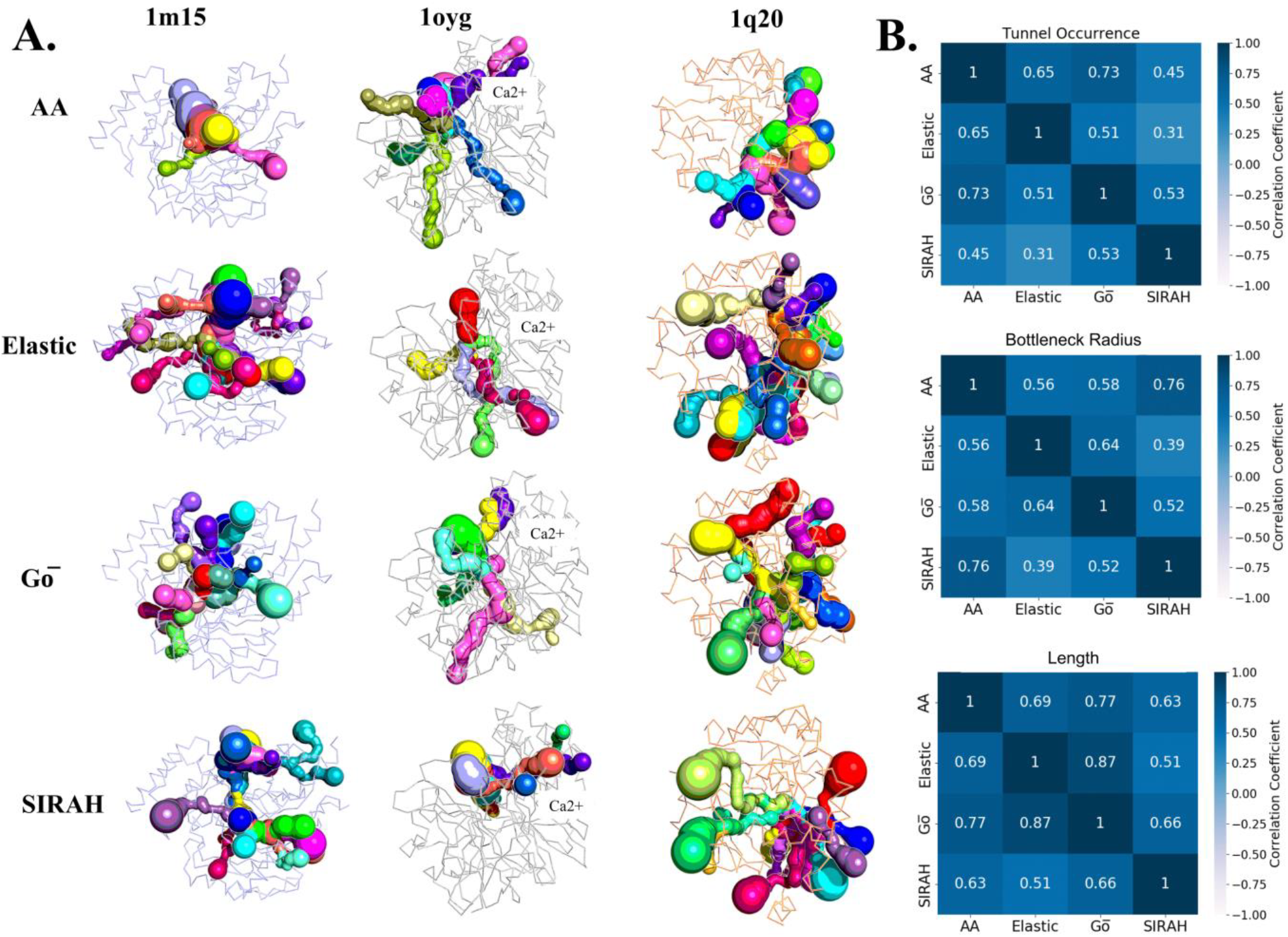
Correspondence of tunnels present in enzymes from EC2 class and their properties between AA and CG models. **A)** Enzymes tunnels with at least 5% and 10% occurrence in AA and CG models, respectively. The tunnels shown are representative of the average length and bottleneck radii for each cluster. **B**) Heatmaps with Pearson correlation coefficients between properties of tunnels present simultaneously in AA and each CG method. Data on the statistical significance of the observed correlations are available in Table S6.

At last, we evaluated the CG methods with the EC3 class of enzymes, which belong to hydrolases, catalyzing hydrolytic breakage of various bonds such as peptide or ester bonds. These enzymes are crucial in digestive processes, breaking down macromolecules like proteins, fats, and nucleic acids into smaller components, aiding in their metabolism and recycling. Following three enzymes with buried active sites from this class were analyzed – phosphodiesterase 10 (PDB ID: 2oup; all-alpha), Salmonella typhimurium LT2 neuraminidase (PDB ID: 1dim; all-beta), and bacterial lipase (PDB ID: 1cvl; alpha/beta). Again, additional tunnels and tunnel branches not present in the AA model were captured through CG methods, mainly concerning tunnels with lesser priority than the ones mutually identified by both CG and AA methods (Figure 7A and Figures S35-S37). Interestingly, for the 1dim enzyme, we have identified only one tunnel cluster in AA, possibly due to its rather shallow active site. For all three enzymes from this EC class, we observed the strongest positive correlation in the structural properties of tunnels (bottleneck radius and length) between CG and AA models yet (Figure 7B and Figures S44-S46), while dynamics governed tunnel occurrences maintained at least moderate agreement with AA.

**Figure 7.**
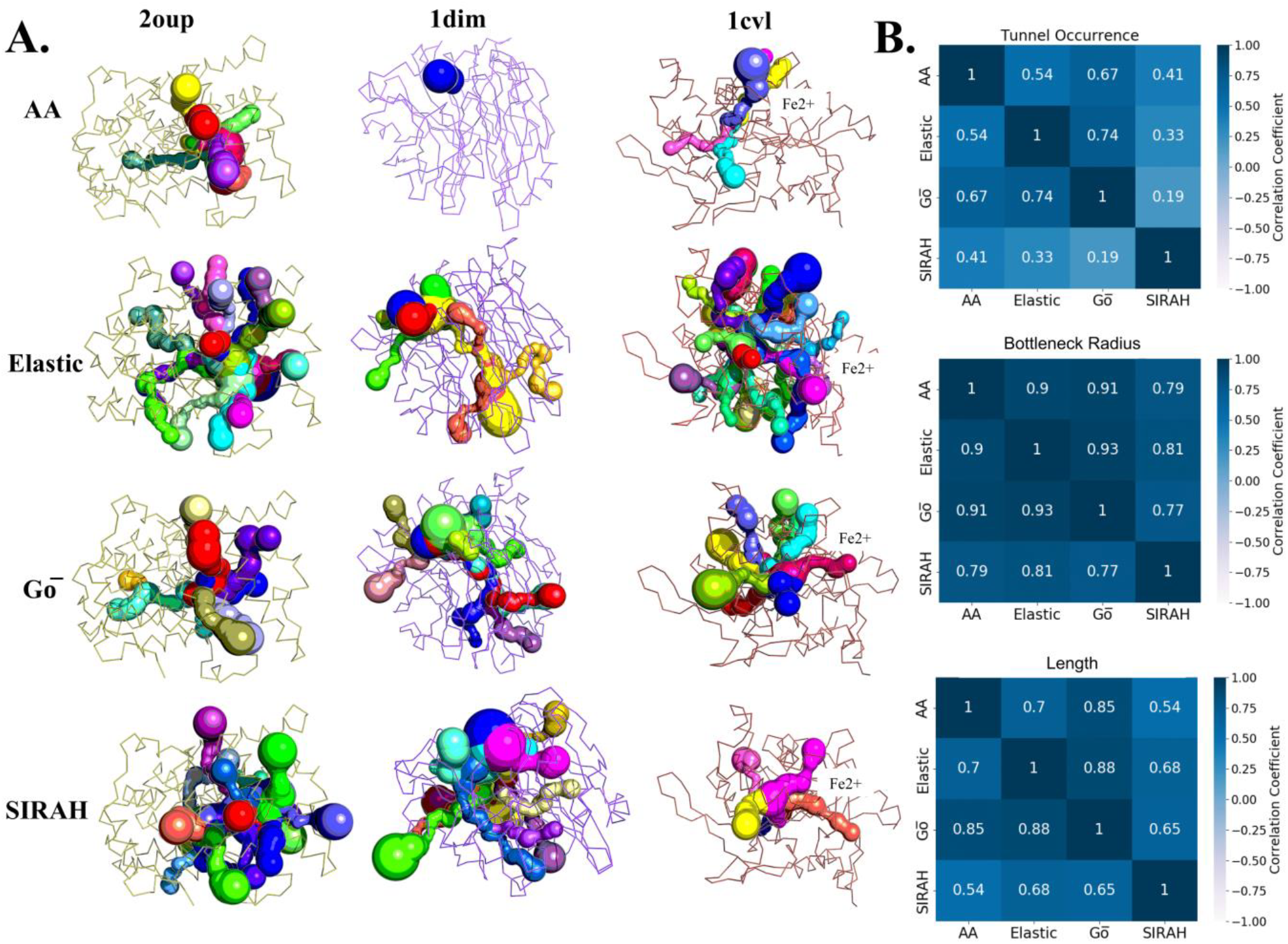
Correspondence of tunnels present in enzymes from EC3 class and their properties between AA and CG models. **A)** Enzymes tunnels with at least 5% and 10% occurrence in AA and CG models, respectively. The tunnels shown are representative of the average length and bottleneck radii for each cluster. **B**) Heatmaps with Pearson correlation coefficients between properties of tunnels present simultaneously in AA and each CG method. Data on the statistical significance of the observed correlations are available in Table S7.

In summary, the CG methods were able to capture similar tunnels to those originating from AA simulations, in particular considering the functionally most important ones with the highest occurrences (Figure 8A). Due to improved sampling and lower resolution, CG methods often detected additional, predominantly rare tunnels while missing only some low-frequency AA tunnels (Figure 8A). Despite the diverse nature of the enzymes in the dataset, covering tunnels of very different lengths, bottleneck radii, and rates of occurrences (Figures S38-S46), CG methods reached considerable agreement with the insights from AA simulations (Figure 8B). The best recapitulation of enzyme dynamics governing tunnel occurrences within our use case was reached with Gō model. Considering the geometries of produced tunnel ensembles, Elastic and SIRAH methods exhibited the highest correlation with bottleneck radii of AA tunnels. However, simulations with SIRAH were more prone to overestimate tunnel lengths, exhibiting the lowest correlation with AA data. Overall, these trends in the performance of individual methods corroborate the conclusions from analyses of LinB variants.

**Figure 8.**
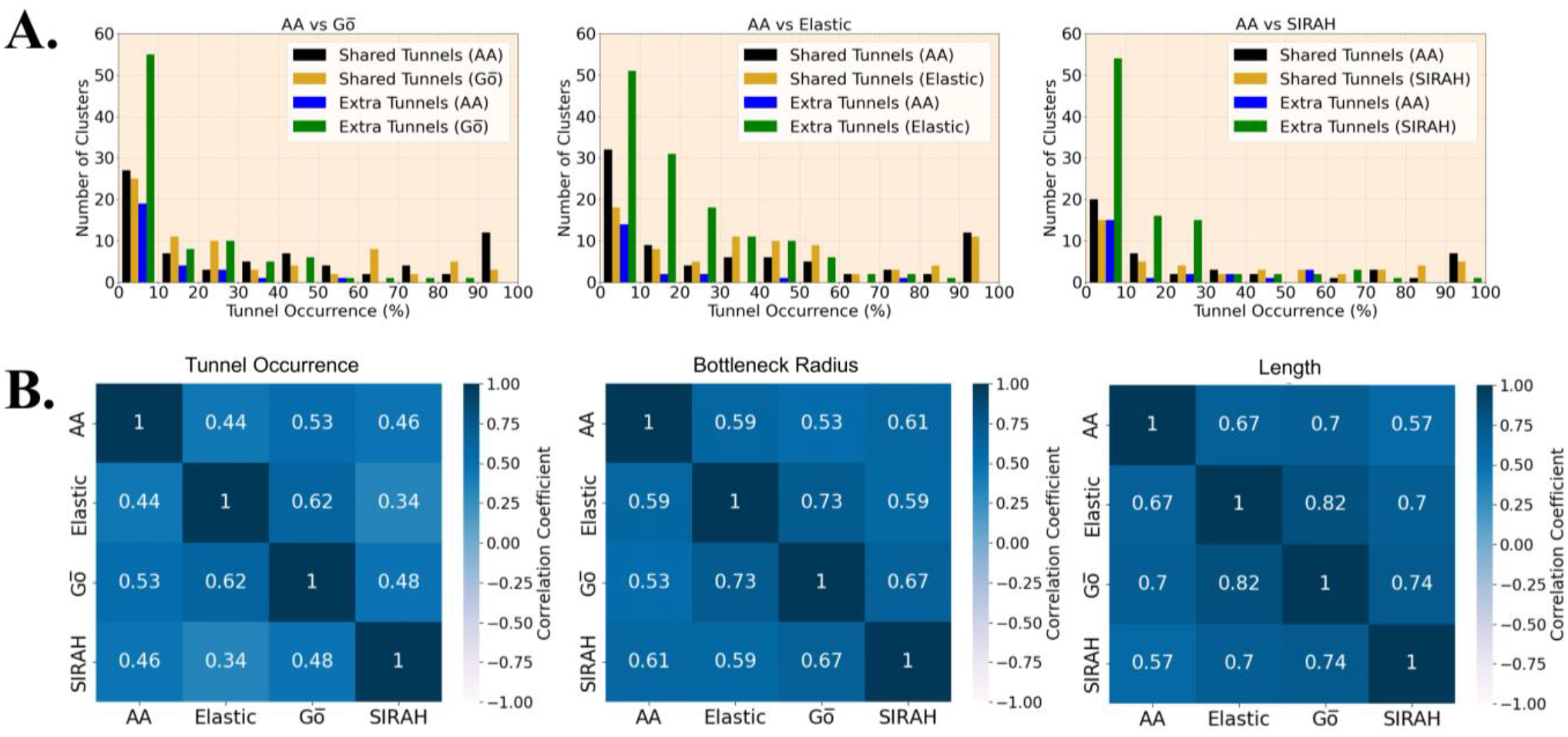
Tunnel co-occurrence and correspondence of their properties between AA and CG methods across all EC classes. **A)** Distribution of tunnel occurrence of the equivalent tunnels shared between AA and CG methods as well as extra tunnels identified exclusively by AA or CG method. See Figure S47 for the plot zoomed in on the extra tunnels. **B**) Heatmaps with Pearson correlation coefficients between properties of tunnels present simultaneously in AA and each CG method across the whole dataset. Data on the statistical significance of the observed correlations are available in Table S8.

## Conclusions

This study revealed a consistently good performance of benchmarked CG methods, i.e., Martini Elastic, Martini Gō, and SIRAH in detecting all well-known functional tunnels in agreement with the AA simulations. The CG tunnel networks include many additional primarily transient tunnels and tunnel branches that were most likely identified due to the increased sampling of protein conformations in CG-MD simulations. However, owing to the coarser nature of the molecular models employed, some additional tunnels could constitute false positive artifacts. Fortunately, the low occurrences of these lower-priority tunnels do not seem to hamper the application of CG models for tunnel analyses. Except for the Elastic model, all CG methods investigated here could distinguish permanent-like and transient tunnels properly. Also, all methods recapitulated well the structural properties of corresponding tunnel ensembles, i.e., their bottleneck radii and lengths. Additionally, very promising results were reached with the Gō model, which was able to accurately reproduce the intricate detail of tunnel engineering effects in LinB variants similar to AA simulations. Importantly, the CG methods provides significant speed up for the simulation in comparison with AA simulations, which significantly reduced the sampling time while accessing more diverse conformational space due to their coarser description of molecular behavior. Despite that CG models, especially Martini Gō, constitute a promising approach to effectively investigate tunnels in proteins, opening new possibilities for identifying tunnels in large protein systems and massive datasets.

## Author Contributions

N.M. setup LinB-Wt and mutants AA models, LinB mutants CG models, EC class 1-3 enzyme CG models, performed all AA and CG simulations, analyzed all tunnels, prepared all figures and drafted the manuscript; N.M. and B.S. devised restraint protocol for stabilization of SIRAH CG model; J.A.S. contributed to setup and optimization of CG simulation protocol with Martini Elastic model; A.B.P. helped setup and optimization of CG simulation protocol with Gō -Martini model for LinB-Wt and supervised its application to other systems; B.S. selected diverse EC2 class enzymes with suitable tunnel networks, and performed their parametrization and optimization of their AA simulation protocol; A.S.T. selected diverse EC3 class enzymes with suitable tunnel networks, and performed their parametrization and optimization of their AA simulation protocol; C.S.B. selected diverse EC1 class enzymes with suitable tunnel networks, and performed their parametrization and optimization of their AA simulation protocol; S.J.M. supervised the setup and optimization of CG simulation protocol with Martini Elastic model; N.M. and J.B. conceived the project and acquired the funding; J.B. coordinated the project and co-written the draft of the manuscript; J.B. and N.M. analyzed the data and interpreted the results; The manuscript was written through the contributions of all authors. All authors have approved the final version of the manuscript.

## Acknowledgments

This work was supported by the National Science Centre, Poland (grant nos. 2023/49/N/NZ2/02567 to N.M., 2022/45/B/NZ1/02519 to A.B.P., and 2017/26/E/NZ1/00548 to J.B.). The training and research stay of N.M. to Molecular Dynamics Group at the University of Groningen was co-funded by the Initiative of Excellence – Research University program of Adam Mickiewicz University (grant no. ID-UB 048/13/UAM/0016). Computations by N.M., B.S., A.S.T, C.S.B., and J.B. were performed at the Poznan Supercomputing and Networking Center. A.B.P. gratefully acknowledges Polish high-performance computing infrastructure PLGrid (HPC Centers: ACK Cyfronet AGH) for providing computer facilities and support within computational grant no. PLG/2024/017332.

## Notes

### Competing Interest Statement

The authors have declared no competing interest.

## References

(1) Monzon, A. M.; Zea, D. J.; Fornasari, M. S.; Saldaño, T. E.; Fernandez-Alberti, S.; Tosatto, S. C. E.; Parisi, G. Conformational Diversity Analysis Reveals Three Functional Mechanisms in Proteins. PLOS Comput. Biol. 2017, 13 (2), e1005398. 10.1371/journal.pcbi.1005398.

(2) Krone, M.; Kozlíková, B.; Lindow, N.; Baaden, M.; Baum, D.; Parulek, J.; Hege, H.-C.; Viola, I. Visual Analysis of Biomolecular Cavities: State of the Art. Comput. Graph. Forum 2016, 35 (3), 527–551. 10.1111/cgf.12928.

(3) Prokop, Z., Gora, A., Brezovsky, J., Chaloupkova, R., Stepankova, V., Damborsky, J. Engineering of Protein Tunnels: Keyhole-Lock-Key Model for Catalysis by the Enzymes with Buried Active Sites; 421-464; Protein Engineering Handbook, Wiley-VCH, Weinheim, 2012.

(4) Petřek, M.; Otyepka, M.; Banáš, P.; Košinová, P.; Koča, J.; Damborský, J. CAVER: A New Tool to Explore Routes from Protein Clefts, Pockets and Cavities. BMC Bioinformatics 2006, 7 (1), 316. 10.1186/1471-2105-7-316.

(5) Pravda, L.; Berka, K.; Svobodová Vařeková, R.; Sehnal, D.; Banáš, P.; Laskowski, R. A.; Koča, J.; Otyepka, M. Anatomy of Enzyme Channels. BMC Bioinformatics 2014, 15 (1), 379. 10.1186/s12859-014-0379-x.

(6) Gora, A.; Brezovsky, J.; Damborsky, J. Gates of Enzymes. Chem. Rev. 2013, 113 (8), 5871–5923. 10.1021/cr300384w.

(7) Marques, S. M.; Daniel, L.; Buryska, T.; Prokop, Z.; Brezovsky, J.; Damborsky, J. Enzyme Tunnels and Gates As Relevant Targets in Drug Design: TUNNELS AND GATES IN DRUG DESIGN. Med. Res. Rev. 2017, 37 (5), 1095–1139. 10.1002/med.21430.

(8) Kingsley, L. J.; Lill, M. A. Substrate Tunnels in Enzymes: Structure-Function Relationships and Computational Methodology: Protein Tunnel Structure-Function Relationship. Proteins Struct. Funct. Bioinforma. 2015, 83 (4), 599–611. 10.1002/prot.24772.

(9) Marques, S. M., Brezovsky, J., Damborsky, J. UNDERSTANDING ENZYMES: FUNCTION, DESIGN, ENGINEERING, AND ANALYSIS; Pan Stanford Publishing, 2016.

(10) Harvey, M. J.; Giupponi, G.; Fabritiis, G. D. ACEMD: Accelerating Biomolecular Dynamics in the Microsecond Time Scale. J. Chem. Theory Comput. 2009, 5 (6), 1632–1639. 10.1021/ct9000685.

(11) Laio, A.; Gervasio, F. L. Metadynamics: A Method to Simulate Rare Events and Reconstruct the Free Energy in Biophysics, Chemistry and Material Science. Rep. Prog. Phys. 2008, 71 (12), 126601. 10.1088/0034-4885/71/12/126601.

(12) Torrie, G. M.; Valleau, J. P. Nonphysical Sampling Distributions in Monte Carlo Free-Energy Estimation: Umbrella Sampling. J. Comput. Phys. 1977, 23 (2), 187–199. 10.1016/0021-9991(77)90121-8.

(13) Mandal, N.; Surpeta, B.; Brezovsky, J. Reinforcing Tunnel Network Exploration in Proteins Using Gaussian Accelerated Molecular Dynamics. J. Chem. Inf. Model. 2024, 64 (16), 6623–6635. 10.1021/acs.jcim.4c00966.

(14) Ingólfsson, H. I.; Lopez, C. A.; Uusitalo, J. J.; De Jong, D. H.; Gopal, S. M.; Periole, X.; Marrink, S. J. The Power of Coarse Graining in Biomolecular Simulations. WIREs Comput. Mol. Sci. 2014, 4 (3), 225–248. 10.1002/wcms.1169.

(15) Kmiecik, S.; Gront, D.; Kolinski, M.; Wieteska, L.; Dawid, A. E.; Kolinski, A. Coarse-Grained Protein Models and Their Applications. Chem. Rev. 2016, 116 (14), 7898–7936. 10.1021/acs.chemrev.6b00163.

(16) Borges-Araújo, L.; Patmanidis, I.; Singh, A. P.; Santos, L. H. S.; Sieradzan, A. K.; Vanni, S.; Czaplewski, C.; Pantano, S.; Shinoda, W.; Monticelli, L.; Liwo, A.; Marrink, S. J.; Souza, P. C. T. Pragmatic Coarse-Graining of Proteins: Models and Applications. J. Chem. Theory Comput. 2023, 19 (20), 7112–7135. 10.1021/acs.jctc.3c00733.

(17) Machado, M. R.; Barrera, E. E.; Klein, F.; Sóñora, M.; Silva, S.; Pantano, S. The SIRAH 2.0 Force Field: Altius, Fortius, Citius. J. Chem. Theory Comput. 2019, 15 (4), 2719–2733. 10.1021/acs.jctc.9b00006.

(18) Periole, X.; Marrink, S.-J. The Martini Coarse-Grained Force Field. In Biomolecular Simulations; Monticelli, L., Salonen, E., Eds.; Methods in Molecular Biology; Humana Press: Totowa, NJ, 2013; Vol. 924, pp 533–565. 10.1007/978-1-62703-017-5_20.

(19) Souza, P. C. T.; Alessandri, R.; Barnoud, J.; Thallmair, S.; Faustino, I.; Grünewald, F.; Patmanidis, I.; Abdizadeh, H.; Bruininks, B. M. H.; Wassenaar, T. A.; Kroon, P. C.; Melcr, J.; Nieto, V.; Corradi, V.; Khan, H. M.; Domański, J.; Javanainen, M.; Martinez-Seara, H.; Reuter, N.; Best, R. B.; Vattulainen, I.; Monticelli, L.; Periole, X.; Tieleman, D. P.; de Vries, A. H.; Marrink, S. J. Martini 3: A General Purpose Force Field for Coarse-Grained Molecular Dynamics. Nat. Methods 2021, 18 (4), 382–388. 10.1038/s41592-021-01098-3.

(20) Periole, X.; Cavalli, M.; Marrink, S.-J.; Ceruso, M. A. Combining an Elastic Network With a Coarse-Grained Molecular Force Field: Structure, Dynamics, and Intermolecular Recognition. J. Chem. Theory Comput. 2009, 5 (9), 2531–2543. 10.1021/ct9002114.

(21) Souza, P. C. T.; Borges-Araújo, L.; Brasnett, C.; Moreira, R. A.; Grünewald, F.; Park, P.; Wang, L.; Razmazma, H.; Borges-Araújo, A. C.; Cofas-Vargas, L. F.; Monticelli, L.; Mera-Adasme, R.; Melo, M. N.; Wu, S.; Marrink, S. J.; Poma, A. B.; Thallmair, S. GōMartini 3: From Large Conformational Changes in Proteins to Environmental Bias Corrections. bioRxiv 2024, 2024.04.15.589479. 10.1101/2024.04.15.589479.

(22) Poma, A. B.; Cieplak, M.; Theodorakis, P. E. Combining the MARTINI and Structure-Based Coarse-Grained Approaches for the Molecular Dynamics Studies of Conformational Transitions in Proteins. J. Chem. Theory Comput. 2017, 13 (3), 1366–1374. 10.1021/acs.jctc.6b00986.

(23) Abraham, M. J.; Murtola, T.; Schulz, R.; Páll, S.; Smith, J. C.; Hess, B.; Lindahl, E. GROMACS: High Performance Molecular Simulations through Multi-Level Parallelism from Laptops to Supercomputers. SoftwareX 2015, 1–2, 19–25. 10.1016/j.softx.2015.06.001.

(24) MacCallum, J. L.; Hu, S.; Lenz, S.; Souza, P. C. T.; Corradi, V.; Tieleman, D. P. An Implementation of the Martini Coarse-Grained Force Field in OpenMM. Biophys. J. 2023, 122 (14), 2864–2870. 10.1016/j.bpj.2023.04.007.

(25) Souza, P. C. T.; Thallmair, S.; Conflitti, P.; Ramírez-Palacios, C.; Alessandri, R.; Raniolo, S.; Limongelli, V.; Marrink, S. J. Protein–Ligand Binding with the Coarse-Grained Martini Model. Nat. Commun. 2020, 11 (1), 3714. 10.1038/s41467-020-17437-5.

(26) D. A. Case; I.Y. Ben-Shalom; S.R. Brozell; D.S. Cerutti; Cheatham, T. E.; V.W.D. Cruzeiro; T.A. Darden; R.E. Duke; D. Ghoreishi; M.K. Gilson; H. Gohlke; A.W. Goetz; D. Greene; Harris, R. C.; N. Homeyer; Yandong Huang; S. Izadi; A. Kovalenko; T. Kurtzman; T.S. Lee; S. LeGrand; P. Li; C. Lin; J. Liu; T. Luchko; R. Luo; D.J. Mermelstein; K.M. Merz; Y. Miao; G. Monard; C. Nguyen; H. Nguyen; I. Omelyan; A. Onufriev; F. Pan; R. Qi; D.R. Roe; A. Roitberg; C. Sagui; S. Schott-Verdugo; Shen, J.; C.L. Simmerling; J. Smith; R. Salomon-Ferrer; J. Swails; R.C. Walker; J. Wang; H. Wei; R.M. Wolf; X. Wu; L. Xiao; D.M. York; P.A. Kollman. Amber 2018, 2018. 10.13140/RG.2.2.31525.68321.

(27) Phillips, J. C.; Hardy, D. J.; Maia, J. D. C.; Stone, J. E.; Ribeiro, J. V.; Bernardi, R. C.; Buch, R.; Fiorin, G.; Hénin, J.; Jiang, W.; McGreevy, R.; Melo, M. C. R.; Radak, B. K.; Skeel, R. D.; Singharoy, A.; Wang, Y.; Roux, B.; Aksimentiev, A.; Luthey-Schulten, Z.; Kalé, L. V.; Schulten, K.; Chipot, C.; Tajkhorshid, E. Scalable Molecular Dynamics on CPU and GPU Architectures with NAMD. J. Chem. Phys. 2020, 153 (4), 044130. 10.1063/5.0014475.

(28) Darré, L.; Machado, M. R.; Dans, P. D.; Herrera, F. E.; Pantano, S. Another Coarse Grain Model for Aqueous Solvation: WAT FOUR? J. Chem. Theory Comput. 2010, 6 (12), 3793–3807. 10.1021/ct100379f.

(29) Klein, F.; Soñora, M.; Helene Santos, L.; Nazareno Frigini, E.; Ballesteros-Casallas, A.; Rodrigo Machado, M.; Pantano, S. The SIRAH Force Field: A Suite for Simulations of Complex Biological Systems at the Coarse-Grained and Multiscale Levels. J. Struct. Biol. 2023, 215 (3), 107985. 10.1016/j.jsb.2023.107985.

(30) Okai, M.; Ohtsuka, J.; Imai, L. F.; Mase, T.; Moriuchi, R.; Tsuda, M.; Nagata, K.; Nagata, Y.; Tanokura, M. Crystal Structure and Site-Directed Mutagenesis Analyses of Haloalkane Dehalogenase LinB from Sphingobium Sp. Strain MI1205. J. Bacteriol. 2013, 195 (11), 2642–2651. 10.1128/JB.02020-12.

(31) Brezovsky, J.; Babkova, P.; Degtjarik, O.; Fortova, A.; Gora, A.; Iermak, I.; Rezacova, P.; Dvorak, P.; Smatanova, I. K.; Prokop, Z.; Chaloupkova, R.; Damborsky, J. Engineering a de Novo Transport Tunnel. ACS Catal. 2016, 6 (11), 7597–7610. 10.1021/acscatal.6b02081.

(32) Gordon, J. C.; Myers, J. B.; Folta, T.; Shoja, V.; Heath, L. S.; Onufriev, A. H++: A Server for Estimating pKas and Adding Missing Hydrogens to Macromolecules. Nucleic Acids Res. 2005, 33 (Web Server), W368–W371. 10.1093/nar/gki464.

(33) Izadi, S.; Anandakrishnan, R.; Onufriev, A. V. Building Water Models: A Different Approach. J. Phys. Chem. Lett. 2014, 5 (21), 3863–3871. 10.1021/jz501780a.

(34) Mermelstein, D. J.; Lin, C.; Nelson, G.; Kretsch, R.; McCammon, J. A.; Walker, R. C. Fast and Flexible Gpu Accelerated Binding Free Energy Calculations within the Amber Molecular Dynamics Package. J. Comput. Chem. 2018, 39 (19), 1354–1358. 10.1002/jcc.25187.

(35) Maier, J. A.; Martinez, C.; Kasavajhala, K.; Wickstrom, L.; Hauser, K. E.; Simmerling, C. ff14SB: Improving the Accuracy of Protein Side Chain and Backbone Parameters from ff99SB. J. Chem. Theory Comput. 2015, 11 (8), 3696–3713. 10.1021/acs.jctc.5b00255.

(36) Zwanzig, R. Nonlinear Generalized Langevin Equations. J. Stat. Phys. 1973, 9 (3), 215–220. 10.1007/BF01008729.

(37) Darden, T.; York, D.; Pedersen, L. Particle Mesh Ewald: An N ⋅log(N) Method for Ewald Sums in Large Systems. J. Chem. Phys. 1993, 98 (12), 10089–10092. 10.1063/1.464397.

(38) Essmann, U.; Perera, L.; Berkowitz, M. L.; Darden, T.; Lee, H.; Pedersen, L. G. A Smooth Particle Mesh Ewald Method. J. Chem. Phys. 1995, 103 (19), 8577–8593. 10.1063/1.470117.

(39) Götz, A. W.; Williamson, M. J.; Xu, D.; Poole, D.; Le Grand, S.; Walker, R. C. Routine Microsecond Molecular Dynamics Simulations with AMBER on GPUs. 1. Generalized Born. J. Chem. Theory Comput. 2012, 8 (5), 1542–1555. 10.1021/ct200909j.

(40) Hopkins, C. W.; Le Grand, S.; Walker, R. C.; Roitberg, A. E. Long-Time-Step Molecular Dynamics through Hydrogen Mass Repartitioning. J. Chem. Theory Comput. 2015, 11 (4), 1864–1874. 10.1021/ct5010406.

(41) Åqvist, J.; Wennerström, P.; Nervall, M.; Bjelic, S.; Brandsdal, B. O. Molecular Dynamics Simulations of Water and Biomolecules with a Monte Carlo Constant Pressure Algorithm. Chem. Phys. Lett. 2004, 384 (4–6), 288–294. 10.1016/j.cplett.2003.12.039.

(42) Berendsen, H. J. C.; Postma, J. P. M.; Van Gunsteren, W. F.; DiNola, A.; Haak, J. R. Molecular Dynamics with Coupling to an External Bath. J. Chem. Phys. 1984, 81 (8), 3684–3690. 10.1063/1.448118.

(43) Basconi, J. E.; Shirts, M. R. Effects of Temperature Control Algorithms on Transport Properties and Kinetics in Molecular Dynamics Simulations. J. Chem. Theory Comput. 2013, 9 (7), 2887–2899. 10.1021/ct400109a.

(44) Roe, D. R.; Cheatham, T. E. PTRAJ and CPPTRAJ: Software for Processing and Analysis of Molecular Dynamics Trajectory Data. J. Chem. Theory Comput. 2013, 9 (7), 3084–3095. 10.1021/ct400341p.

(45) Machado, M. R.; Pantano, S. SIRAH Tools: Mapping, Backmapping and Visualization of Coarse-Grained Models. Bioinformatics 2016, 32 (10), 1568–1570. 10.1093/bioinformatics/btw020.

(46) Machado, M. R.; Pantano, S. Split the Charge Difference in Two! A Rule of Thumb for Adding Proper Amounts of Ions in MD Simulations. J. Chem. Theory Comput. 2020, 16 (3), 1367–1372. 10.1021/acs.jctc.9b00953.

(47) De Jong, D. H.; Liguori, N.; Van Den Berg, T.; Arnarez, C.; Periole, X.; Marrink, S. J. Atomistic and Coarse Grain Topologies for the Cofactors Associated with the Photosystem II Core Complex. J. Phys. Chem. B 2015, 119 (25), 7791–7803. 10.1021/acs.jpcb.5b00809.

(48) Mahmood, Md. I.; Poma, A. B.; Okazaki, K. Optimizing Gō-MARTINI Coarse-Grained Model for F-BAR Protein on Lipid Membrane. Front. Mol. Biosci. 2021, 8, 619381. 10.3389/fmolb.2021.619381.

(49) Liu, Z.; Moreira, R. A.; Dujmović, A.; Liu, H.; Yang, B.; Poma, A. B.; Nash, M. A. Mapping Mechanostable Pulling Geometries of a Therapeutic Anticalin/CTLA-4 Protein Complex. Nano Lett. 2022, 22 (1), 179–187. 10.1021/acs.nanolett.1c03584.

(50) Kroon, P.; Grunewald, F.; Barnoud, J.; van Tilburg, M.; Souza, P.; Wassenaar, T.; Marrink, S. Martinize2 and Vermouth: Unified Framework for Topology Generation. eLife12 2024, RP90627. 10.7554/elife.90627.2.

(51) Gorelov, S.; Titov, A.; Tolicheva, O.; Konevega, A.; Shvetsov, A. DSSP in GROMACS: Tool for Defining Secondary Structures of Proteins in Trajectories. J. Chem. Inf. Model. 2024, 64 (9), 3593–3598. 10.1021/acs.jcim.3c01344.

(52) Bussi, G.; Donadio, D.; Parrinello, M. Canonical Sampling through Velocity Rescaling. J. Chem. Phys. 2007, 126 (1), 014101. 10.1063/1.2408420.

(53) Bernetti, M.; Bussi, G. Pressure Control Using Stochastic Cell Rescaling. J. Chem. Phys. 2020, 153 (11), 114107. 10.1063/5.0020514.

(54) Ke, Q.; Gong, X.; Liao, S.; Duan, C.; Li, L. Effects of Thermostats/Barostats on Physical Properties of Liquids by Molecular Dynamics Simulations. J. Mol. Liq. 2022, 365, 120116. 10.1016/j.molliq.2022.120116.

(55) Sousa, F. M.; Lima, L. M. P.; Arnarez, C.; Pereira, M. M.; Melo, M. N. Coarse-Grained Parameterization of Nucleotide Cofactors and Metabolites: Protonation Constants, Partition Coefficients, and Model Topologies. J. Chem. Inf. Model. 2021, 61 (1), 335–346. 10.1021/acs.jcim.0c01077.

(56) Chovancova, E.; Pavelka, A.; Benes, P.; Strnad, O.; Brezovsky, J.; Kozlikova, B.; Gora, A.; Sustr, V.; Klvana, M.; Medek, P.; Biedermannova, L.; Sochor, J.; Damborsky, J. CAVER 3.0: A Tool for the Analysis of Transport Pathways in Dynamic Protein Structures. PLoS Comput. Biol. 2012, 8 (10), e1002708. 10.1371/journal.pcbi.1002708.

(57) Sequeiros-Borja, C.; Surpeta, B.; Marchlewski, I.; Brezovsky, J. Divide-and-Conquer Approach to Study Protein Tunnels in Long Molecular Dynamics Simulations. MethodsX 2023, 10, 101968. 10.1016/j.mex.2022.101968.

(58) Brezovsky, J.; Thirunavukarasu, A. S.; Surpeta, B.; Sequeiros-Borja, C. E.; Mandal, N.; Sarkar, D. K.; Dongmo Foumthuim, C. J.; Agrawal, N. TransportTools: A Library for High-Throughput Analyses of Internal Voids in Biomolecules and Ligand Transport through Them. Bioinformatics 2022, 38 (6), 1752–1753. 10.1093/bioinformatics/btab872.

(59) Sequeiros-Borja, C.; Surpeta, B.; Thirunavukarasu, A. S.; Dongmo Foumthuim, C. J.; Marchlewski, I.; Brezovsky, J. Water Will Find Its Way: Transport through Narrow Tunnels in Hydrolases. J. Chem. Inf. Model. 2024, 64 (15), 6014–6025. 10.1021/acs.jcim.4c00094.

(60) Wassenaar, T. A.; Pluhackova, K.; Böckmann, R. A.; Marrink, S. J.; Tieleman, D. P. Going Backward: A Flexible Geometric Approach to Reverse Transformation from Coarse Grained to Atomistic Models. J. Chem. Theory Comput. 2014, 10 (2), 676–690. 10.1021/ct400617g.

(61) Virtanen, P.; Gommers, R.; Oliphant, T. E.; Haberland, M.; Reddy, T.; Cournapeau, D.; Burovski, E.; Peterson, P.; Weckesser, W.; Bright, J.; Van Der Walt, S. J.; Brett, M.; Wilson, J.; Millman, K. J.; Mayorov, N.; Nelson, A. R. J.; Jones, E.; Kern, R.; Larson, E.; Carey, C. J.; Polat, İ.; Feng, Y.; Moore, E. W.; VanderPlas, J.; Laxalde, D.; Perktold, J.; Cimrman, R.; Henriksen, I.; Quintero, E. A.; Harris, C. R.; Archibald, A. M.; Ribeiro, A. H.; Pedregosa, F.; Van Mulbregt, P.; SciPy 1.0 Contributors; Vijaykumar, A.; Bardelli, A. P.; Rothberg, A.; Hilboll, A.; Kloeckner, A.; Scopatz, A.; Lee, A.; Rokem, A.; Woods, C. N.; Fulton, C.; Masson, C.; Häggström, C.; Fitzgerald, C.; Nicholson, D. A.; Hagen, D. R.; Pasechnik, D. V.; Olivetti, E.; Martin, E.; Wieser, E.; Silva, F.; Lenders, F.; Wilhelm, F.; Young, G.; Price, G. A.; Ingold, G.-L.; Allen, G. E.; Lee, G. R.; Audren, H.; Probst, I.; Dietrich, J. P.; Silterra, J.; Webber, J. T.; Slavič, J.; Nothman, J.; Buchner, J.; Kulick, J.; Schönberger, J. L.; De Miranda Cardoso, J. V.; Reimer, J.; Harrington, J.; Rodríguez, J. L. C.; Nunez-Iglesias, J.; Kuczynski, J.; Tritz, K.; Thoma, M.; Newville, M.; Kümmerer, M.; Bolingbroke, M.; Tartre, M.; Pak, M.; Smith, N. J.; Nowaczyk, N.; Shebanov, N.; Pavlyk, O.; Brodtkorb, P. A.; Lee, P.; McGibbon, R. T.; Feldbauer, R.; Lewis, S.; Tygier, S.; Sievert, S.; Vigna, S.; Peterson, S.; More, S.; Pudlik, T.; Oshima, T.; Pingel, T. J.; Robitaille, T. P.; Spura, T.; Jones, T. R.; Cera, T.; Leslie, T.; Zito, T.; Krauss, T.; Upadhyay, U.; Halchenko, Y. O.; Vázquez-Baeza, Y. SciPy 1.0: Fundamental Algorithms for Scientific Computing in Python. Nat. Methods 2020, 17 (3), 261–272. 10.1038/s41592-019-0686-2.

(62) Sarkar, D. K.; Surpeta, B.; Brezovsky, J. Incorporating Prior Knowledge in the Seeds of Adaptive Sampling Molecular Dynamics Simulations of Ligand Transport in Enzymes with Buried Active Sites. J. Chem. Theory Comput. 2024, acs.jctc.4c00452. 10.1021/acs.jctc.4c00452.

(63) Brezovsky, J.; Chovancova, E.; Gora, A.; Pavelka, A.; Biedermannova, L.; Damborsky, J. Software Tools for Identification, Visualization and Analysis of Protein Tunnels and Channels. Biotechnol. Adv. 2013, 31 (1), 38–49. 10.1016/j.biotechadv.2012.02.002.

(64) Javanainen, M.; Martinez-Seara, H.; Vattulainen, I. Excessive Aggregation of Membrane Proteins in the Martini Model. PLOS ONE 2017, 12 (11), e0187936. 10.1371/journal.pone.0187936.

(65) Stark, A. C.; Andrews, C. T.; Elcock, A. H. Toward Optimized Potential Functions for Protein– Protein Interactions in Aqueous Solutions: Osmotic Second Virial Coefficient Calculations Using the MARTINI Coarse-Grained Force Field. J. Chem. Theory Comput. 2013, 9 (9), 4176–4185. 10.1021/ct400008p.

(66) Majumder, A.; Straub, J. E. Addressing the Excessive Aggregation of Membrane Proteins in the MARTINI Model. J. Chem. Theory Comput. 2021, 17 (4), 2513–2521. 10.1021/acs.jctc.0c01253.

(67) McDonald, A. G.; Tipton, K. F. Enzyme Nomenclature and Classification: The State of the Art. FEBS J. 2023, 290 (9), 2214–2231. 10.1111/febs.16274.

