## Supporting Information Figures and Tables for "Benchmarking coarse-grained simulation methods for investigation of transport tunnels in enzymes"

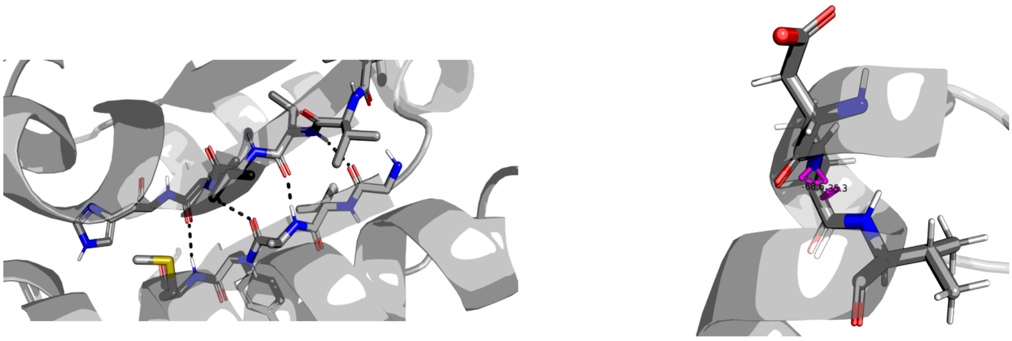

H-bond restraint on β-sheets    Dihedral angle restraint on α-helix

**Figure S1. Restraints applied to SIRAH CG simulations to keep the α-helix and β-sheets stable**. H-bond-mimicking restraints were applied as 20 kcal/mol/Å$^2$ and dihedral angle restraints stabilizing helices were applied as 4 kcal/mol/Å$^2$.

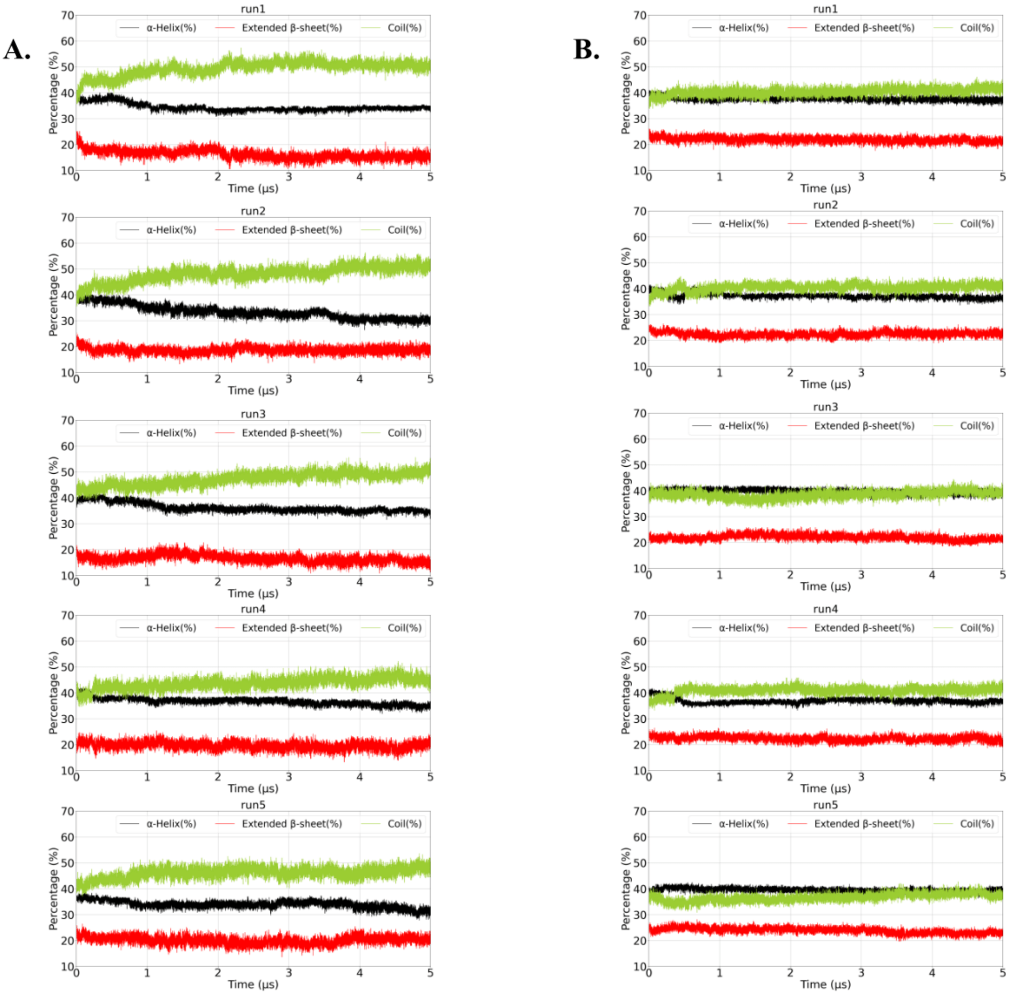


**Figure S2. Secondary structure analysis in LinB-Wt simulations with SIRAH CG model**. Outputs from SIRAH-tools shows decreasing the fraction of structure present as coil and stability in the content of α-helix and β-sheet. (A) without the restraints and (B) with the use of restraints.

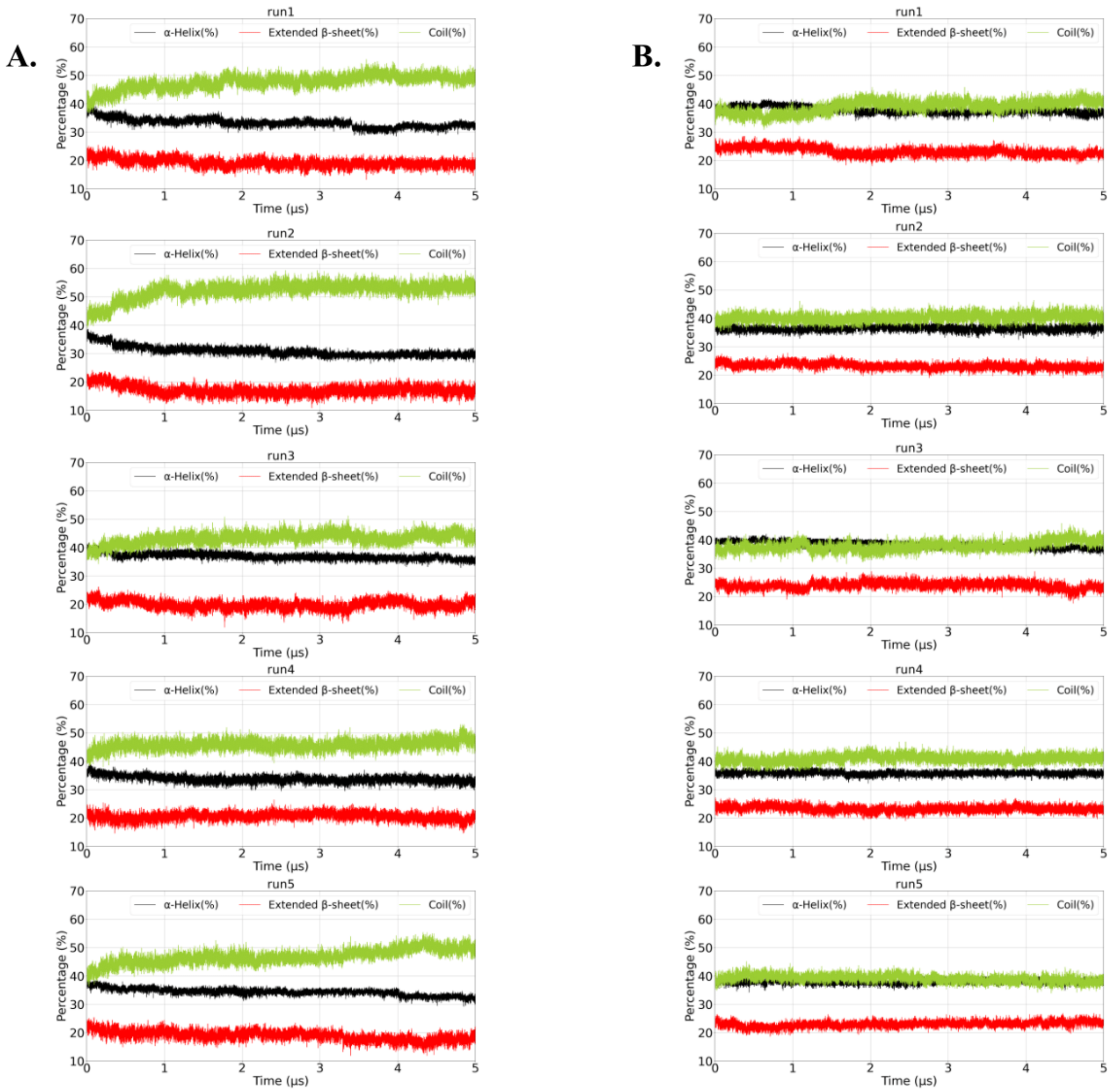


**Figure S3. Secondary structure analysis in LinB-Closed simulations with SIRAH CG model.** Outputs from SIRAH-tools shows decreasing the fraction of structure present as coil and stability in the content of α-helix and β-sheet. (A) without the restraints and (B) with the use of restraints.

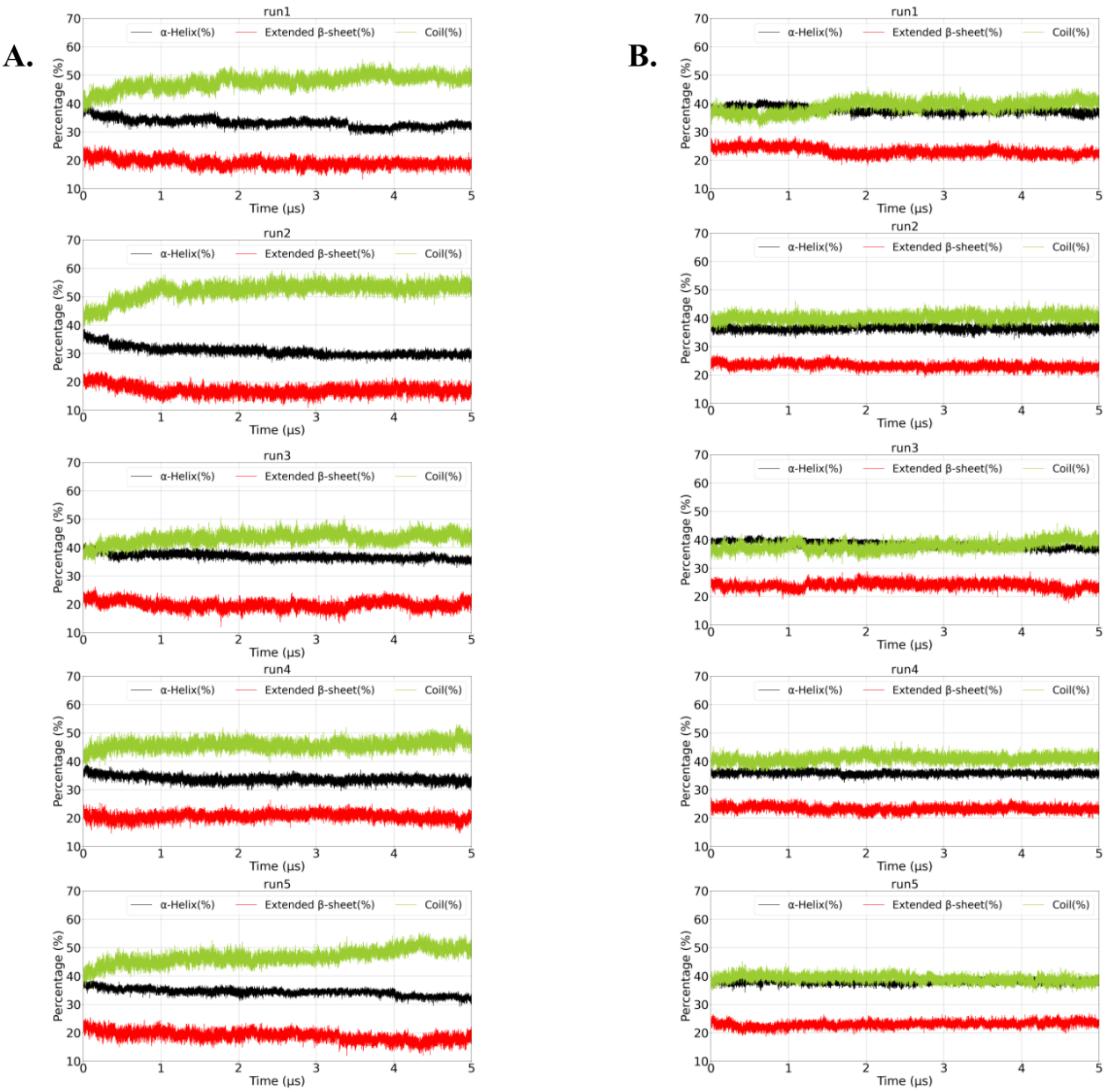


**Figure S4. Secondary structure analysis in LinB-Open simulations with SIRAH CG model.** Outputs from SIRAH-tools shows decreasing the fraction of structure present as coil and stability in the content of α-helix and β-sheet. (A) without the restraints and (B) with the use of restraints.

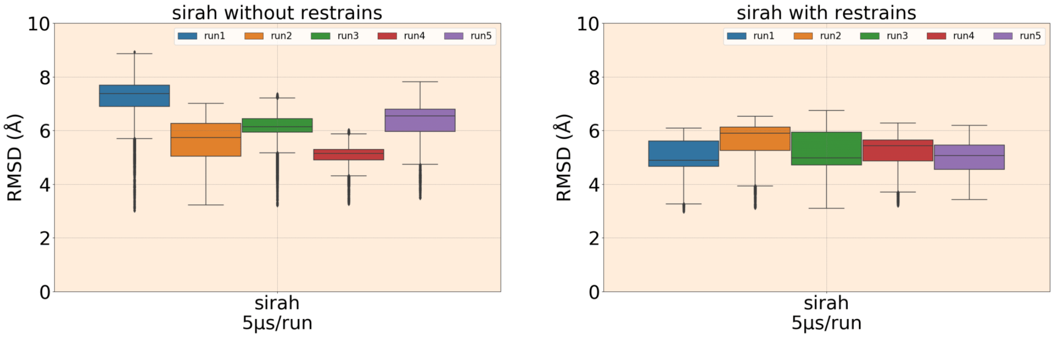


**Figure S5. RMSD of LinB-Wt with and without restraints in SIRAH, all 5 replicas (run 1-5).** The box plot shows median (middle line in the box) and the box represents the interquartile range, which is range from 25th percentile to 75th percentile and gives a sense of how spread out the middle 50% data is. The whiskers extending from the box represents range of the data.

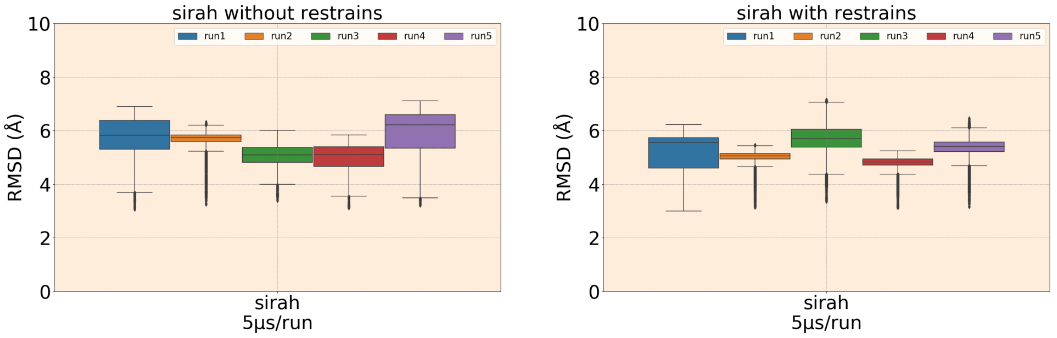


**Figure S6. RMSD of LinB-Closed mutant with and without restraints in SIRAH, all 5 replicas (run 1-5).** The box plot shows median (middle line in the box) and the box represents the interquartile range, which is range from 25th percentile to 75th percentile and gives a sense of how spread out the middle 50% data is. The whiskers extending from the box represents range of the data.

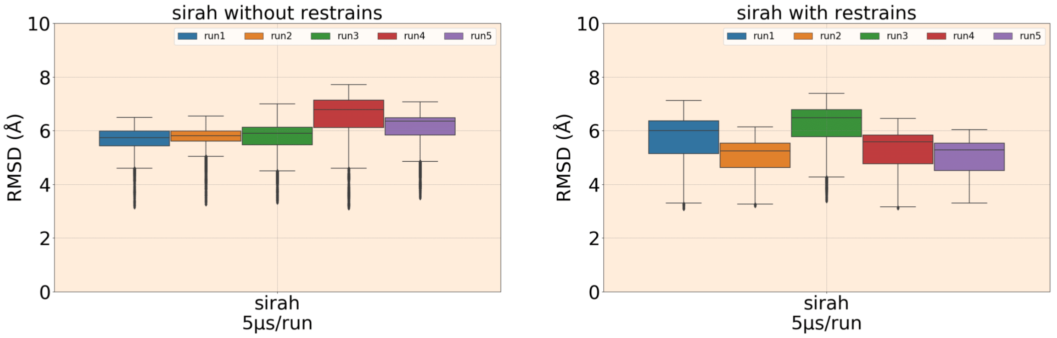


**Figure S7. RMSD of LinB-Open mutant with and without restraints in SIRAH, all 5 replicas (run 1-5).** The box plot shows median (middle line in the box) and the box represents the interquartile range, which is range from 25th percentile to 75th percentile and gives a sense of how spread out the middle 50% data is. The whiskers extending from the box represents range of the data.

**Table S1. The numer of extra contacts/bonds/restraints used to structurally bias CG simulations to maintain stability of LinB variants in individual methods.**

| Enzyme | Initial structure ID | Elastic | Gō | SIRAH (dihedral / distance) |
|---|---|---|---|---|
| LinB-Wt | 1 | 1341 | 657 | 210 / 33 |
| | 2 | 1334 | 701 | 216 / 41 |
| | 3 | 1319 | 693 | 214 / 37 |
| | 4 | 1319 | 693 | 200 / 37 |
| | 5 | 1351 | 701 | 202 / 38 |
| LinB-Closed | 1 | 1344 | 700 | 204 / 37 |
| | 2 | 1324 | 698 | 206 / 24 |
| | 3 | 1332 | 700 | 216 / 39 |
| | 4 | 1343 | 694 | 202 / 36 |
| | 5 | 1325 | 696 | 214 / 39 |
| LinB-Open | 1 | 1364 | 698 | 194 / 37 |
| | 2 | 1354 | 700 | 208 / 24 |
| | 3 | 1367 | 691 | 206 / 37 |
| | 4 | 1332 | 694 | 198 / 40 |
| | 5 | 1364 | 698 | 200 / 37 |

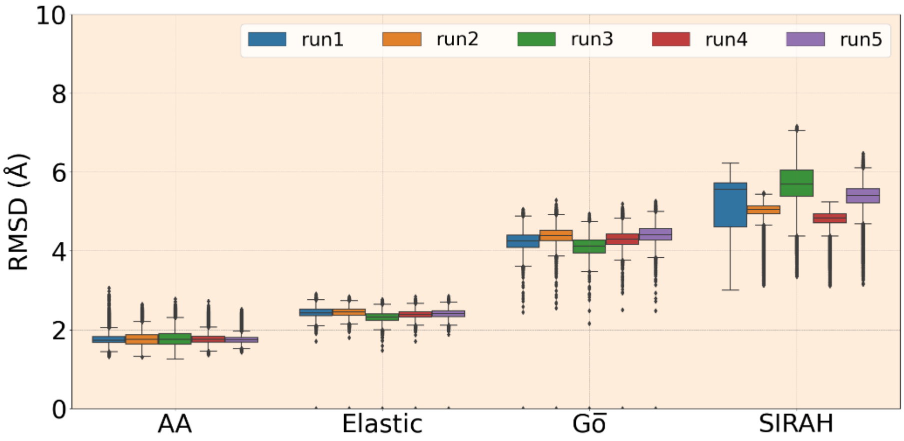


**Figure S8. RMSD of whole LinB-Closed protein with all residues in five replicates using AA and CG methods – SIRAH, Elastic, and Gō.** The box plot shows median (middle line in the box) and the box represents the interquartile range, which is range from 25$^{th}$ percentile to 75$^{th}$ percentile and gives a sense of how spread out the middle 50% data is. The whiskers extending from the box represents range of the data.

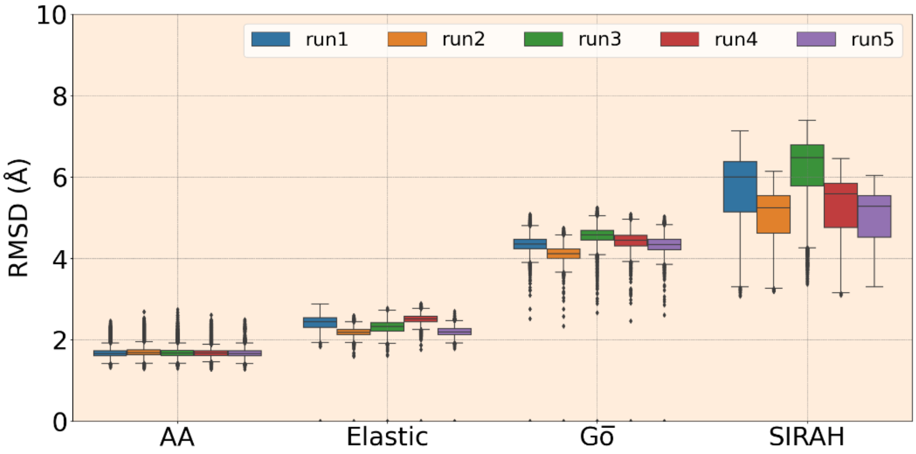


**Figure S9. RMSD of whole LinB-Open protein with all residues in five replicates using AA and CG methods – SIRAH, Elastic, and Gō.** The box plot shows median (middle line in the box) and the box represents the interquartile range, which is range from 25$^{th}$ percentile to 75$^{th}$ percentile and gives a sense of how spread out the middle 50% data is. The whiskers extending from the box represents range of the data.

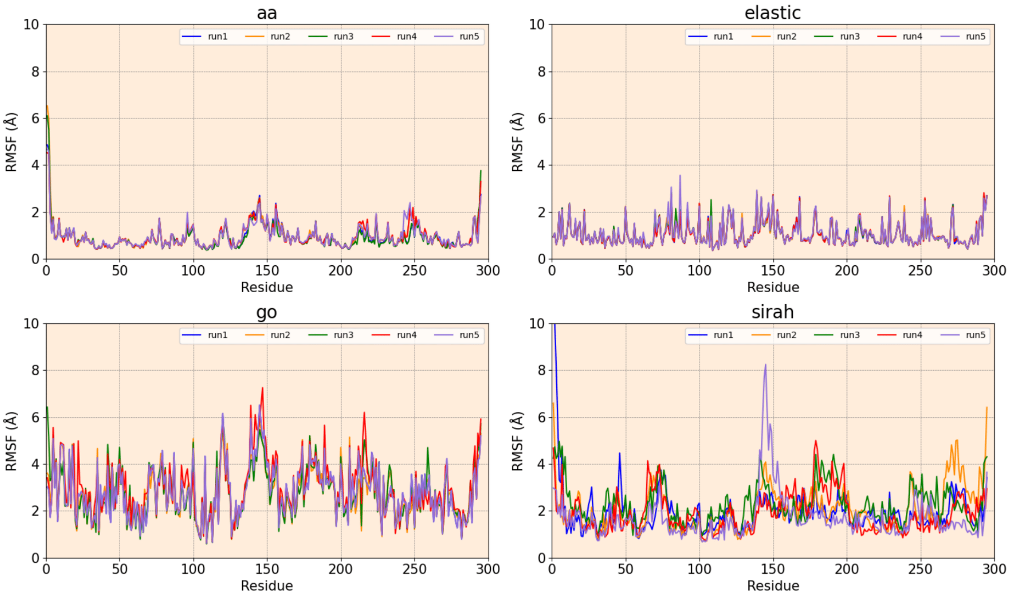


**Figure S10. RMSF of LinB-Wt using all the methods – AA, Gō, Elastic, and SIRAH, comprising 5 replicas of 5 µs each.**

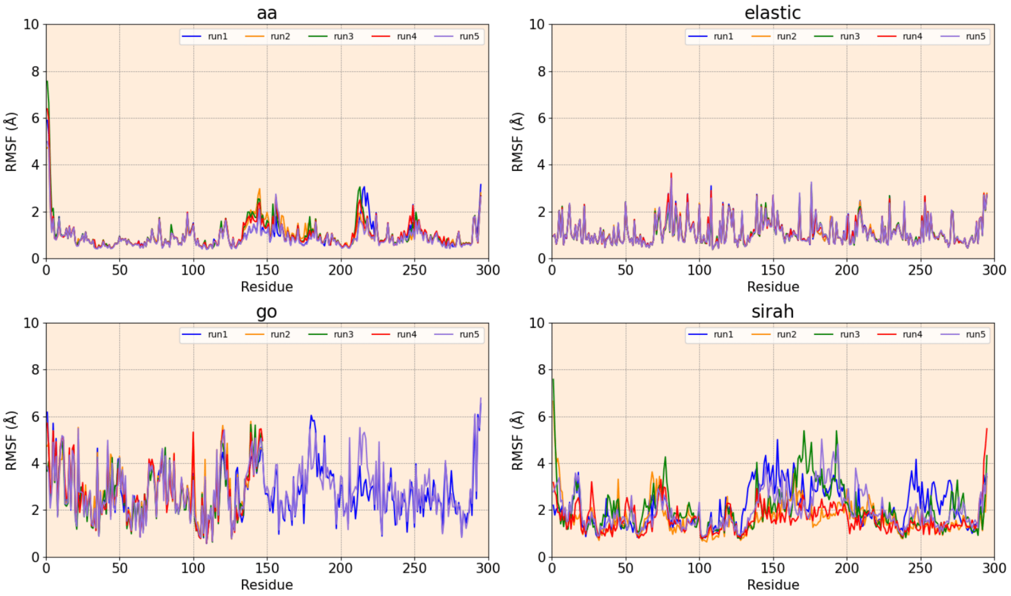


**Figure S11. RMSF of LinB-Closed mutant using all the methods – AA, Gō, Elastic, and SIRAH, comprising 5 replicas of 5 µs each.**

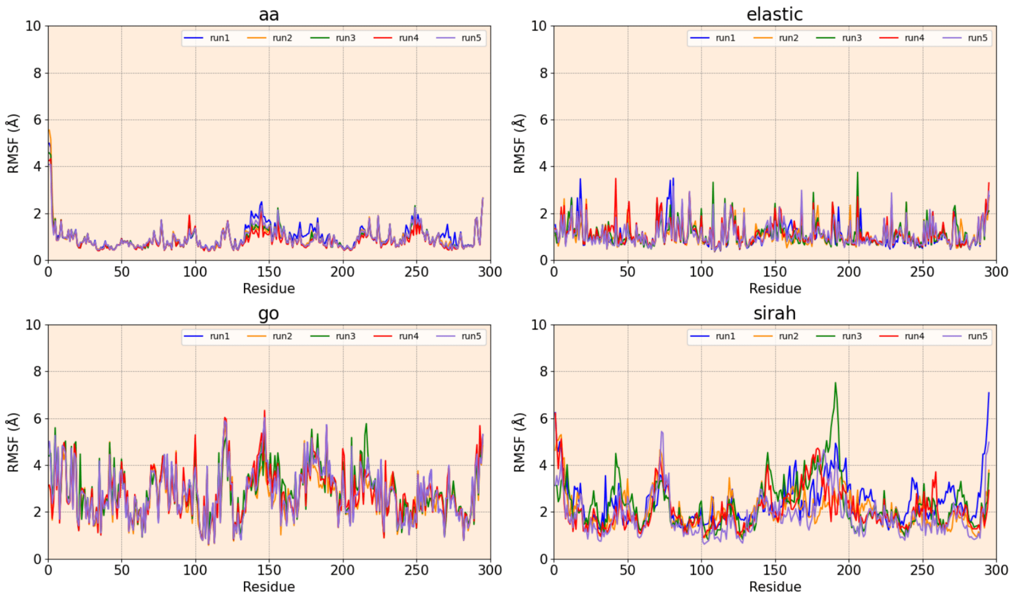


**Figure S12. RMSF of LinB-Open mutant using all the methods – AA, Gō, Elastic, and SIRAH, comprising 5 replicas of 5 μs each.**

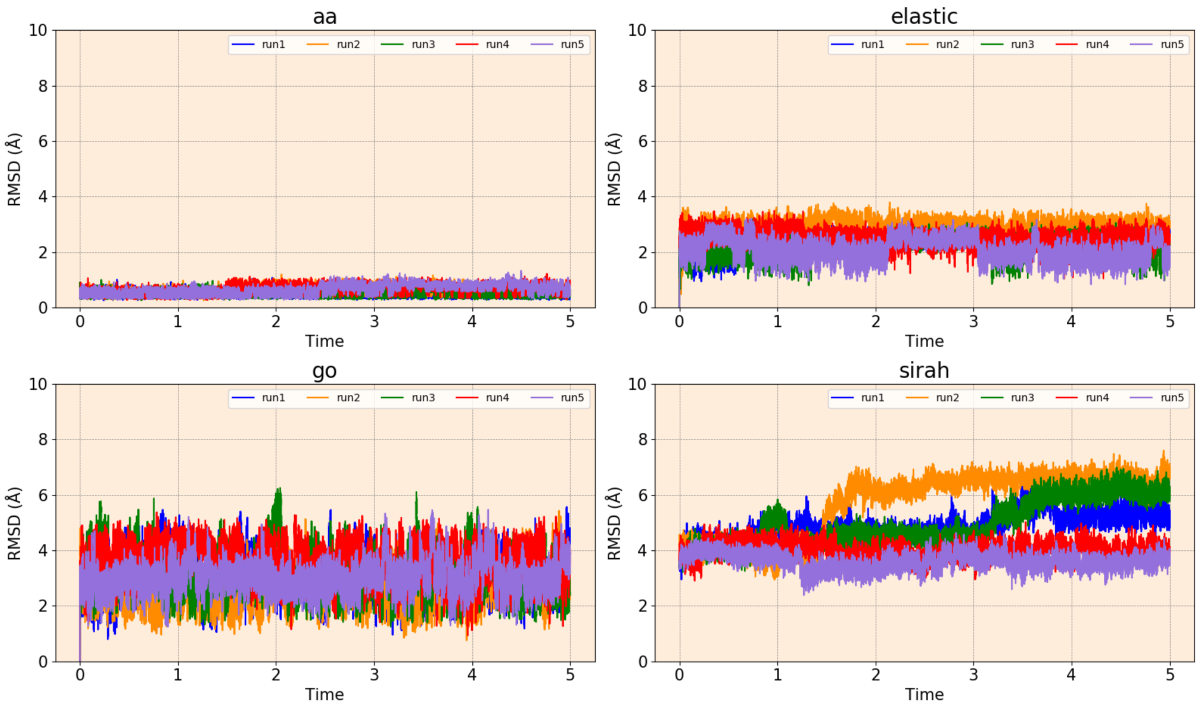


**Figure S13. RMSD evolution of catalytic pentad of LinB in 5 replicas of 5 μs simulations with AA, Gō, Elastic, and SIRAH methods.**

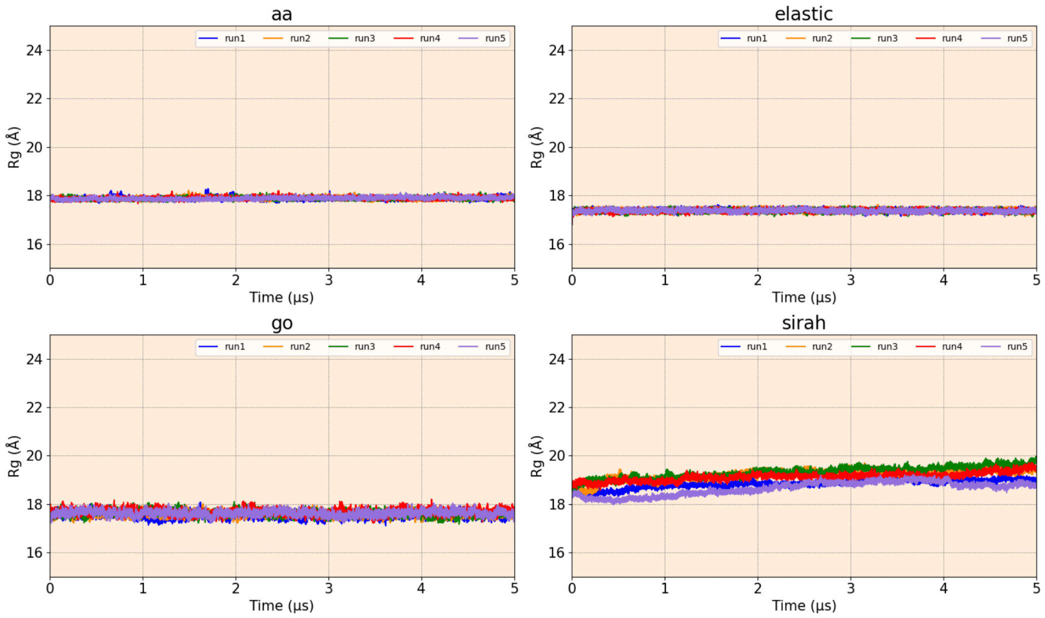


**Figure S14. Rg of LinB-Wt using all the methods – AA, Gō, Elastic, and SIRAH, comprising 5 replicas of 5 µs each.**

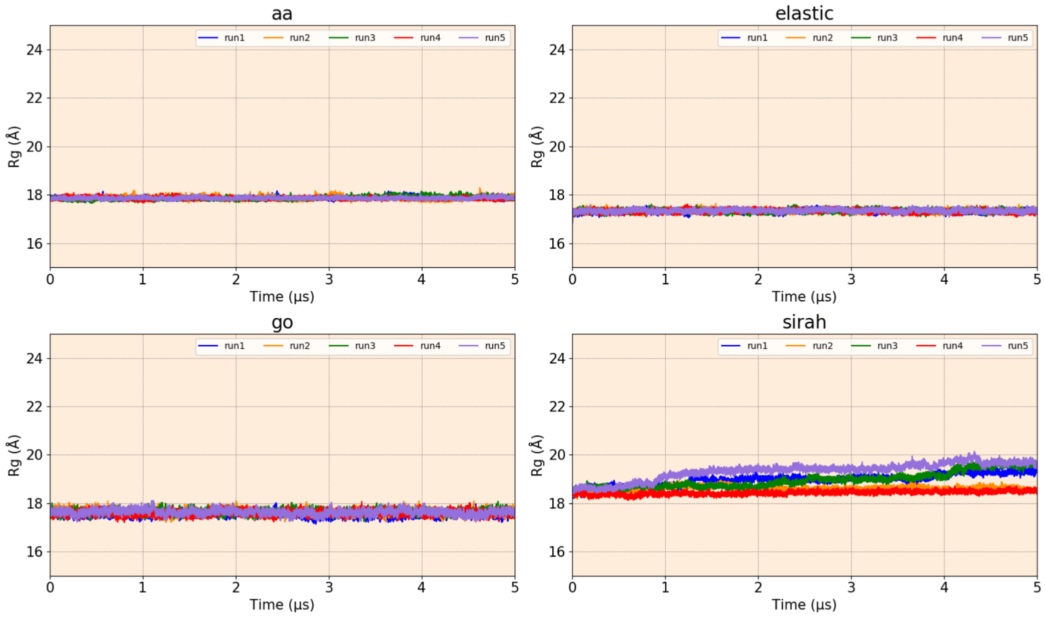


**Figure S15. Rg of LinB-Closed using all the methods – AA, Gō, Elastic, and SIRAH, comprising 5 replicas of 5 µs each.**

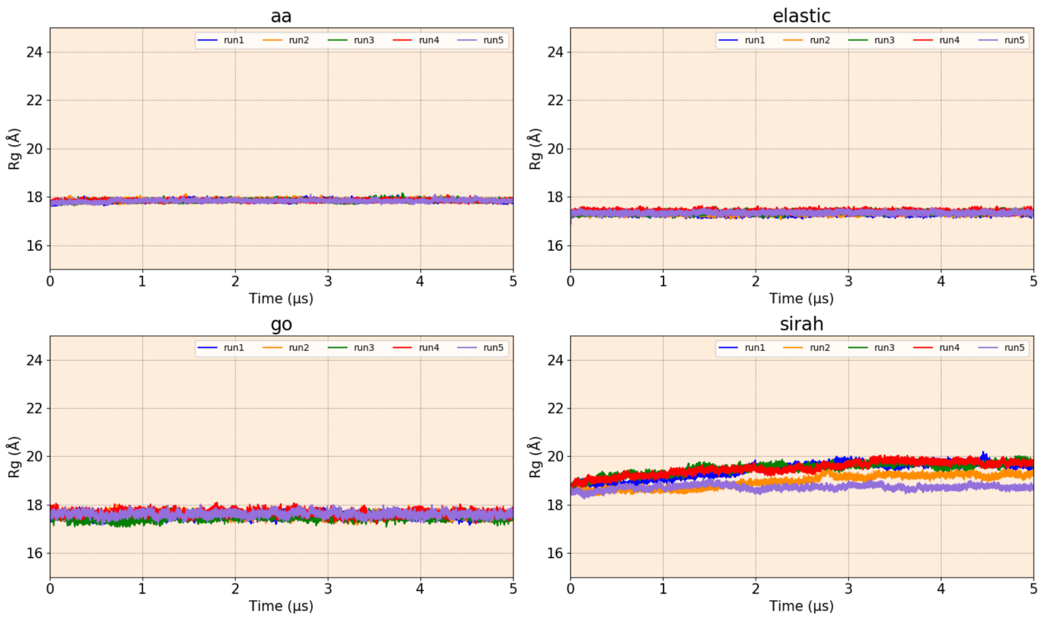


**Figure S16. Rg of LinB-Open using all the methods – AA, Gō, Elastic, and SIRAH, comprising 5 replicas of 5 µs each.**

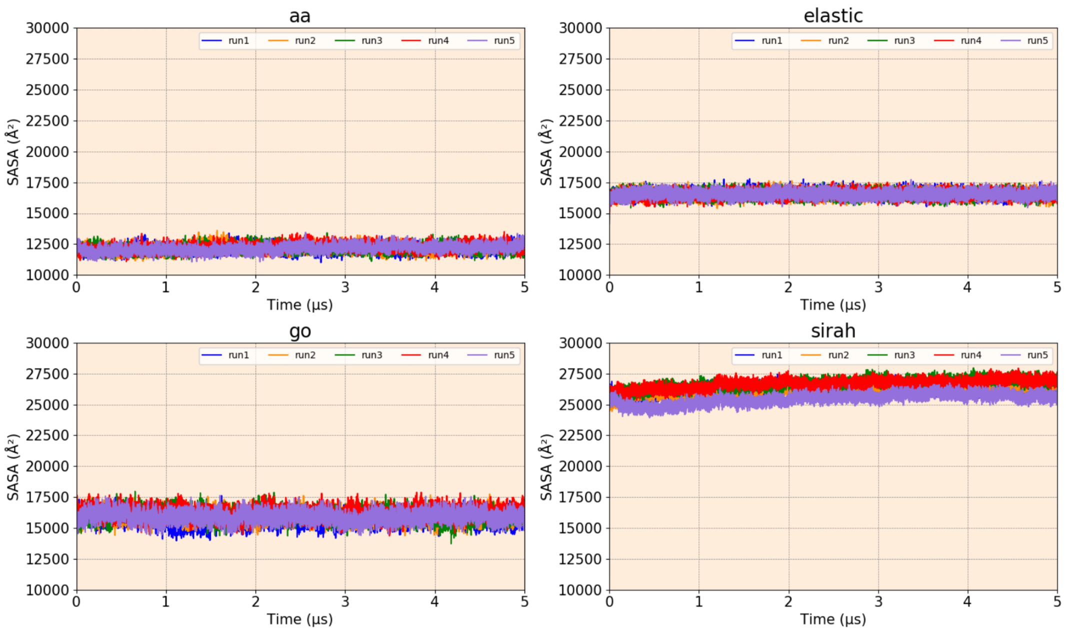


**Figure S17. SASA of LinB-Wt using all the methods – AA, Gō, Elastic, and SIRAH, comprising 5 replicas of 5 µs each.**

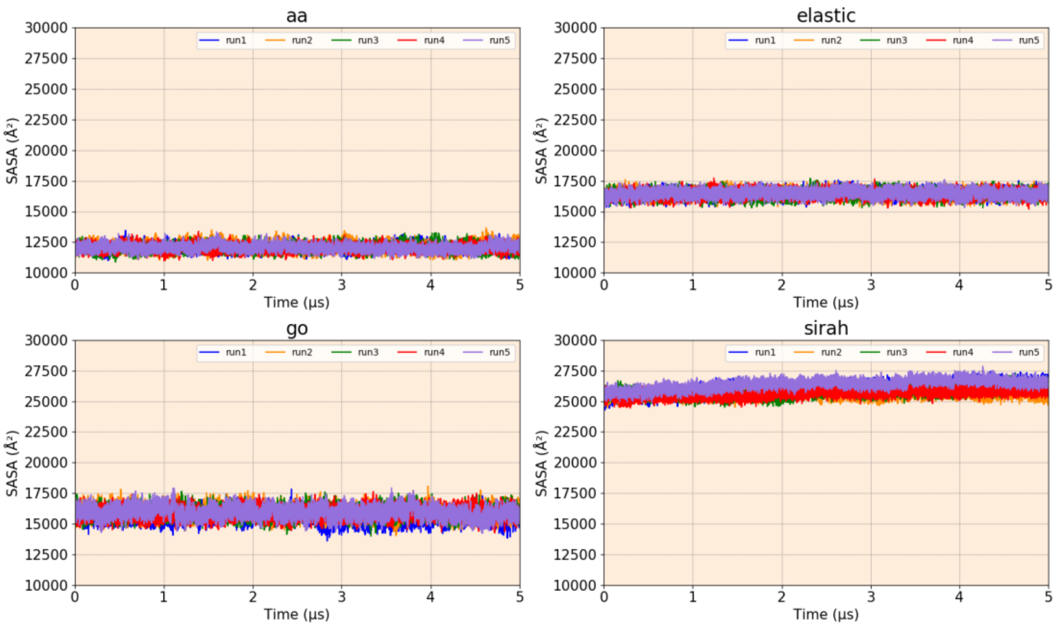


**Figure S18. SASA of LinB-Closed using all the methods – AA, Gō, Elastic, and SIRAH, comprising 5 replicas of 5 µs each.**

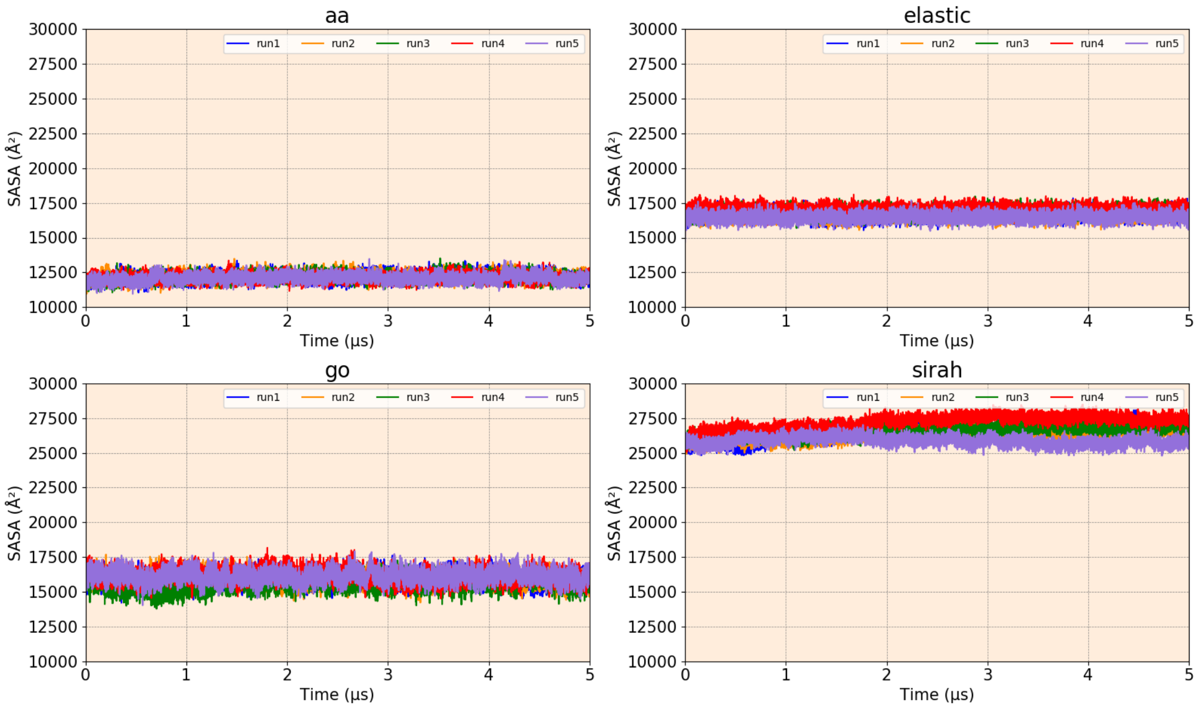


**Figure S19. SASA of LinB-Open using all the methods – AA, Gō, Elastic, and SIRAH, comprising 5 replicas of 5 µs each.**

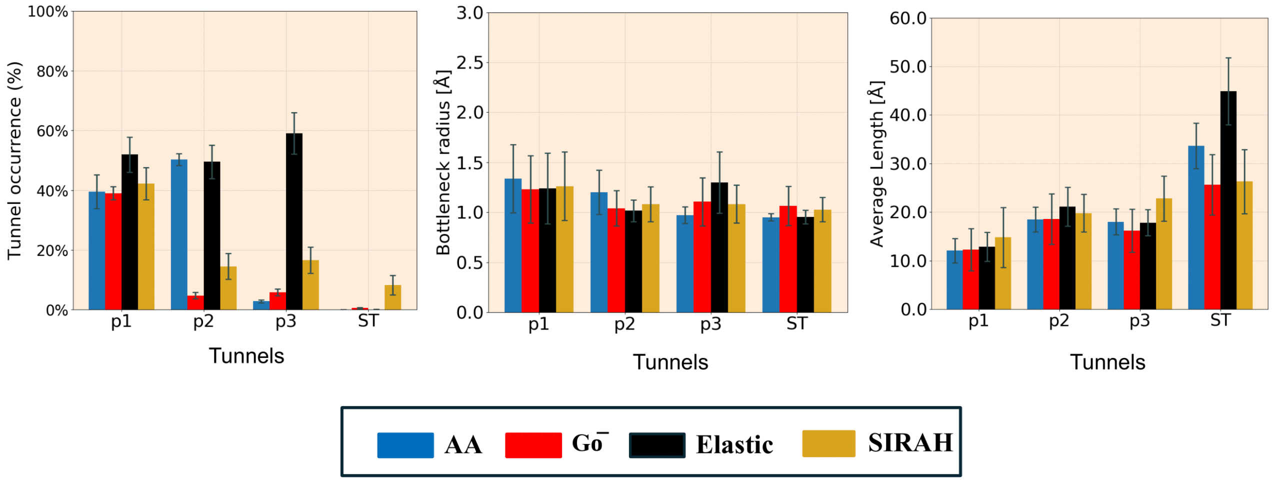


**Figure S20. Properties of main tunnels captured in LinB-Closed mutant using different methods.** Data represents average±SD obtained from 5 simulation replicas. See Table S3 for details on statistical significance

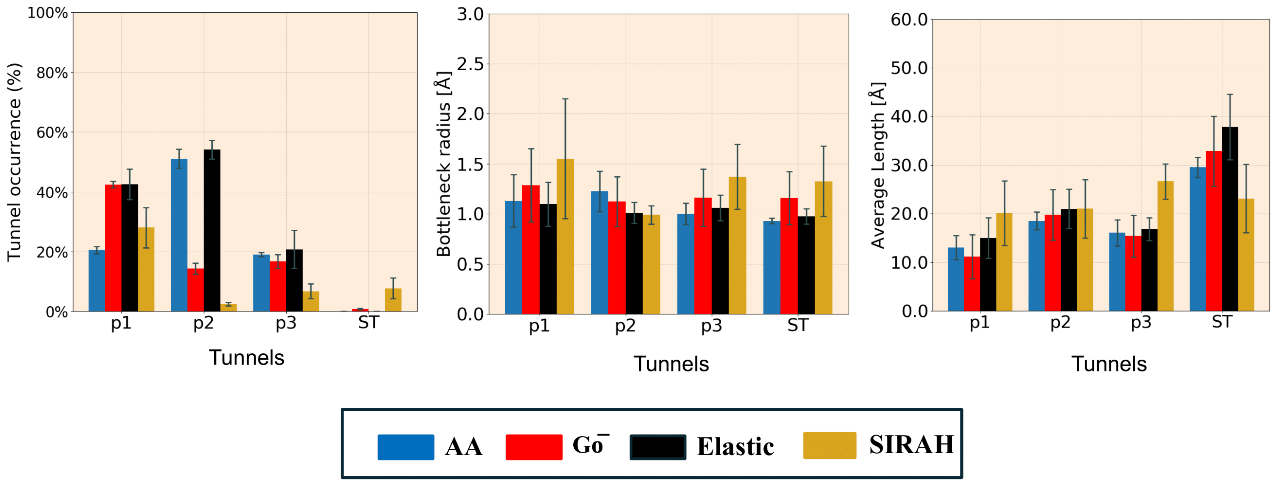


**Figure S21. Properties of main tunnels captured in LinB-Open mutant using different methods.** Data represents average±SD obtained from 5 simulation replicas. See Table S4 for details on statistical significance

**Table S2. Statistical significance tests contrasting tunnel properties in LinB-Wt between AA and CG methods.**

| Property | Tunnel | AA versus SIRAH | | AA versus Elastic | | AA versus Gō | |
|---|---|---|---|---|---|---|---|
| | | T-statistics | P-value | T-statistics | P-value | T-statistics | P-value |
| Occurrence | P1 | 22.72 | **0.00** | 7.71 | **0.00** | 12.39 | **0.00** |
| | P2 | 49.56 | **0.00** | -25.11 | **0.00** | 39.41 | **0.00** |
| | P3 | 7.24 | **0.00** | -92.4 | **0.00** | -9.68 | **0.00** |
| | ST | -4.67 | **0.01** | 6.53 | **0.00** | -9.16 | **0.00** |
| Bottleneck radius | P1 | 0.53 | 0.62 | 0.51 | 0.64 | 0.65 | 0.55 |
| | P2 | 1.61 | 0.18 | 1.20 | 0.29 | 0.91 | 0.41 |
| | P3 | -1.54 | 0.19 | -2.29 | 0.08 | -1.26 | 0.27 |
| | ST | -1.51 | 0.20 | -0.28 | 0.79 | -2.04 | 0.11 |
| Length | P1 | -0.37 | 0.73 | -2.69 | 0.05 | 0.48 | 0.66 |
| | P2 | -2.17 | 0.09 | -1.52 | 0.20 | -1.24 | 0.28 |
| | P3 | -3.27 | **0.03** | 1.00 | 0.37 | 0.42 | 0.69 |
| | ST | 3.16 | **0.03** | -2.05 | 0.10 | -0.04 | 0.97 |

**Table S3. Statistical significance tests contrasting tunnel properties in LinB-Closed between AA and CG methods.**

| Property | Tunnel | AA versus SIRAH | | AA versus Elastic | | AA versus Gō | |
|---|---|---|---|---|---|---|---|
| | | T-statistics | P-value | T-statistics | P-value | T-statistics | P-value |
| Occurrence | P1 | -0.79 | 0.47 | -3.41 | **0.03** | 0.17 | 0.87 |
| | P2 | 16.85 | **0.00** | 0.28 | 0.78 | 45.78 | **0.00** |
| | P3 | -6.98 | **0.00** | -18.19 | **0.00** | -5.35 | **0.01** |
| | ST | -5.63 | **0.00** | -4.59 | **0.01** | -5.59 | **0.00** |
| Bottleneck radius | P1 | 0.35 | 0.74 | 0.45 | 0.67 | 0.50 | 0.64 |
| | P2 | 0.91 | 0.39 | 1.65 | 0.17 | 1.26 | 0.27 |
| | P3 | -1.21 | 0.29 | -2.29 | 0.08 | -1.17 | 0.31 |
| | ST | -1.36 | 0.25 | -0.03 | 0.97 | -1.29 | 0.27 |
| Length | P1 | -0.92 | 0.41 | -0.47 | 0.67 | -0.09 | 0.93 |
| | P2 | -0.61 | 0.57 | -1.23 | 0.29 | -0.02 | 0.98 |
| | P3 | -1.99 | 0.11 | 0.10 | 0.92 | 0.79 | 0.47 |
| | ST | 2.04 | 0.11 | -3.00 | **0.04** | 2.31 | 0.08 |

**Table S4. Statistical significance tests contrasting tunnel properties in LinB-Open between AA and CG methods.**

| Property | Tunnel | AA versus SIRAH | | AA versus Elastic | | AA versus Gō | |
|---|---|---|---|---|---|---|---|
| | | T-statistics | P-value | T-statistics | P-value | T-statistics | P-value |
| Occurrence | P1 | -2.46 | 0.07 | -9.39 | **0.00** | -30.6 | **0.00** |
| | P2 | 33.57 | **0.00** | -1.56 | 0.19 | 22.4 | **0.00** |
| | P3 | 10.82 | **0.00** | -0.59 | 0.58 | 2.19 | 0.09 |
| | ST | -4.99 | **0.01** | -1.56 | 0.19 | -7.37 | **0.00** |
| Bottleneck radius | P1 | -1.45 | 0.22 | 0.20 | 0.85 | -0.78 | 0.48 |
| | P2 | 2.35 | 0.07 | 2.10 | 0.10 | 0.71 | 0.52 |
| | P3 | -2.44 | 0.07 | -0.79 | 0.47 | -1.20 | 0.29 |
| | ST | -2.52 | 0.06 | -1.22 | 0.29 | -1.93 | 0.13 |
| Length | P1 | -2.21 | 0.09 | -0.90 | 0.41 | 0.81 | 0.46 |
| | P2 | -0.89 | 0.42 | -1.24 | 0.28 | -0.52 | 0.63 |
| | P3 | -5.20 | **0.00** | -0.47 | 0.66 | 0.30 | 0.78 |
| | ST | 1.96 | 0.12 | -2.62 | 0.06 | -0.98 | 0.38 |

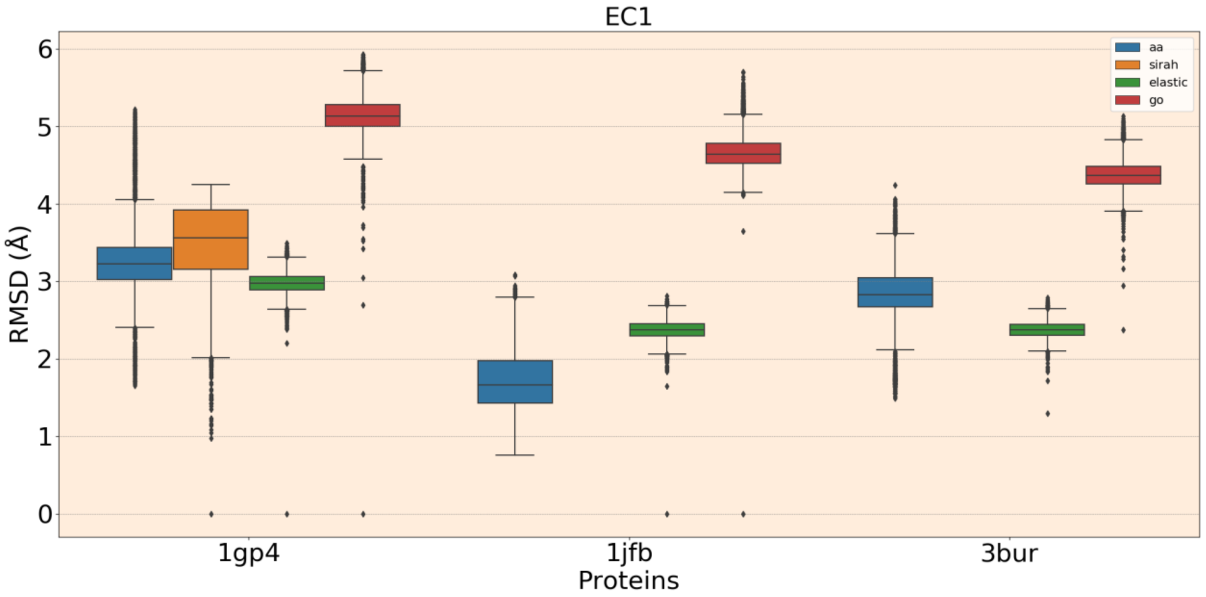


**Figure S23. RMSD of enzymes from EC1 class with all residues in one replicates using AA and CG methods – SIRAH, Elastic, and Gō̄.** The box plot shows median (middle line in the box) and the box represents the interquartile range, which is range from 25th percentile to 75th percentile and gives a sense of how spread out the middle 50% data is. The whiskers extending from the box represents range of the data. Data from SIRAH simulation of 1fjb and 3bur enzymes are not available due to the lack of parameters for the cofactors

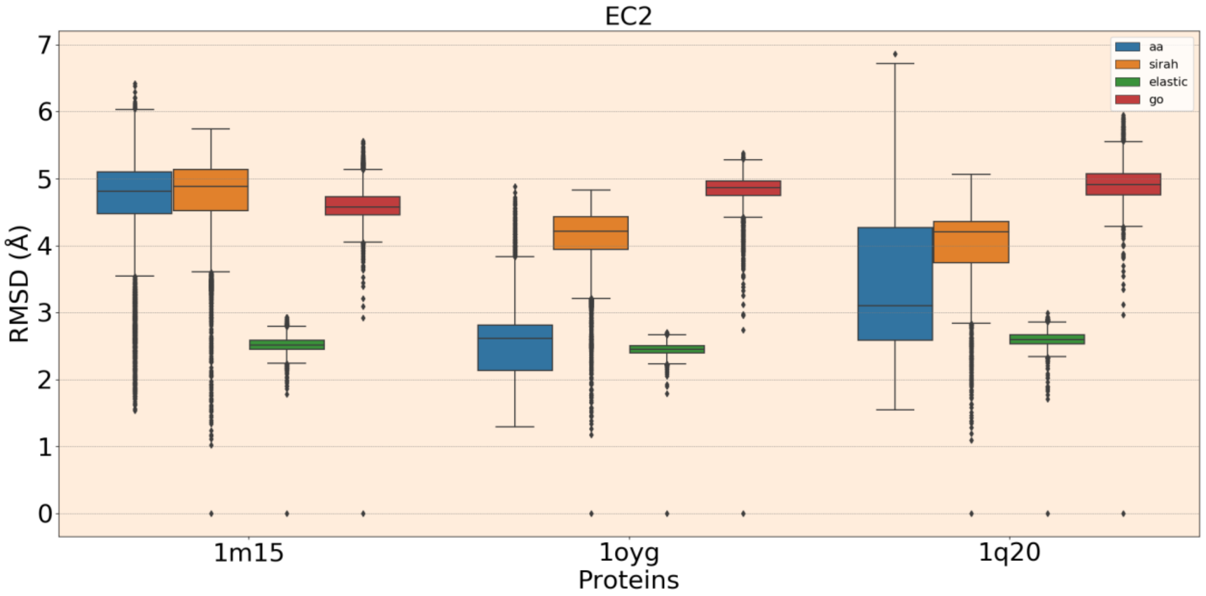


**Figure S24. RMSD of enzymes from EC2 class with all residues in one replicates using AA and CG methods – SIRAH, Elastic, and Gō̄.** The box plot shows median (middle line in the box) and the box represents the interquartile range, which is range from 25th percentile to 75th percentile and gives a sense of how spread out the middle 50% data is. The whiskers extending from the box represents range of the data.

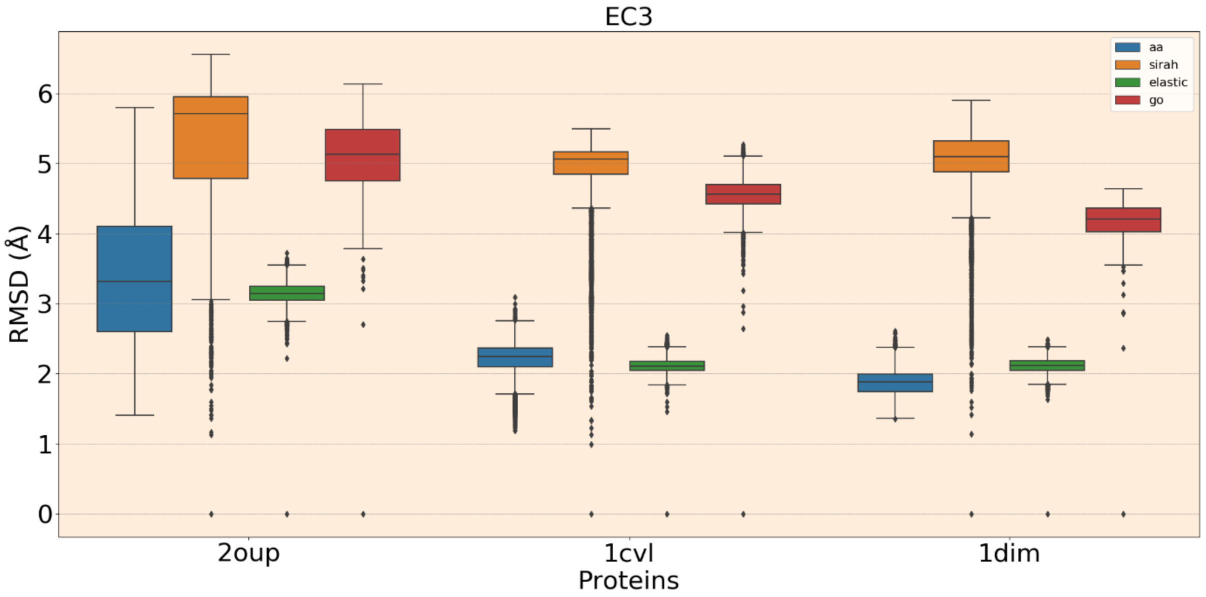


**Figure S25. RMSD of enzymes from EC3 class with all residues in one replicates using AA and CG methods – SIRAH, Elastic, and Gō.** The box plot shows median (middle line in the box) and the box represents the interquartile range, which is range from 25th percentile to 75th percentile and gives a sense of how spread out the middle 50% data is. The whiskers extending from the box represents range of the data.

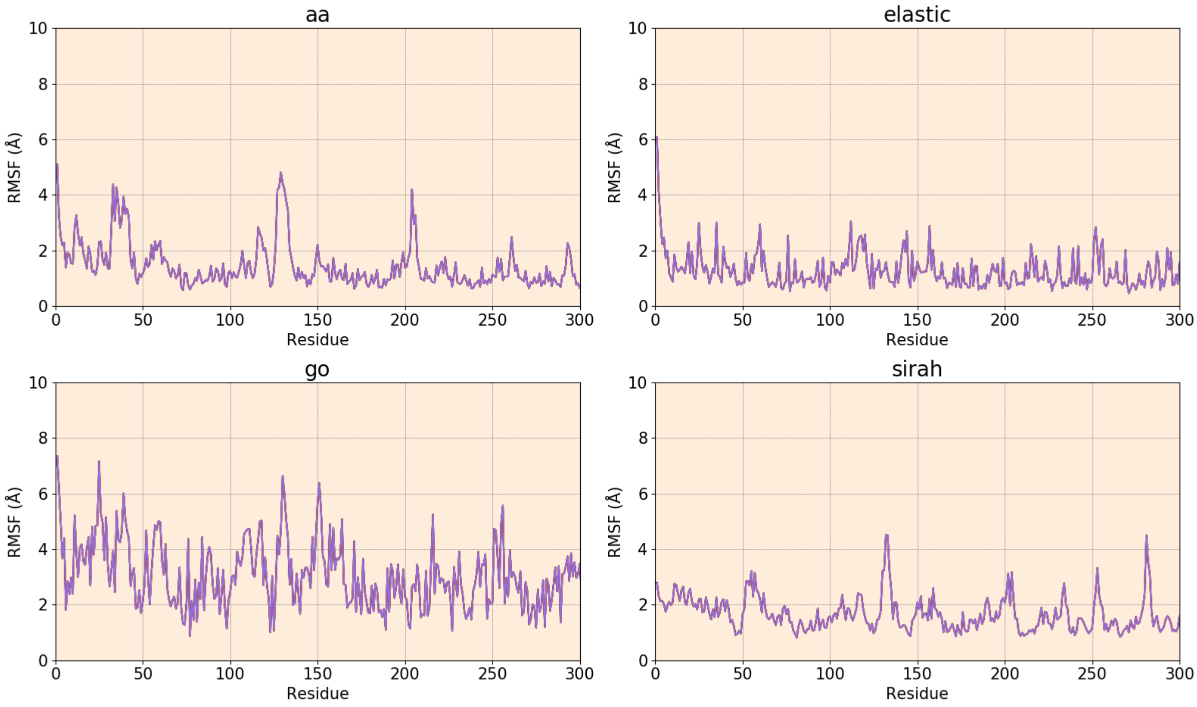


**Figure S26. RMSF of EC1 class enzyme 1gp4 using all the methods – AA, Gō, Elastic, and SIRAH.**

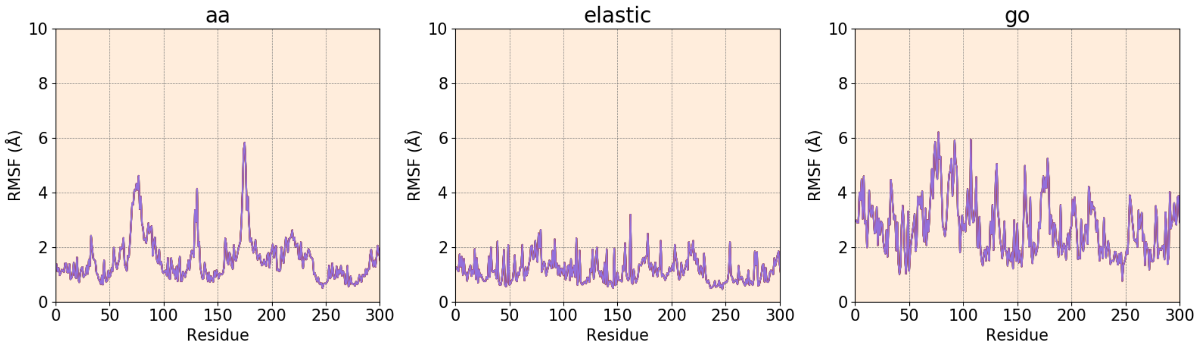


**Figure S27. RMSF of EC1 class enzyme 1jfb using three methods – AA, Gō, and Elastic.** SIRAH simulation is not available due to the lack of parameters for the cofactor.

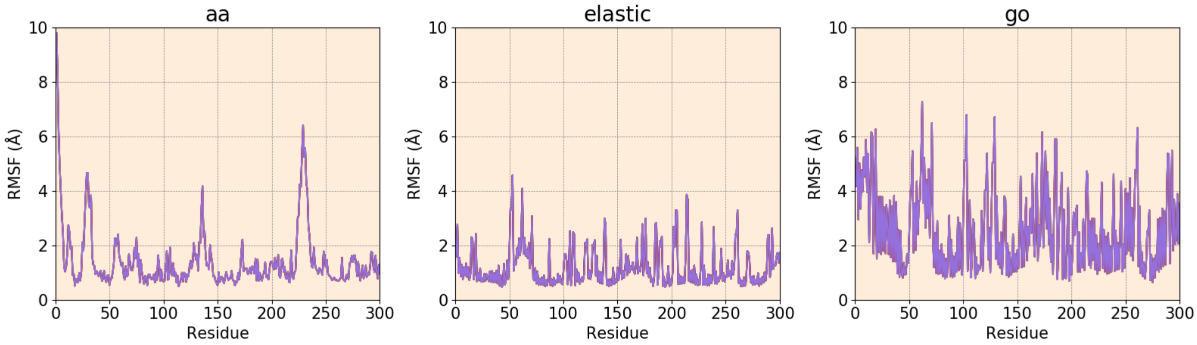


**Figure S28. RMSF of EC1 class enzyme 3bur using three methods – AA, Gō, and Elastic.** SIRAH simulation is not available due to the lack of parameters for the cofactor.

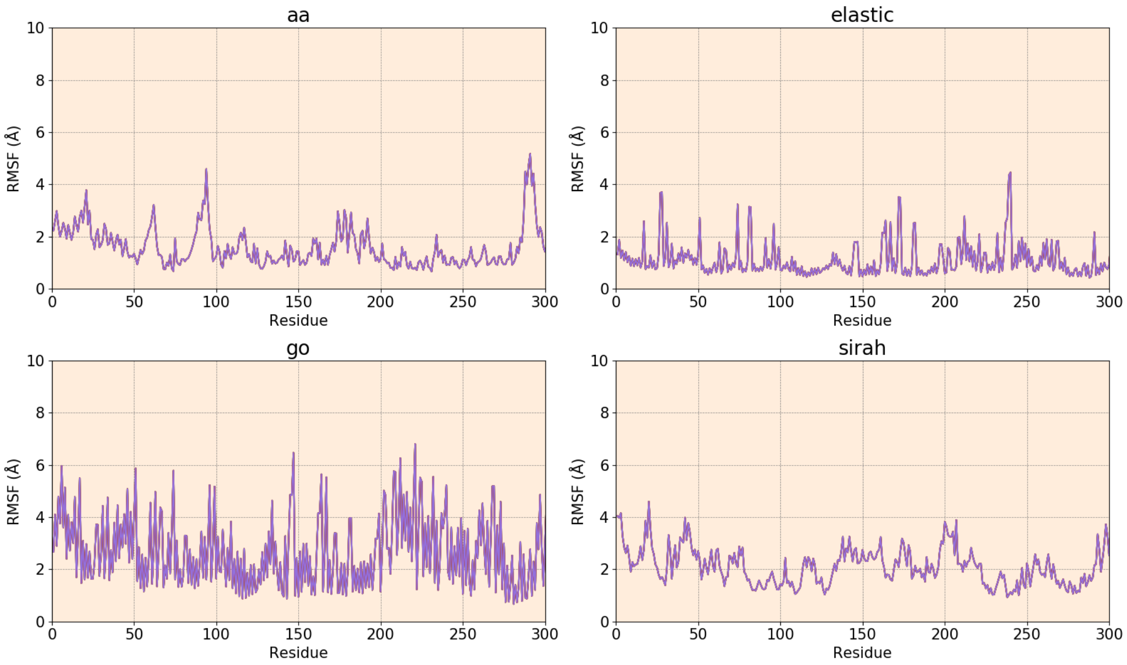


**Figure S29. RMSF of EC2 class enzyme 1m15 using all the methods – AA, Gō, Elastic, and SIRAH.**

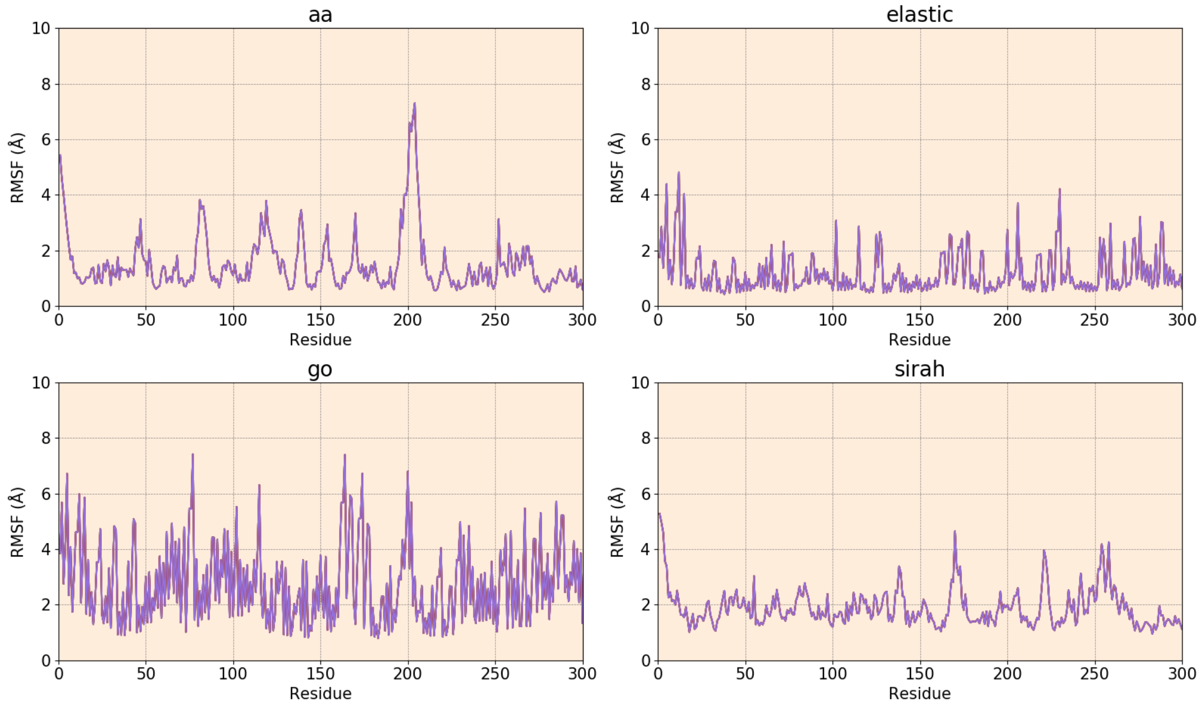


**Figure S30. RMSF of EC2 class enzyme 1oyg using all the methods – AA, Gō, Elastic, and SIRAH.**

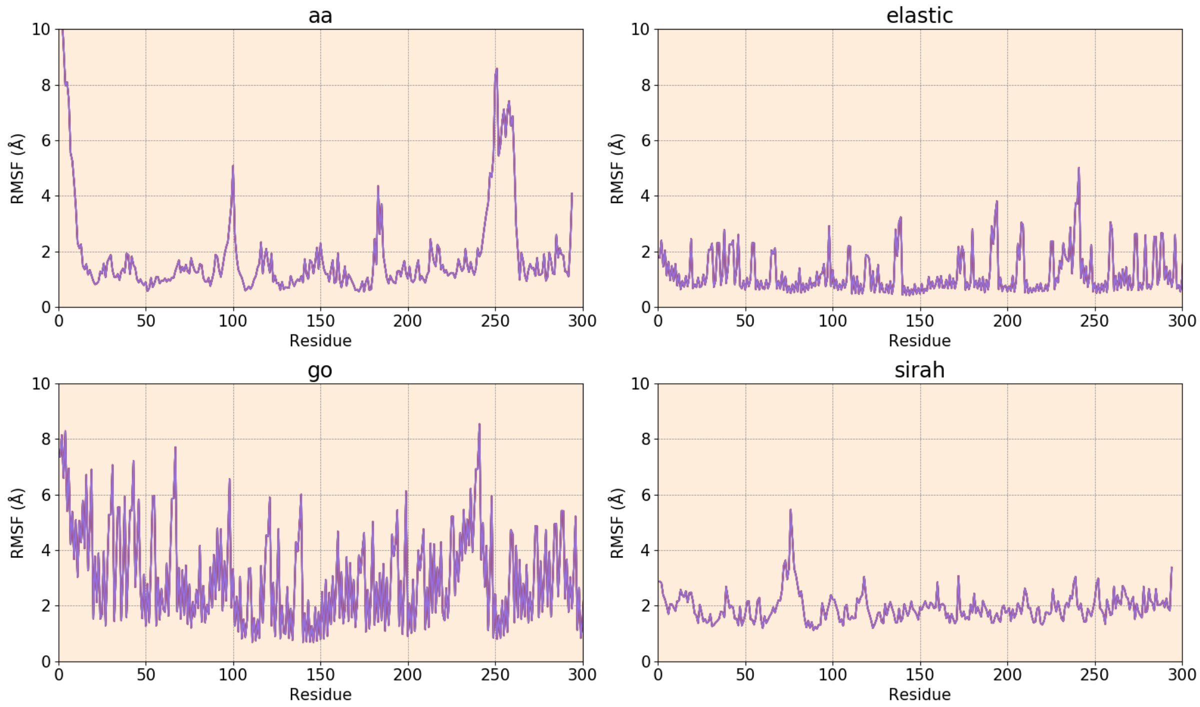


**Figure S31. RMSF of EC2 class enzyme 1q20 using all the methods – AA, Gō, Elastic, and SIRAH.**

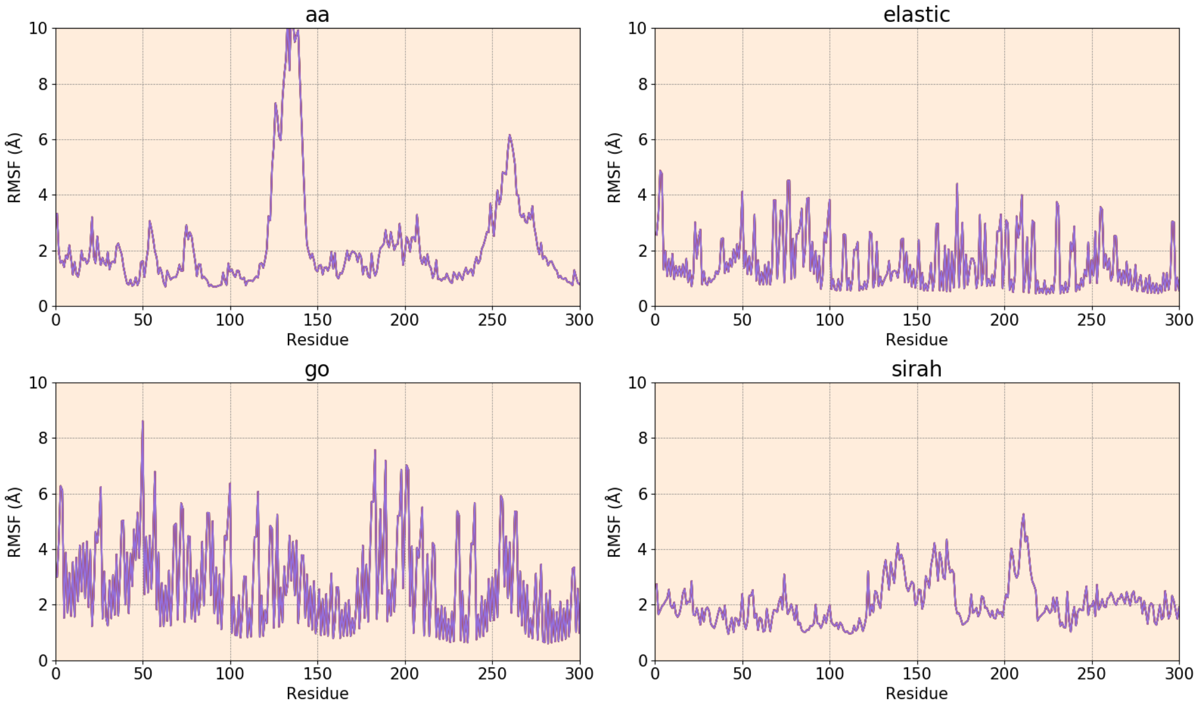


**Figure S32. RMSF of EC3 class enzyme 2oup using all the methods – AA, Gō, Elastic, and SIRAH.**

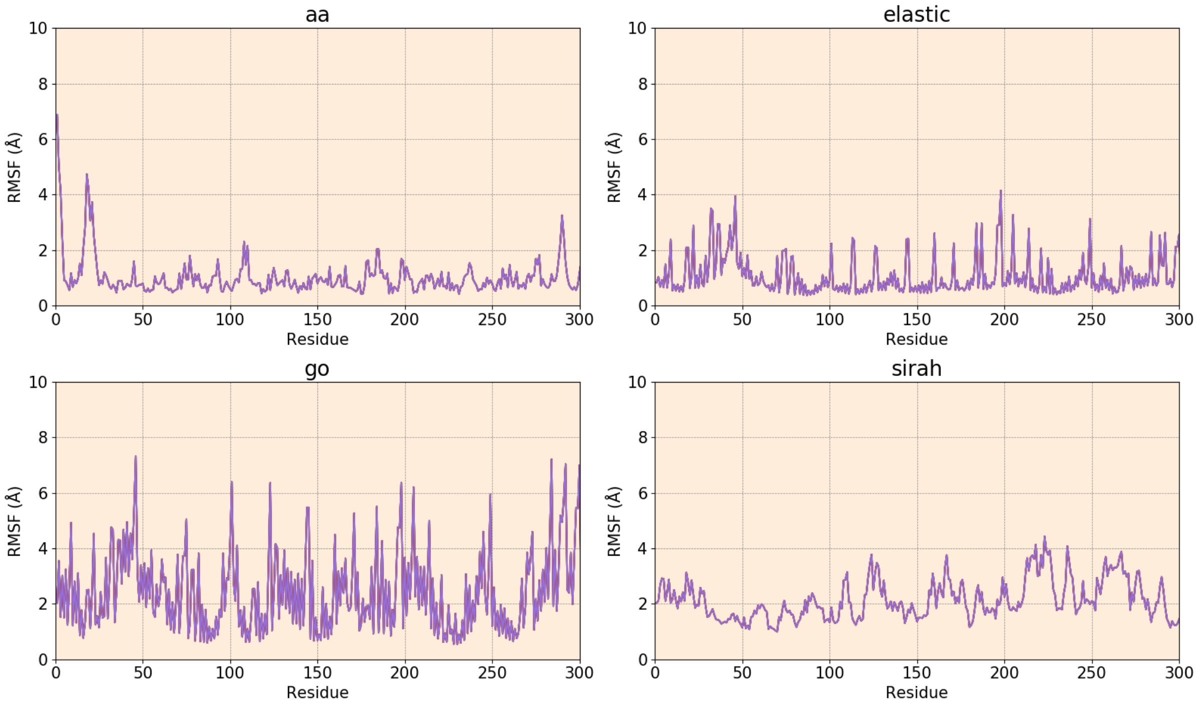


**Figure S33. RMSF of EC3 class enzyme 1dim using all the methods – AA, Gō, Elastic, and SIRAH.**

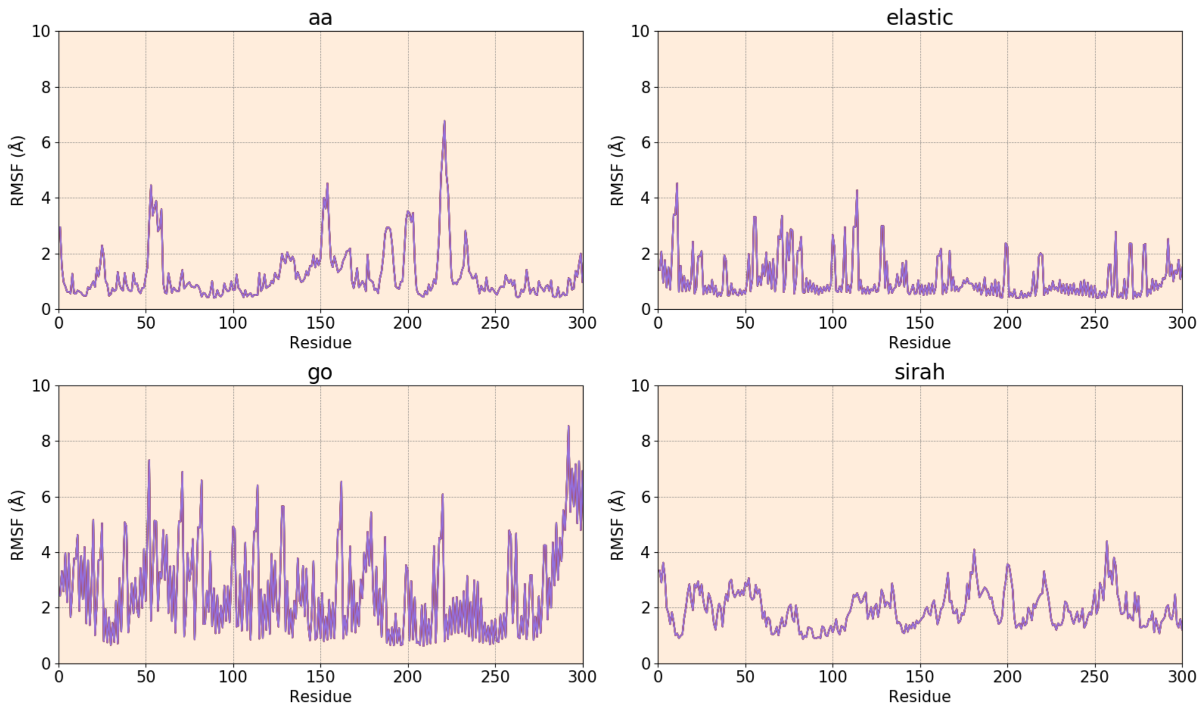


**Figure S34. RMSF of EC3 class enzyme 1cvl using all the methods – AA, Gō, Elastic, and SIRAH.**

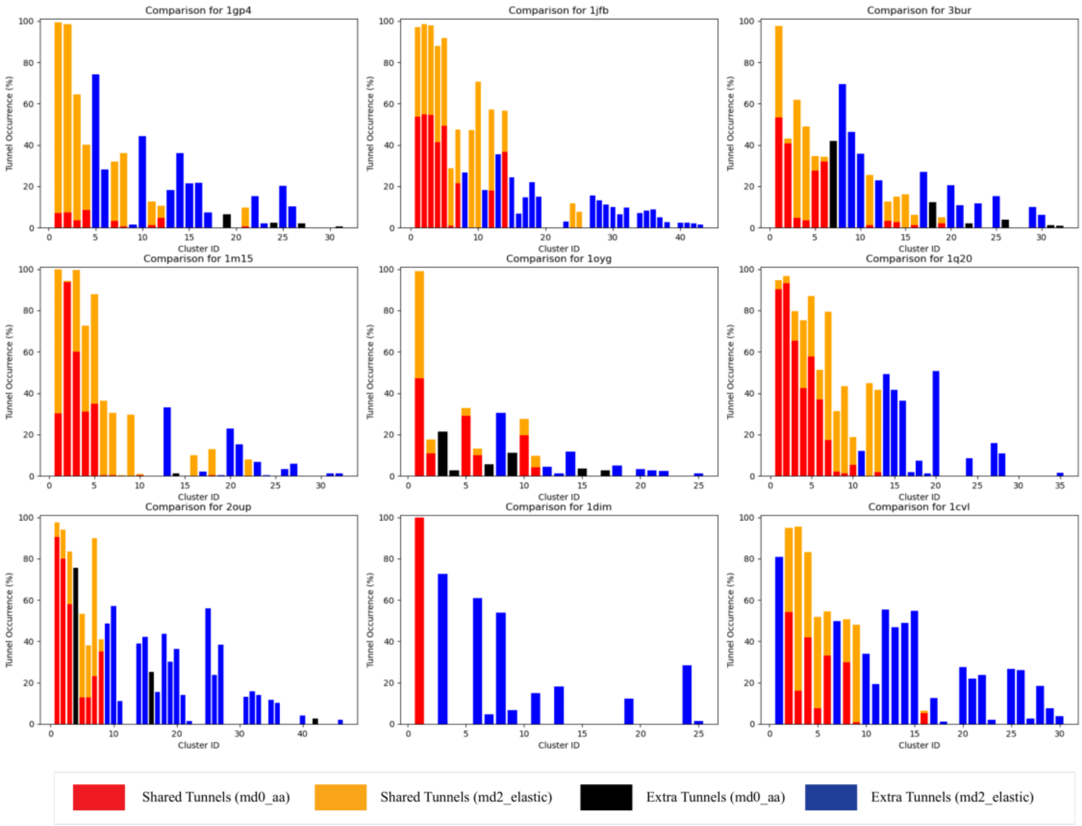


**Figure S35. Comparison of tunnel occurrences of equivalent tunnel clusters identified by Elastic and AA methods.** Note that the cluster ID is assigned based on the overall tunnel priority, i.e., tunnels from cluster 1 will in general be shorter, wider and have higher occurrence then tunnels from clusters with higher IDs.

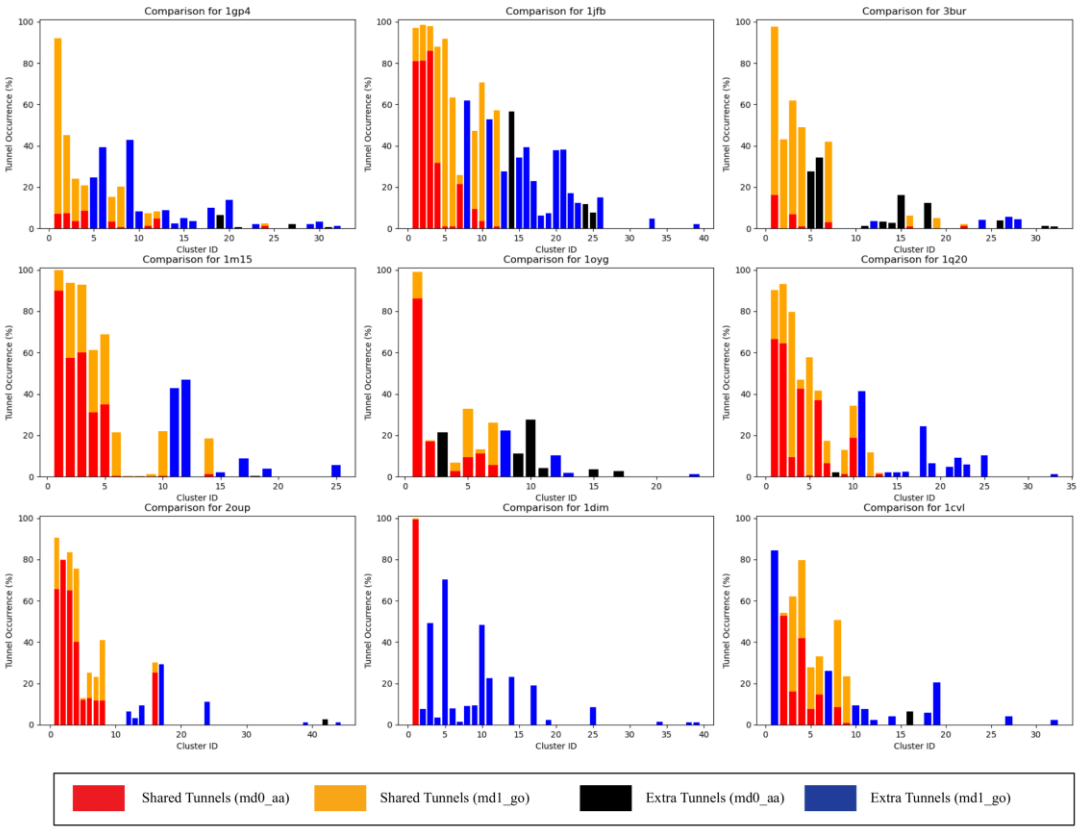


**Figure S36. Comparison of tunnel occurrences of equivalent tunnel clusters identified by Gō and AA methods.** Note that the cluster ID is assigned based on the overall tunnel priority, i.e., tunnels from cluster 1 will in general be shorter, wider and have higher occurrence then tunnels from clusters with higher IDs.

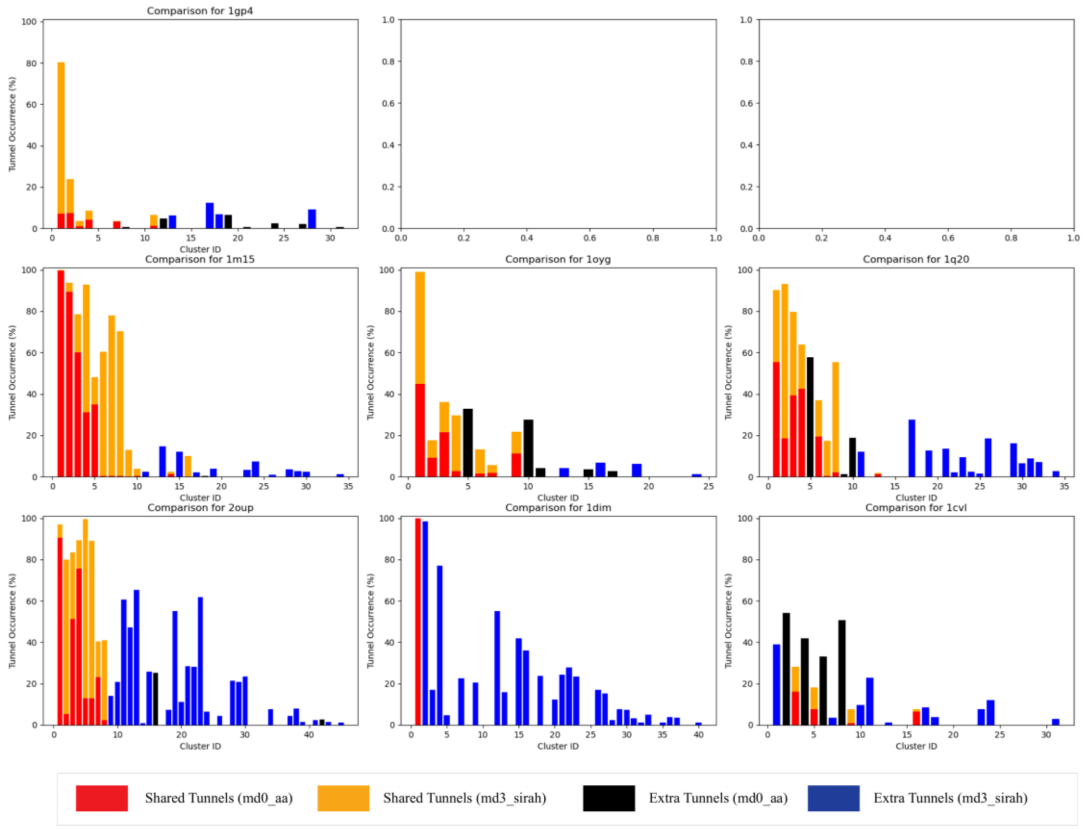


**Figure S37. Comparison of tunnel occurrences of equivalent tunnel clusters identified by SIRAH and AA methods.** Note that the cluster ID is assigned based on the overall tunnel priority, i.e., tunnels from cluster 1 will in general be shorter, wider and have higher occurrence then tunnels from clusters with higher IDs. Data from SIRAH simulation of 1fjb and 3bur enzymes are not available due to the lack of parameters for the cofactors.

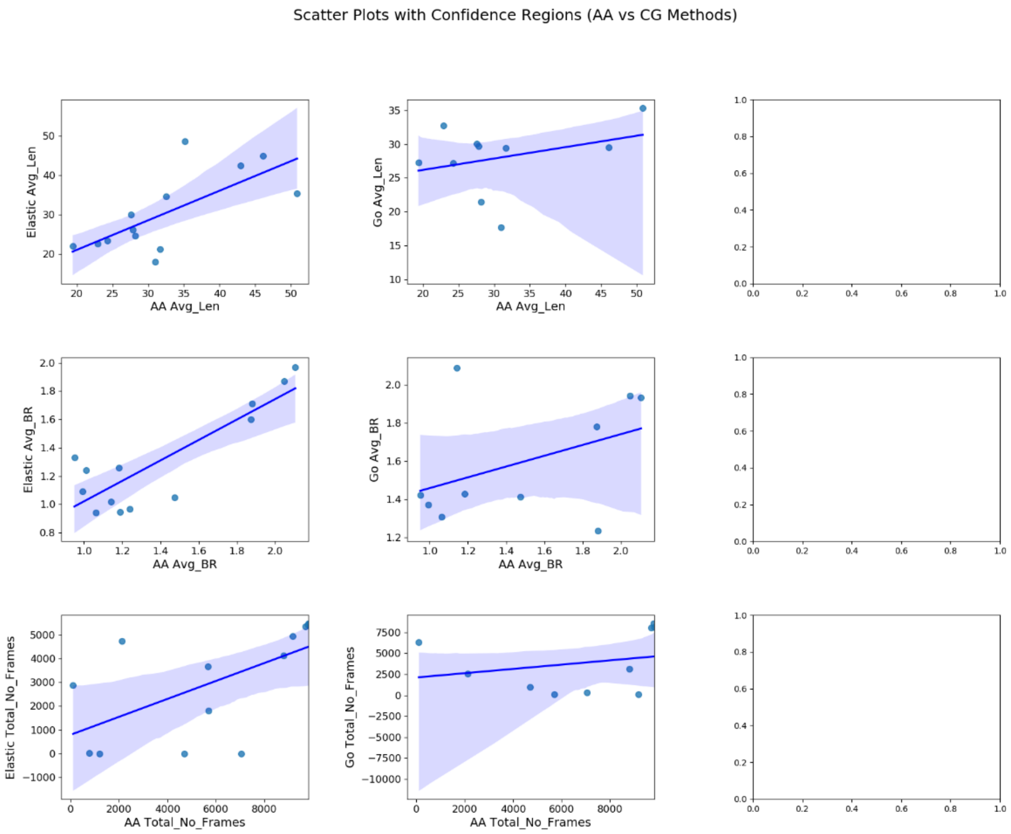


**Figure S38. Scatter plot between AA and CG methods in 1jfb from EC1.** Each point shows how properties of tunnels from AA method relates to the same property of equivalent tunnel found in a CG method (from left to right: Elastic, Gō, and SIRAH). The blue line in each plot is a linear regression fit to the data points, representing the overall trend between the two methods. The shaded region around the regression line represents the 95% confidence interval for the regression fit. Data from SIRAH simulation is not available due to the lack of parameters for the cofactor.

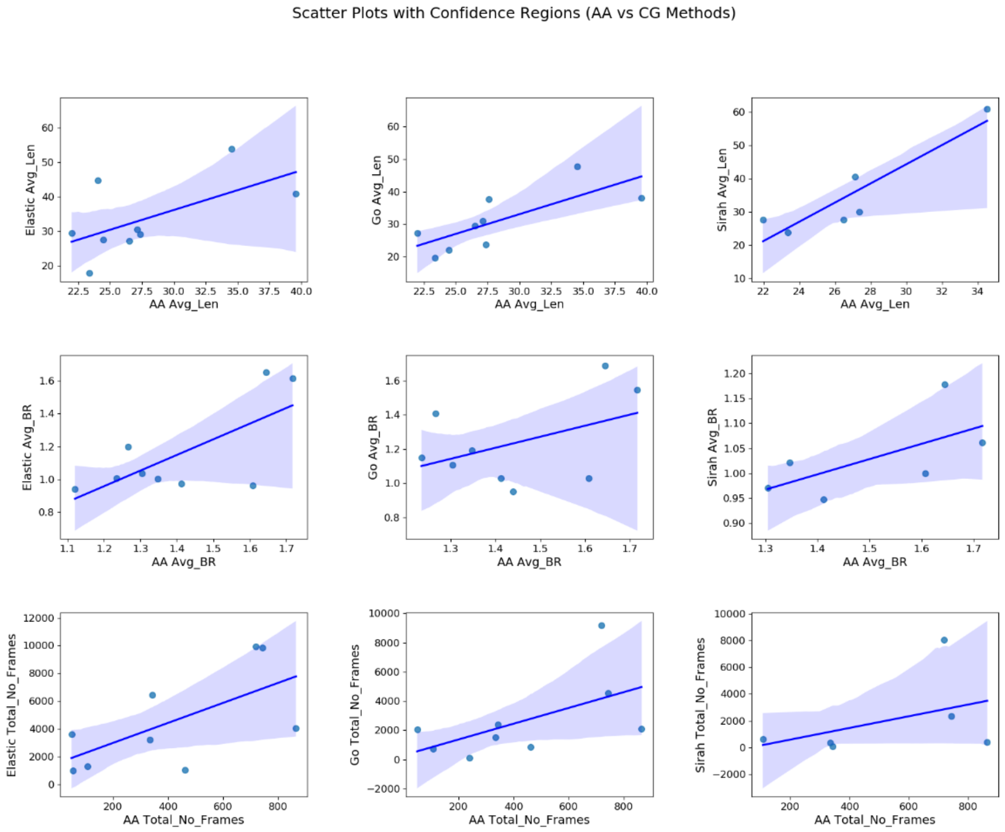


**Figure S39. Scatter plot between AA and CG methods in 1gp4 from EC1.** Each point shows how properties of tunnels from AA method relates to the same property of equivalent tunnel found in a CG method (from left to right: Elastic, Gō, and SIRAH). The blue line in each plot is a linear regression fit to the data points, representing the overall trend between the two methods. The shaded region around the regression line represents the 95% confidence interval for the regression fit.

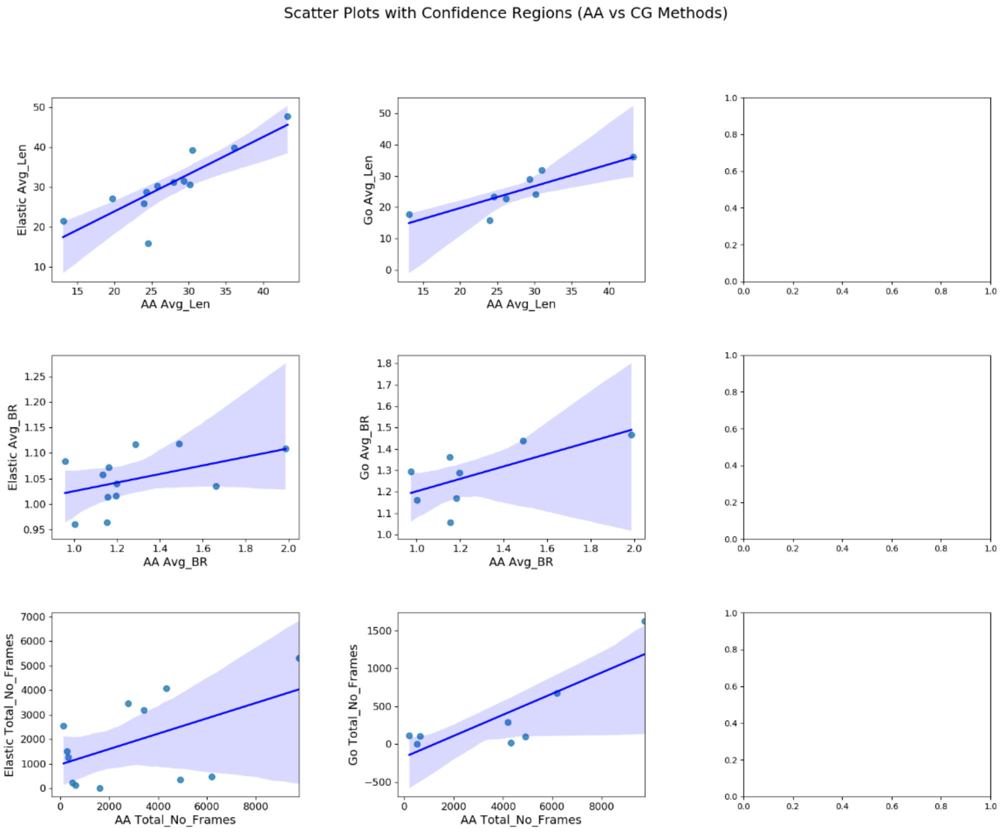


**Figure S40. Scatter plot between AA and CG methods in 3bur from EC1.** Each point shows how properties of tunnels from AA method relates to the same property of equivalent tunnel found in a CG method (from left to right: Elastic, Gō, and SIRAH). The blue line in each plot is a linear regression fit to the data points, representing the overall trend between the two methods. The shaded region around the regression line represents the 95% confidence interval for the regression fit. Data from SIRAH simulation is not available due to the lack of parameters for the cofactor.

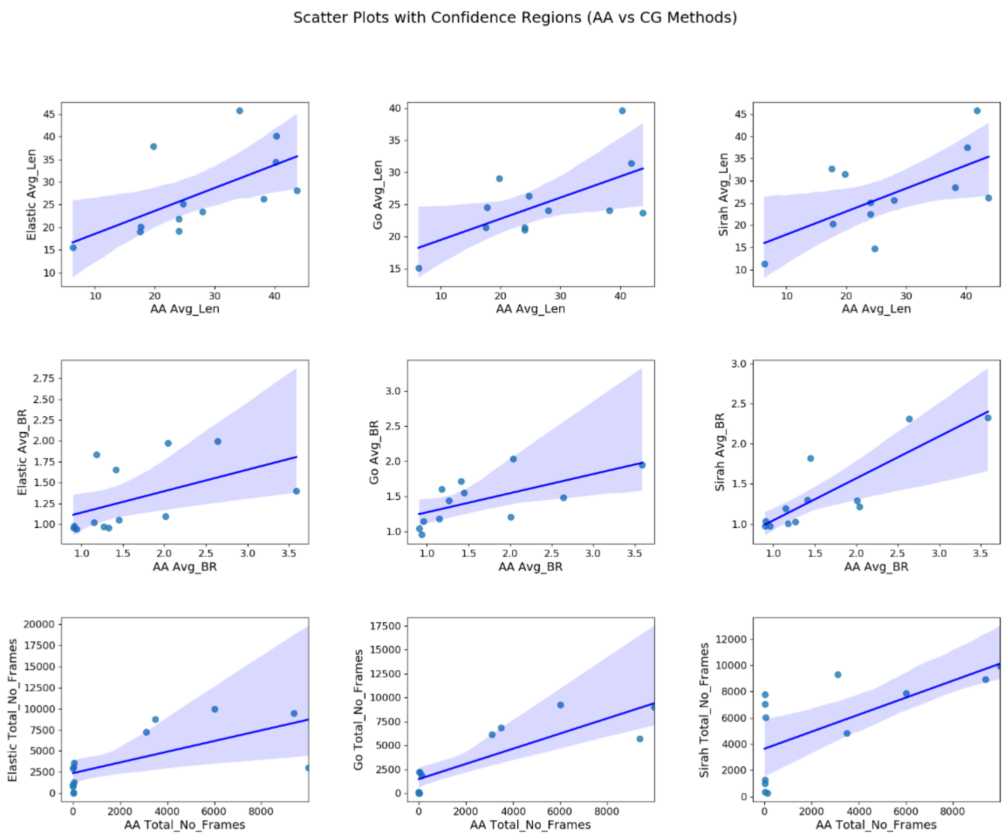


**Figure S41. Scatter plot between AA and CG methods in 1m15 from EC2.** Each point shows how properties of tunnels from AA method relates to the same property of equivalent tunnel found in a CG method (from left to right: Elastic, Gō, and SIRAH). The blue line in each plot is a linear regression fit to the data points, representing the overall trend between the two methods. The shaded region around the regression line represents the 95% confidence interval for the regression fit.

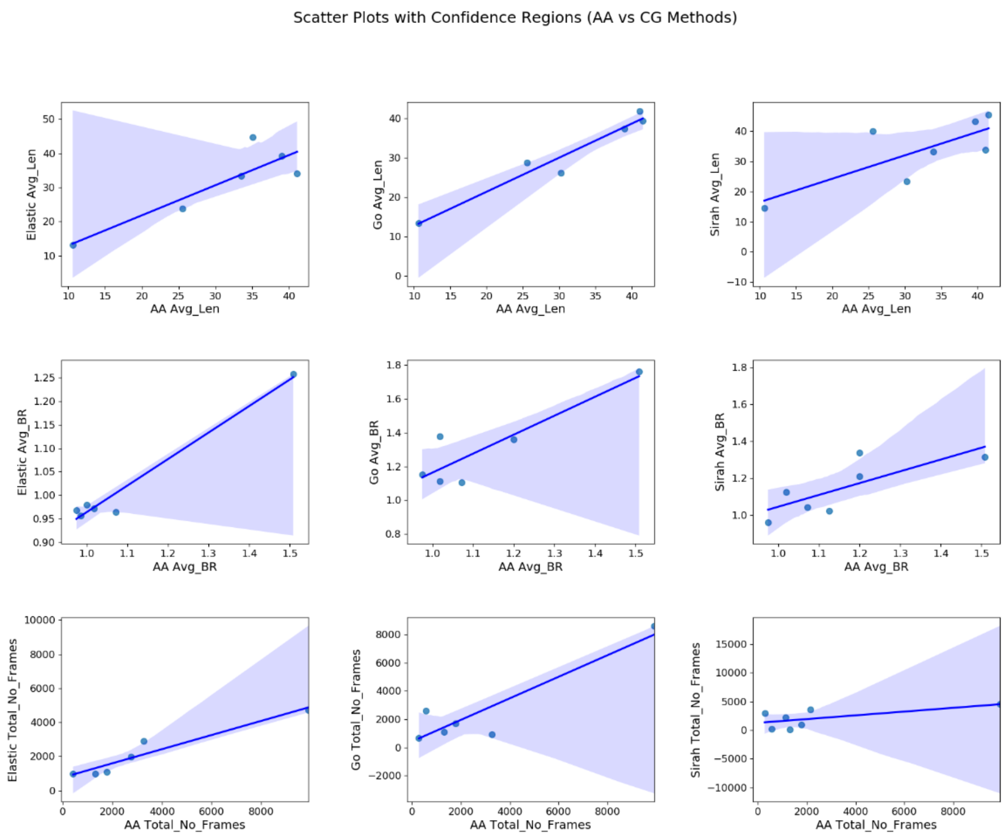


**Figure S42. Scatter plot between AA and CG methods in 1oyg from EC2.** Each point shows how properties of tunnels from AA method relates to the same property of equivalent tunnel found in a CG method (from left to right: Elastic, Gō, and SIRAH). The blue line in each plot is a linear regression fit to the data points, representing the overall trend between the two methods. The shaded region around the regression line represents the 95% confidence interval for the regression fit.

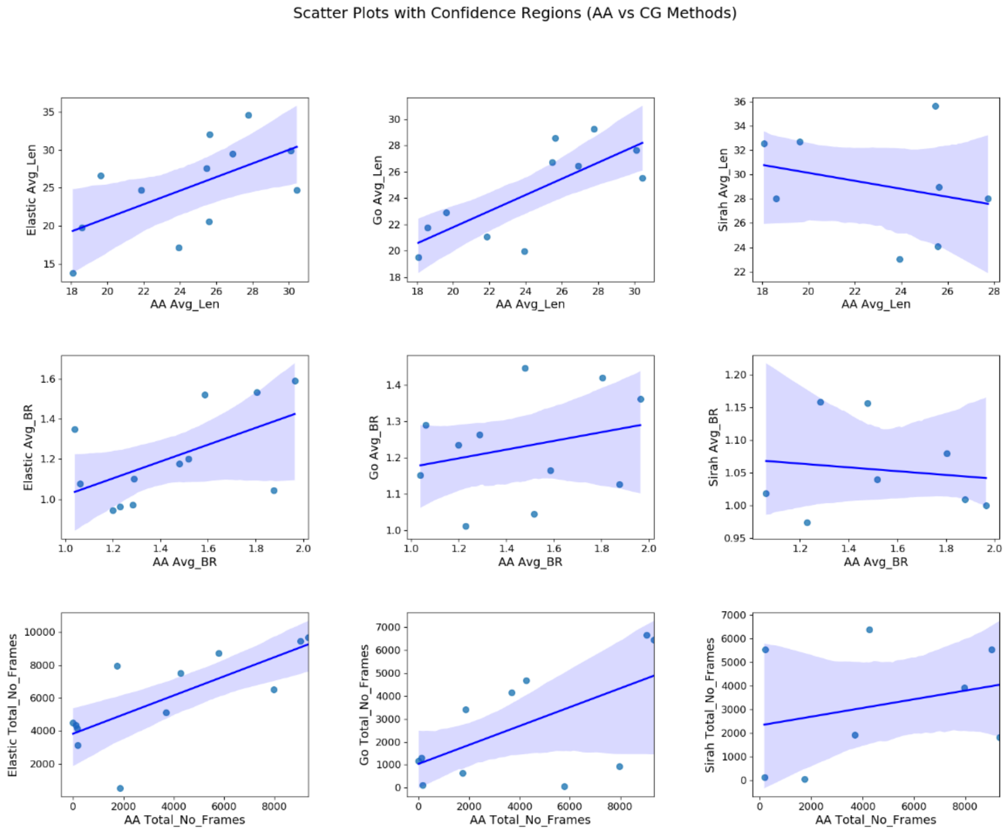


**Figure S43. Scatter plot between AA and CG methods in 1q20 from EC2.** Each point shows how properties of tunnels from AA method relates to the same property of equivalent tunnel found in a CG method (from left to right: Elastic, Gō, and SIRAH). The blue line in each plot is a linear regression fit to the data points, representing the overall trend between the two methods. The shaded region around the regression line represents the 95% confidence interval for the regression fit.

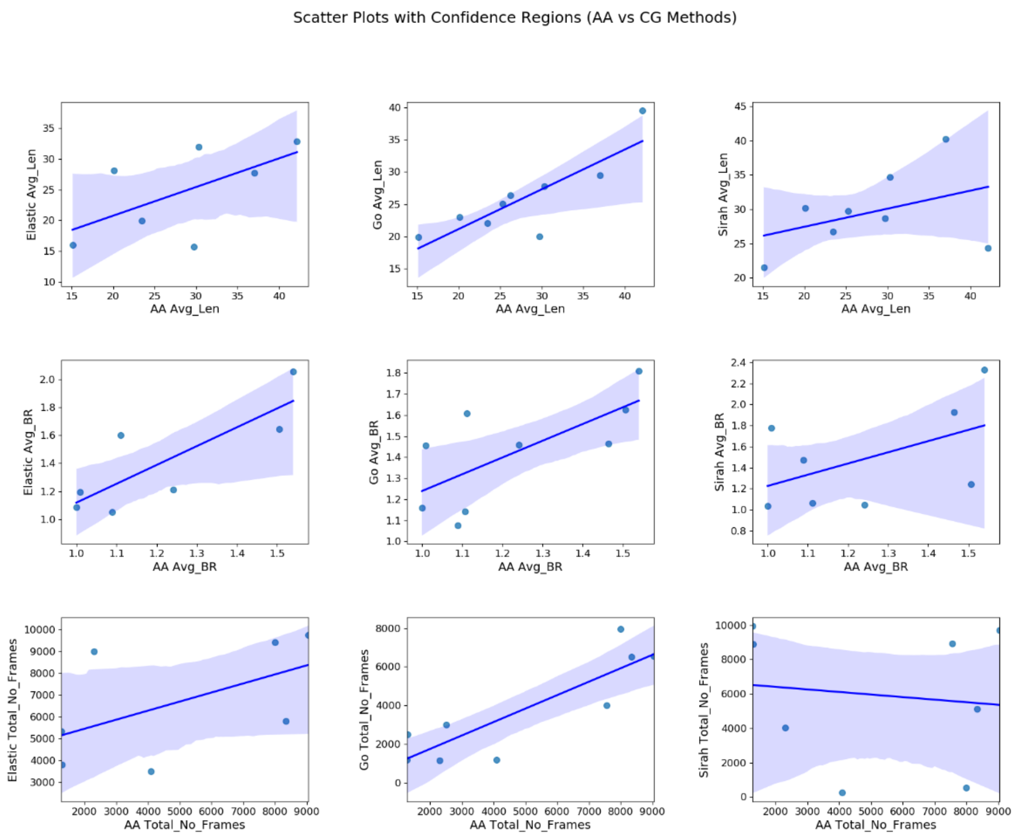


**Figure S44. Scatter plot between AA and CG methods in 2oup from EC3.** Each point shows how properties of tunnels from AA method relates to the same property of equivalent tunnel found in a CG method (from left to right: Elastic, Gō, and SIRAH). The blue line in each plot is a linear regression fit to the data points, representing the overall trend between the two methods. The shaded region around the regression line represents the 95% confidence interval for the regression fit.

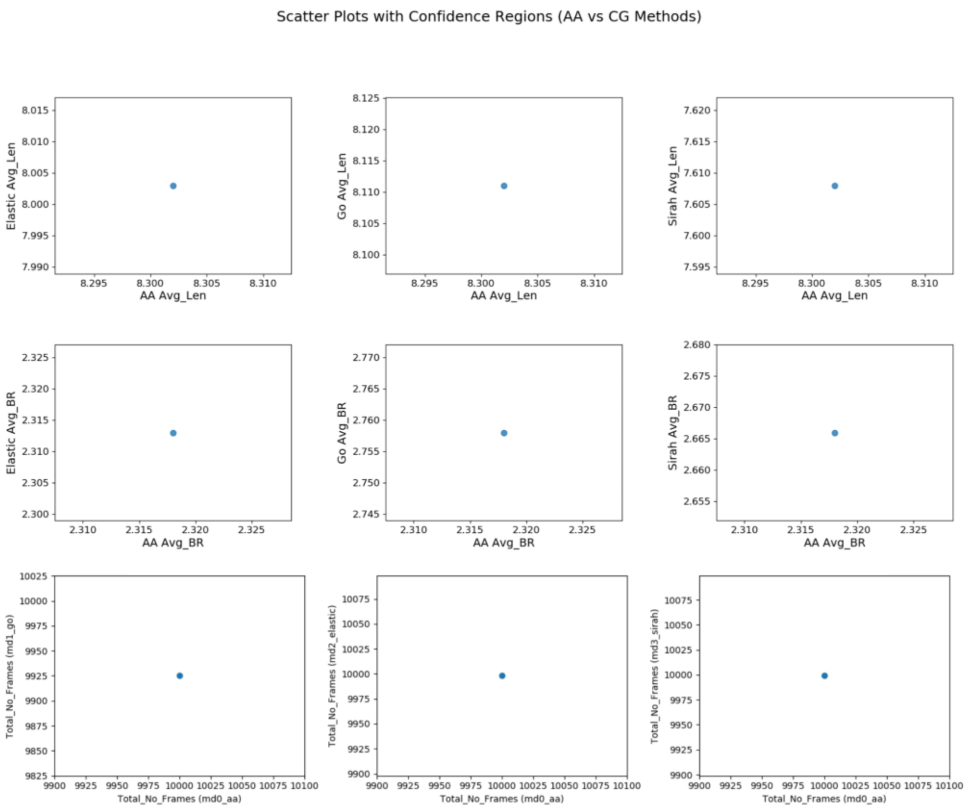


**Figure S45. Scatter plot between AA and CG methods in 1dim from EC3.** Each point shows how properties of tunnels from AA method relates to the same property of equivalent tunnel found in a CG method (from left to right: Elastic, Gō, and SIRAH). Here only 1 tunnel cluster was observed in AA cMD, disallowing correlation analysis for the protein.

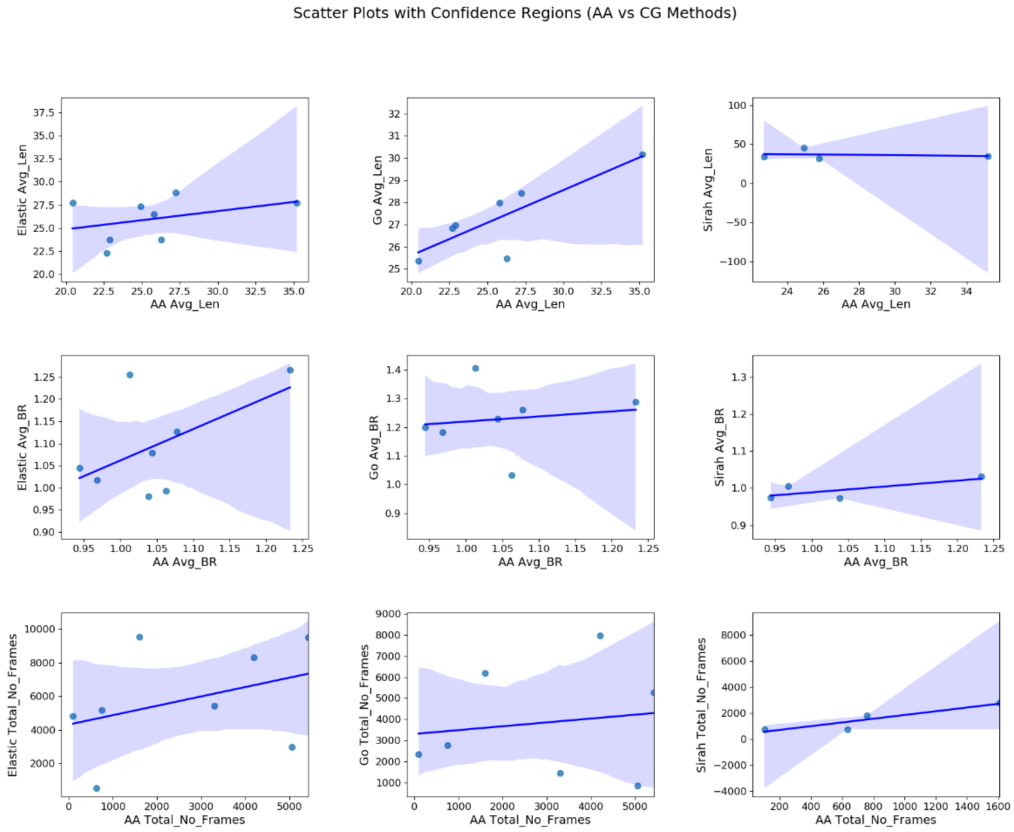


**Figure S46. Scatter plot between AA and CG methods in 1cvl from EC3.** Each point shows how properties of tunnels from AA method relates to the same property of equivalent tunnel found in a CG method (from left to right: Elastic, Gō, and SIRAH). The blue line in each plot is a linear regression fit to the data points, representing the overall trend between the two methods. The shaded region around the regression line represents the 95% confidence interval for the regression fit.

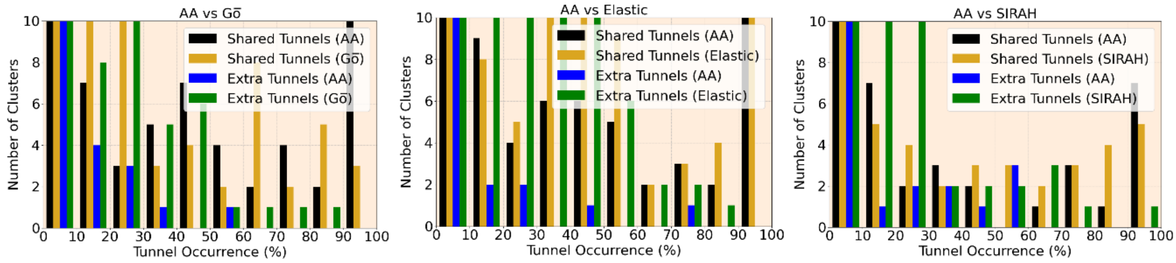


**Figure S47. Zoomed in distribution of tunnel occurrence of the equivalent tunnels shared between AA and CG methods as well as extra tunnels identified exclusively by AA or CG method.** Please note that this plot is zoomed in on extra tunnels. The number of clusters at some of the occurrence bins can extend over 50. See Figure 8A in the main text for the full view.

**Table S5. P-values from the two-sided test of the null hypothesis that the compared distributions are uncorrelated for dataset of EC1 class enzymes.**

| **Tunnel Occurrence** | | | | |
|---|---|---|---|---|
| | AA | Elastic | Gō | SIRAH |
| AA | 0.0000 | 0.2501 | 0.1901 | 0.4027 |
| Elastic | 0.2501 | 0.0000 | 0.0004 | 0.1229 |
| Gō | 0.1901 | 0.0004 | 0.0000 | 0.0017 |
| SIRAH | 0.4027 | 0.1229 | 0.0017 | 0.0000 |
| **Bottleneck Radius** | | | | |
| | AA | Elastic | Gō | SIRAH |
| AA | 0.0000 | 0.0000 | 0.0276 | 0.1746 |
| Elastic | 0.0000 | 0.0000 | 0.0009 | 0.0299 |
| Gō | 0.0276 | 0.0009 | 0.0000 | 0.0085 |
| SIRAH | 0.1746 | 0.0299 | 0.0085 | 0.0000 |
| **Length** | | | | |
| | AA | Elastic | Gō | SIRAH |
| AA | 0.0000 | 0.0000 | 0.0043 | 0.0117 |
| Elastic | 0.0000 | 0.0000 | 0.0000 | 0.0042 |
| Gō | 0.0043 | 0.0000 | 0.0000 | 0.0042 |
| SIRAH | 0.0117 | 0.0042 | 0.0042 | 0.0000 |

**Table S6. P-values from the two-sided test of the null hypothesis that the compared distributions are uncorrelated for dataset of EC2 class enzymes.**

| **Tunnel Occurrence** | | | | |
|---|---|---|---|---|
| | AA | Elastic | Gō | SIRAH |
| AA | 0.0000 | 0.0001 | 0.0000 | 0.0196 |
| Elastic | 0.0001 | 0.0000 | 0.0082 | 0.1597 |
| Gō | 0.0000 | 0.0082 | 0.0000 | 0.0098 |
| SIRAH | 0.0196 | 0.1597 | 0.0098 | 0.0000 |
| **Bottleneck Radius** | | | | |
| | AA | Elastic | Gō | SIRAH |
| AA | 0.0000 | 0.0012 | 0.0010 | 0.0000 |
| Elastic | 0.0012 | 0.0000 | 0.0004 | 0.0756 |
| Gō | 0.0010 | 0.0004 | 0.0000 | 0.0106 |
| SIRAH | 0.0000 | 0.0756 | 0.0106 | 0.0000 |
| **Length** | | | | |
| | AA | Elastic | Gō | SIRAH |
| AA | 0.0000 | 0.0000 | 0.0000 | 0.0005 |
| Elastic | 0.0000 | 0.0000 | 0.0000 | 0.0162 |
| Gō | 0.0000 | 0.0000 | 0.0000 | 0.0005 |
| SIRAH | 0.0005 | 0.0162 | 0.0005 | 0.0000 |

**Table S7. P-values from the two-sided test of the null hypothesis that the compared distributions are uncorrelated for dataset of EC3 class enzymes.**

| Tunnel Occurrence | | | | |
|---|---|---|---|---|
| | AA | Elastic | Gō | SIRAH |
| AA | 0.0000 | 0.0327 | 0.0033 | 0.1670 |
| Elastic | 0.0327 | 0.0000 | 0.0016 | 0.3023 |
| Gō | 0.0033 | 0.0016 | 0.0000 | 0.5438 |
| SIRAH | 0.1670 | 0.3023 | 0.5438 | 0.0000 |
| **Bottleneck Radius** | | | | |
| | AA | Elastic | Gō | SIRAH |
| AA | 0.0000 | 0.0000 | 0.0000 | 0.0012 |
| Elastic | 0.0000 | 0.0000 | 0.0000 | 0.0013 |
| Gō | 0.0000 | 0.0000 | 0.0000 | 0.0034 |
| SIRAH | 0.0012 | 0.0013 | 0.0034 | 0.0000 |
| **Length** | | | | |
| | AA | Elastic | Gō | SIRAH |
| AA | 0.0000 | 0.0028 | 0.0000 | 0.0564 |
| Elastic | 0.0028 | 0.0000 | 0.0000 | 0.0141 |
| Gō | 0.0000 | 0.0000 | 0.0000 | 0.0221 |
| SIRAH | 0.0564 | 0.0141 | 0.0221 | 0.0000 |

**Table S8. P-values from the two-sided test of the null hypothesis that the compared distributions are uncorrelated for combined dataset of EC1-EC3 classess of enzymes.**

| Tunnel Occurrence | | | | |
|---|---|---|---|---|
| | AA | Elastic | Gō | SIRAH |
| AA | 0.0000 | 0.0000 | 0.0000 | 0.0014 |
| Elastic | 0.0000 | 0.0000 | 0.0000 | 0.0319 |
| Gō | 0.0000 | 0.0000 | 0.0000 | 0.0017 |
| SIRAH | 0.0014 | 0.0319 | 0.0017 | 0.0000 |
| **Bottleneck Radius** | | | | |
| | AA | Elastic | Gō | SIRAH |
| AA | 0.0000 | 0.0000 | 0.0000 | 0.0000 |
| Elastic | 0.0000 | 0.0000 | 0.0000 | 0.0001 |
| Gō | 0.0000 | 0.0000 | 0.0000 | 0.0000 |
| SIRAH | 0.0000 | 0.0001 | 0.0000 | 0.0000 |
| **Length** | | | | |
| | AA | Elastic | Gō | SIRAH |
| AA | 0.0000 | 0.0000 | 0.0000 | 0.0014 |
| Elastic | 0.0000 | 0.0000 | 0.0000 | 0.0319 |
| Gō | 0.0000 | 0.0000 | 0.0000 | 0.0017 |
| SIRAH | 0.0014 | 0.0319 | 0.0017 | 0.0000 |
